# Mapping the Cellular Origin and Early Evolution of Leukemia in Down Syndrome

**DOI:** 10.1101/2020.11.29.402800

**Authors:** Elvin Wagenblast, Joana Araújo, Olga I. Gan, Sarah K. Cutting, Alex Murison, Gabriela Krivdova, Maria Azkanaz, Jessica L. McLeod, Sabrina A. Smith, Sajid A Marhon, Martino Gabra, Michelle Chan-Seng-Yue, Laura Garcia-Prat, Leonardo Salmena, Daniel D De Carvalho, Karen Chong, Maian Roifman, Patrick Shannon, Jean C Y Wang, Johann K. Hitzler, David Chitayat, John E. Dick, Eric R. Lechman

## Abstract

Children with Down syndrome have a 150-fold increased risk of developing myeloid leukemia, but the mechanism of predisposition is unclear. As Down syndrome leukemogenesis initiates during fetal development, we characterized the cellular context of preleukemic initiation and leukemic progression using gene editing in human disomic and trisomic fetal liver hematopoietic cells and xenotransplantation. *GATA1* mutations caused transient preleukemia only when introduced into trisomy 21 long-term hematopoietic stem cells, where a subset of chromosome 21 miRNAs triggers predisposition to preleukemia. By contrast, progression to leukemia was independent of trisomy 21 and originated in various stem and progenitor cells through additional mutations in cohesin genes. CD117+/KIT cells mediated the propagation of preleukemia and leukemia, and functional KIT inhibition targeted preleukemic stem cells, blocking progression to leukemia.

## Introduction

Children with Down syndrome (trisomy 21, T21) have a 150-fold increased risk of developing acute myeloid leukemia, termed myeloid leukemia associated with Down syndrome (ML-DS), in the first 5 years of life (*1*). However, the mechanism by which an extra copy of chromosome 21 predisposes and cooperates with genetic events in Down syndrome leukemogenesis is not known. In pediatric leukemia, the initiating genetic events occur before birth and generate preleukemic cells (*2*), which are the evolutionary ancestors of leukemia that arises after birth. However, characterizing human fetal preleukemia is challenging due to our inability to directly access it, rendering the identity of the cell of origin and the steps of leukemia evolution largely unknown. Up to 30% of newborns with Down syndrome exhibit transient abnormal myelopoiesis (TAM), a preleukemia characterized by a clonal proliferation of immature myeloid cells (mostly megakaryoblasts) carrying somatic mutations in the erythroid–megakaryocyte transcription factor *GATA1* (**fig. S1A**) (*3-5*). Mutations in *GATA1* occur *in utero* after 25 weeks of gestation (*6*) and lead to the expression of a truncated isoform (GATA1-short, GATA1s). The preleukemia resolves spontaneously in the majority of newborns, however in 20% of cases ML-DS evolves within 4 years from the GATA1s-mutated preleukemic clone by acquisition of additional mutations, predominantly in genes of the cohesin complex or CTCF (*7-9*). Comprehensive sequencing studies have shown that mutations in the cohesin subunit *STAG2* are most frequently implicated in ML-DS development (*10, 11*). Based on these observations, it is hypothesized that evolution of Down syndrome leukemia requires three distinct genetic events: T21, GATA1s and additional mutations such as *STAG2*. However, the identity of the human hematopoietic cell type that acquires GATA1s and originates preleukemia and the cell type in which subsequent mutations accumulate to generate leukemia are unknown. Furthermore, the cellular origin of preleukemic mutations in pediatric leukemias in general is currently unclear and Down syndrome leukemogenesis offers a novel disease setting to uncover generalized principles regarding this phenomenon.

Currently, there are no effective strategies to predict and potentially prevent the progression from preleukemia to ML-DS. Life-threatening symptoms associated with preleukemia are treated with cytarabine (*12, 13*), however this treatment does not prevent subsequent development of leukemia. Prevention of ML-DS would be associated with unequivocal advantages (*14*). Children with Down syndrome could be spared cytotoxic chemotherapy, which causes more adverse effects and complications in this group (*8*). Additionally, despite the generally favorable response of ML-DS to standard chemotherapy, outcomes are dismal for those with refractory or relapsed disease, with an overall survival rate of less than 20% (*15-17*). Therefore, therapeutic targeting of preleukemic clones would represent a potential new strategy to prevent development of leukemia and as a result improve survival outcomes for Down syndrome newborns with preleukemia. Experimentally, the mechanistic study of T21 and Down syndrome preleukemia has been challenging, primarily due to the fetal origin of the disease and the overall lack of suitable *in vivo* models (*18-20*). In order to circumvent these limitations, we used disomic and trisomic hematopoietic cells that were isolated from primary human fetal livers, the major hematopoietic organ during prenatal development, to investigate the mechanisms underlying preleukemia and leukemia.

Here, we describe a model that faithfully recapitulates the full spectrum of pre-malignant and malignant stages of Down syndrome leukemia using CRISPR/Cas9 methodology optimized for single human hematopoietic stem cells (HSC) (*21*) in disomic and trisomic primary human fetal liver-derived HSCs and downstream progenitors (*22*). Using this tool, we delineate the genetic events and cellular contexts of Down syndrome leukemogenesis and provide proof of concept for targeting the preleukemic stage of the disease.

## Results

### GATA1s induces a megakaryocytic bias in hematopoietic stem and progenitor cells

To study the initiating events in Down syndrome preleukemia, we first sorted HSPCs from normal (disomic) and T21 fetal livers (N-FL and T21-FL) obtained at 16 to 19 weeks of gestation and carried out phenotypic analysis of the hematopoietic stem and progenitor cell (HSPC) hierarchy (**Fig. 1A and fig. S1B**). Trisomy 21 karyotype of sorted HSPCs was confirmed by droplet digital PCR with a set of probes against chromosome 21 (**fig. S1C**). Sanger sequencing of *GATA1* exon 2 in T21-FL-derived HSPCs did not detect any pre-existing GATA1 mutations (**fig. S1C**). The impact of T21 on the HSPC hierarchy in fetal liver revealed a 30% increase in the percentage of total long-term HSCs (LT-HSC) and a simultaneous decrease in short-term HSCs (ST-HSC) compared to N-FL. In addition, an expansion of myeloid-erythroid progenitors (MEP) was seen in T21-FL compared to N-FL, as previously reported (**fig. S1, D to E, and fig. S2A**) (*23*). Quantification of GATA1 and its isoforms revealed comparable expression in N-FL and T21-FL HSPC subpopulations with a gradual increase in expression upon differentiation commitment (**fig. S2B**).

**Fig. 1.**
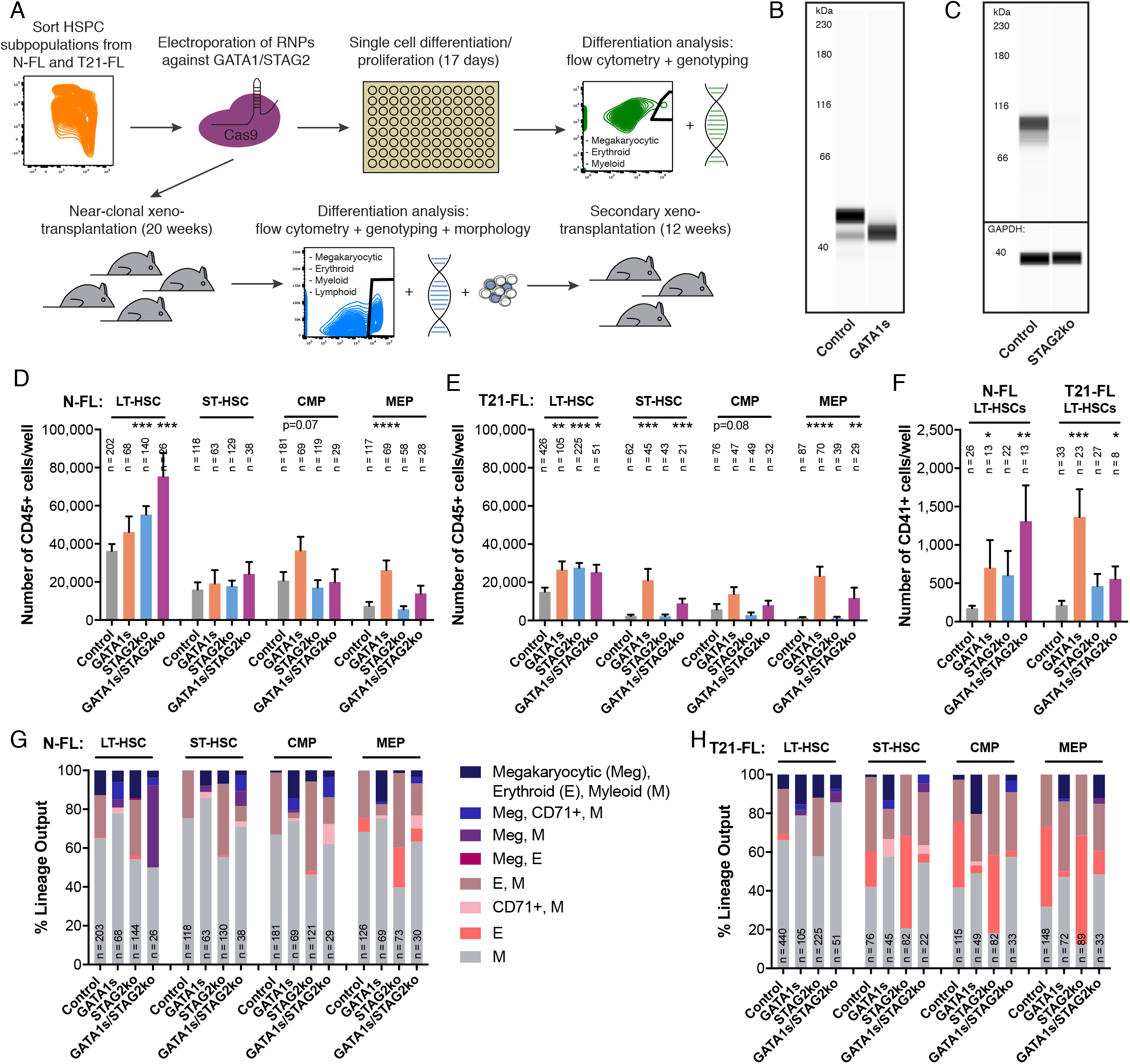
GATA1s induces a megakaryocytic bias in hematopoietic stem and progenitor cells. **(A)** Experimental overview of *in vitro* single cell differentiation/proliferation assay and near-clonal xenotransplantation. **(B)** Western blot assay of GATA1 in combined CMP and MEP cells, which were CRISPR/Cas9-edited with control and GATA1s gRNAs (n = 1 experiment). **(C)** Western blot assay of STAG2 in combined CMP and MEP cells, which were CRISPR/Cas9-edited with control and STAG2 knockout gRNAs (n = 1 experiment). **(D)** Proliferation capacity assessed by the overall number of CD45+ cells from *in vitro* single cell assay of individual CRISPR/Cas9-edited cells for N-FL. Numbers of single cell colonies with appropriate positive genotype are indicated for each condition (n = 2 experiments). **(E)** Proliferation capacity described in (D) for T21-FL (§§§§p for combined T21-FL versus N-FL, n = 2-3 experiments). **(F)** Proliferation capacity assessed for number of CD41+ cells from N-FL and T21-FL LT-HSCs from single cell assays described in (D) and (E). **(G)** Lineage output from *in vitro* single cell assay described in (D) (*p for Meg in GATA1s versus control, p=0.12 for Meg in GATA1s/STAG2ko versus control and **p for E in GATA1s versus control among all cell types, n = 2 experiments). **(H)** Lineage output from *in vitro* single cell assay as described in (G) for T21-FL (***p for Meg in GATA1s versus control, **p for Meg in GATA1s/STAG2ko versus control and p=0.16 for E in GATA1s versus control among all cell types, n = 2-3 experiments). Unpaired t test: *p < 0.05; **p < 0.01; ***p < 0.001; ****/§§§§p<0.0001; error bars represent standard error of the mean.

To examine the role of GATA1s and *STAG2* in leukemogenesis both individually or in combination, we performed CRISPR/Cas9 editing of sorted HSPC subpopulations to express the short isoform of *GATA1* (GATA1s) under its endogenous promoter and/or to knock-out the cohesin subunit STAG2 (STAG2ko). Using our optimized methodology (*21*), editing efficiency exceeded 75% (**fig. S2C**). Karyotyping analysis of N-FL HSPCs revealed no structural abnormalities after CRISPR/Cas9 editing (**fig. S2D**) and whole genome sequencing in N-FL LT-HSCs at 30x coverage revealed either very rare or no off-target indels at sites that were similar to the gRNA sequence (**table S1**). In the few cases where off-target indels were detected, the allelic depth was 6% or lower. Western blot assays of CRISPR/Cas9-edited common myeloid progenitors (CMP) and MEPs showed exclusive expression of GATA1s and undetectable protein levels of STAG2 (**Fig. 1, B and C**), confirming CRISPR/Cas9 editing of the respective genes at the protein level.

In order to elucidate the functional consequences of GATA1s and STAG2ko in different HSPC subpopulations, N-FL and T21-FL LT-HSCs, ST-HSCs, CMPs and MEPs that were CRISPR/Cas9-edited for control, GATA1s, STAG2ko or GATA1s/STAG2ko were placed into single cell *in vitro* differentiation and proliferation assays using erythro-myeloid-promoting media (*24*). The phenotype and genotype of all ~3,000 single cell-derived colonies used in our study was determined (**fig. S2, E and F**). Consistent with previous reports (*23*) colony-forming efficiency was higher in CRISPR/Cas9-edited T21-FL LT-HSCs compared to N-FL (**fig. S3, A and B**). The single cell CRISPR/Cas9 editing efficiency was above 80% for both control and STAG2ko colonies (**fig. S3, C and D**). For GATA1s, the CRISPR/Cas9 efficiency was ~40% since only colonies with confirmed complete excision of exon 2 were included in the analysis.

Proliferation measured by total CD45+ cell output was lower in T21-FL HSPC subpopulations compared to N-FL (**Fig. 1, D and E**). A sharp increase in cell numbers was observed in T21-FL GATA1s and GATA1s/STAG2ko subpopulations compared to control colonies. A similar trend was noticed in N-FL progenitor subpopulations but to a much lesser extent. The increase in proliferative capacity in GATA1s and GATA1s/STAG2ko LT-HSCs was accompanied by a significant increase in the production of CD41+ megakaryocytes (**Fig. 1F**). To investigate if the decreased proliferative capacity of T21-FL was related to a change in the number of cycling or quiescent cells, we performed cell cycle analysis. Interestingly, T21-FL HSPC subpopulations contained a lower percentage of cells in S-phase and a higher frequency of cells arrested in G0/G1-phase compared to N-FL (**fig. S3E**). No difference was observed in the ratio of quiescent G0 to G1 cells between N-FL and T21-FL (**fig. S3F**). Thus, despite an increased proportion of the LT-HSC compartment in T21-FL (**fig. S2A**) and higher colony forming capacity (**fig. S3, A and B**), T21-FL cells exhibit a significant proliferation restraint compared to their N-FL counterparts. However, the acquisition of GATA1 mutation drastically increases their proliferative capacity, providing a selection advantage for GATA1s T21-FL HSPCs.

Phenotypic analysis of colonies derived from single N-FL HSPCs revealed a drastic shift towards megakaryocytic differentiation and a concomitant decrease in erythroid lineage output in GATA1s and GATA1s/STAG2ko colonies compared to controls (**Fig. 1G**). A subset of these GATA1s and GATA1s/STAG2ko colonies expressed the early erythroid marker CD71 but not the mature erythroid marker GlyA, consistent with a block in erythroid differentiation (**fig. S3G**). Similar to N-FL, T21-FL GATA1s and GATA1s/STAG2ko HSPC subpopulations displayed a significant megakaryocytic bias and a decrease in erythroid output compared to controls (**Fig. 1H and fig. S3H**). In contrast, both N-FL and T21-FL STAG2ko cells exhibited an increase in erythroid output compared to controls. Collectively these *in vitro* results indicate that exclusive expression of GATA1s with or without STAG2ko resulted in increased megakaryocytic output in all HSPC subpopulations, with no major differences between N-FL and T21-FL.

### Trisomy 21 is required for preleukemia initiation but dispensable for leukemia development

To evaluate the functional effects of GATA1s and STAG2ko *in vivo*, we utilized LT-HSCs, as these are the only cells that have the ability to permanently repopulate the entire hematopoietic system following transplantation (*22*) and performed xenotransplantation assays using NSG and NSGW41 recipients. The repopulating cell frequency of N-FL LT-HSCs injected into mice evaluated at 20 weeks was approximately 1 in 300 (**fig. S3, I and J**). We therefore transplanted N-FL and T21-FL control, GATA1s, STAG2ko and GATA1s/STAG2ko LT-HSCs at cell doses of 300 to 400 into mice in order to obtain near-clonal grafts. After 20 weeks, human engraftment was analyzed in the bone marrow and extramedullary hematopoiesis was assessed in the spleen. Only mice bearing confirmed CRISPR/Cas9-edited grafts were used in the subsequent analysis (**fig. S4, A to C**). To evaluate the clonality of xenografts, cells harvested from bone marrow of transplanted mice were plated in methylcellulose colony assays. Sanger sequencing of CRISPR/Cas9-mediated indels in individual colonies showed clonal engraftment in ~60-100% of mice (**fig. S4, D and E**), thereby validating our *in vivo* experimental approach.

On average, the human CD45+ engraftment level in bone marrow was ~25% for mice transplanted with N-FL LT-HSCs and lower for T21-FL LT-HSCs, with the exception of mice transplanted with GATA1s/STAG2ko LT-HSCs, which displayed engraftment levels of ~30% (**Fig. 2, A and B**). Lineage marker analysis revealed increased myeloid and decreased lymphoid lineage cells in T21-FL control grafts compared to N-FL (**Fig. 2, C and D, and fig. S3, F to R**). Importantly, the proportion of human CD41+CD45– megakaryocytic lineage cells was at least 3-fold higher in GATA1s and GATA1s/STAG2ko grafts compared to control for both N-FL and T21-FL, consistent with the results observed in the *in vitro* single cell assays. Moreover, immunohistochemistry (IHC) staining for the megakaryocytic marker CD61 in bone sections of humeri revealed an increase in megakaryocytic cells in mice engrafted with GATA1s and GATA1s/STAG2ko cells from both N-FL and T21-FL (**Fig. 2E and fig. S5, A to C**). Engraftment patterns were similar in NSGW41 and NSG recipients of CRISPR/Cas9-edited NFL and T21-FL (**fig. S4, D to L**).

**Fig. 2.**
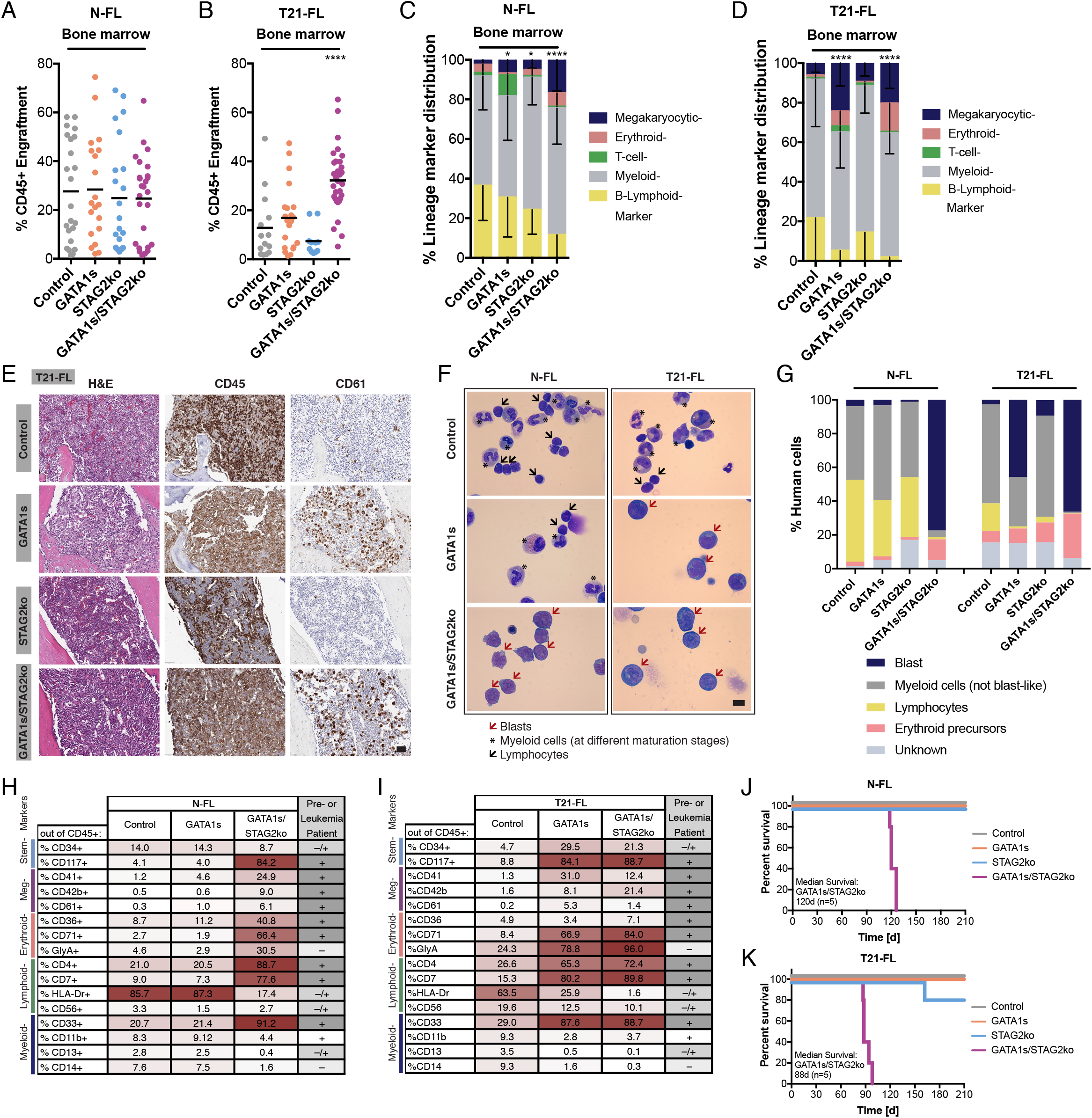
Trisomy 21 is required for preleukemia initiation but dispensable for leukemia development. **(A)** Engraftment levels of N-FL LT-HSC grafts in NSG mice. Engraftment was assessed based on human CD45+ expression in bone marrow (only mice with >1% of CD45+ cells in bone marrow and >90% CRISPR/Cas9 efficiency were taken for the analysis, n = 3 cohorts). **(B)** Engraftment levels as described in (A) for T21-FL (n = 4 cohorts). **(C)** Lineage marker distribution based on cell surface markers in N-FL grafts in NSG mice. Megakaryocytic cells were identified as CD41+CD45–, erythroid lineage as GlyA+CD45–, myeloid cells as CD33+CD45+, B-lymphoid cells as CD19+CD45+ and T-cells as CD3+CD45+. **(D)** Lineage marker distribution as described in (C) for T21-FL (§p for T21-FL control myeloid cells versus N-FL control myeloid cells and §p for T21-FL control lymphoid cells versus N-FL control lymphoid cells). **(E)** Hematoxylin and Eosin (H&E) and immunohistochemistry (IHC) stainings for human CD45 and human megakaryocytic marker CD61 in humeri of T21-FL grafts (scale 50μm). **(F)** Morphological analysis of human cells in primary xenografts of N-FL and T21-FL grafts. Human cells were cytospinned and stained with Giemsa (100x magnification, scale 10μm). **(G)** Quantification of cell morphology as seen in (F) (n = 400 cells per condition). **(H)** Percent expression of cell surface markers within the CD45+ blast population in N-FL grafts in NSG mice. Individual percentages are shaded based on a 2-color scale, normalized by the whole table and this also applies to (I) (data from pooled samples of multiple xenografts). **(I)** Percent expression of cell surface markers as described in (H) for T21-FL (data from pooled samples of multiple xenografts). **(J)** Survival curve of N-FL LT-HSC grafts in NSGW41 mice (n = 5 mice per condition). **(K)** Survival curve as described in (J) for T21-FL (n = 5 mice per condition). Unpaired t test: */§p < 0.05; **p < 0.01; ***p < 0.001; ****p < 0.001; error bars represent standard deviation.

To investigate whether the observed lineage shifts were associated with development of preleukemia or malignant transformation to full leukemia, we assessed xenografts for the presence of immature blast cells and found a dramatic increase in blasts to ~30-40% in T21 GATA1s, but surprisingly not in N-FL GATA1s xenografts (**Fig. 2, F and G, and fig. S5M**). Higher blast percentages of ~50-80% were observed in both N-FL and T21-FL GATA1s/STAG2ko grafts. We carried out a detailed flow cytometric analysis of lineage markers on large non-granulated cells in the blast gate (**fig. S6A**). Notably, N-FL and T21-FL control and N-FL GATA1s grafts had no enrichment of this gated population. The blast population of T21-FL GATA1s grafts expressed the primitive stem cell markers CD34 and CD117 (KIT), megakaryocytic marker CD41, erythroid markers CD71 and GlyA, myeloid marker CD33 and also aberrantly expressed lymphoid markers CD4 and CD7. This immunophenotype accurately recapitulates the clinical phenotype seen in patients with preleukemic TAM (**Fig. 2, H and I**) (*12, 25-27*). Interestingly, blasts in both N-FL and T21-FL GATA1s/STAG2ko grafts had immunophenotypes nearly identical to those of T21-FL GATA1s grafts, in keeping with the clinical observation that blasts from patients in the preleukemic and leukemic stages are often indistinguishable (*27, 28*). The blast immunophenotype of grafts generated in NSGW41 mice followed a comparable pattern (**fig. S6, B and C**).

We next assessed the survival of NSGW41 mice transplanted with 1,300 N-FL or T21-FL control, GATA1s, STAG2ko or GATA1s/STAG2ko LT-HSCs. No effect on overall survival was found in mice transplanted with control, GATA1s, or STAG2ko LT-HSCs from N-FL and T21-FL during the observation period of 210 days. In contrast, mice transplanted with either N-FL or T21-FL GATA1s/STAG2ko cells had a shorter median survival of 120 and 88 days, respectively (**Fig. 2, J and K**), highlighting an important difference between the preleukemic and leukemic disease in this model. Our findings unexpectedly demonstrate that T21 is necessary for preleukemia development driven by GATA1s but dispensable for leukemic progression upon acquisition of STAG2ko.

### CD117 marks preleukemia and leukemia initiating cells, which possess a more MEP-like chromatin accessibility landscape

To assess the self-renewal properties of T21 GATA1s-induced preleukemia and GATA1s/STAG2ko-induced leukemia, we carried out secondary xenotransplantation assays. As CD34 expression is absent in some Down syndrome leukemia cases (*27*), we sorted all primitive CD34+ and CD117+ cells from primary xenografts and transplanted them at defined doses into secondary NSGW41 recipients (**Fig. 3A**). We observed dramatic differences in self-renewal as measured by the ability of N-FL versus T21-FL GATA1s cells to propagate hematopoiesis in secondary recipients (**Fig. 3B**). Whereas secondary grafts originating from N-FL GATA1s cells were phenotypically similar to control grafts after 12 weeks, preleukemic T21-FL GATA1s secondary grafts contained characteristic blast populations equivalent to those seen in primary recipients (**fig. S6, D and E**), although with a lower preleukemia-initiating cell frequency of ~1/150,000. Interestingly, both N-FL and T21-FL STAG2ko grafts had higher initiating-cell frequencies compared to controls, albeit 8-fold less in T21-FL compared to N-FL (**Fig. 3B**), consistent with the previously reported increase in HSC self-renewal in a STAG2 knock-out mouse model (*29*). Both N-FL and T21-FL GATA1s/STAG2ko cells from primary grafts were able to generate secondary leukemic grafts containing characteristic blast populations (**fig. S6, D and E**), with initiating-cell frequencies of ~1/45,000 and ~1/90,000, respectively.

**Fig. 3.**
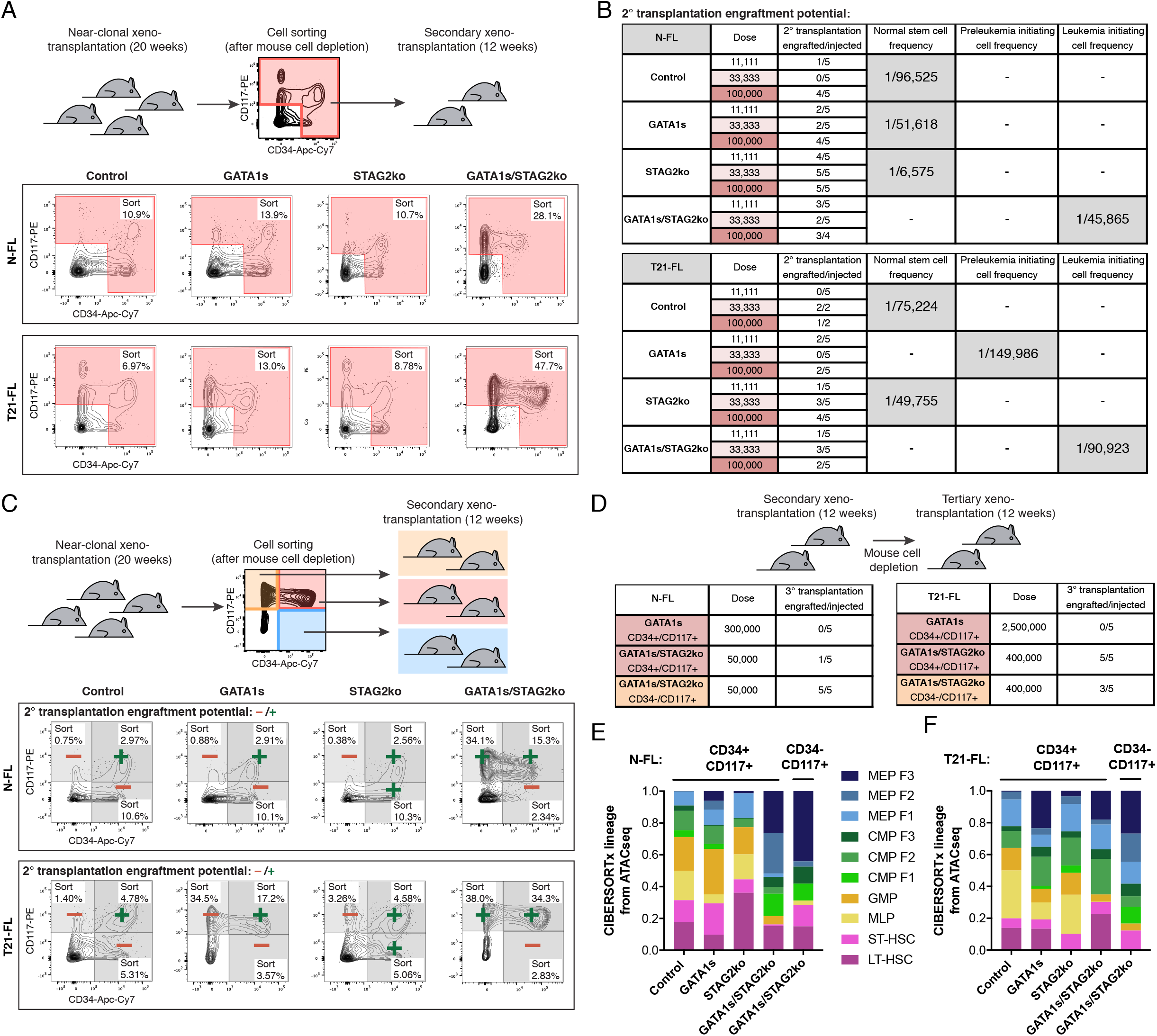
CD117 marks preleukemia and leukemia initiating cells. **(A)** Experimental overview of secondary xenotransplantation experiments. Flow cytometry plots of sorted human fractions from primary grafts are depicted. **(B)** Stem cell frequencies based on secondary xenotransplantations as described in (A). Limiting dilution analysis was used to assess normal, preleukemia and leukemia initiating cell frequencies (>0.1% CD45+ cells in bone marrow, n = 2-5 mice for each condition and dose, in total 113 mice). **(C)** Experimental overview of secondary xenotransplantations using sorted fractions of CD34+CD117-, CD34+CD117+ and CD34-CD117+ from primary grafts. Flow cytometry plots of sorted human fractions are shown and highlighted cells were transplanted at defined doses into NSG mice. A green plus sign indicates engraftment with CD45+ cells and a red minus sign indicates no engraftment with CD45+ cells in secondarily transplanted mice (n = 2-5 mice for each condition and dose, in total 333 mice, see table S2 for cell frequencies). **(D)** Tertiary xenotransplantations of N-FL and T21-FL grafts in NSG mice for 12 weeks (>0.1% CD45+ cells in bone marrow, n = 5 mice per condition). **(E)** CIBERSORTx analysis to computationally quantify cell type lineage in each of the sorted fractions from N-FL grafts (n = 3 replicates per condition). **(F)** CIBERSORTx analysis as described in (E) for T21-FL (n = 3 replicates per condition).

To evaluate the importance of CD34 and CD117 expression independently, cells from primary xenografts were further sorted into CD34-CD117+, CD34+CD117+ and CD34+CD117-fractions and transplanted at defined doses into secondary NSG recipients (**Fig. 4C and table S2**). For both N-FL and T21-FL controls, only cells from the CD34+CD117+ fraction were able to generate serial grafts at 12 weeks (**Fig. 3C and table S2**). Similarly, only CD34+CD117+ cells from preleukemic T21-FL GATA1s primary grafts were able to generate secondary grafts, with a low preleukemia-initiating cell frequency of ~1/380,000. For STAG2ko primary grafts, both CD34+CD117+ and CD34+CD117-cells were able to engraft in secondary recipients. Notably, for leukemic N-FL and T21-FL GATA1s/STAG2ko primary grafts, cells from both CD34+CD117+ and CD34-CD117+ fractions propagated engraftment in secondary recipients, indicating that CD117 might be a better marker than CD34 for leukemia-initiating cells in Down syndrome leukemia. Both preleukemic and leukemic engraftments were confirmed by the appearance of characteristic blast populations (**fig. S6, F to H**). N-FL and T21-FL GATA1s/STAG2ko grafts but not T21-FL GATA1s grafts could be serially reproduced in tertiary mice (**Fig. 3D**). These findings highlight the transient nature of GATA1s-mediated preleukemia versus the propagating leukemia induced by GATA1s/STAG2ko.

**Fig. 4.**
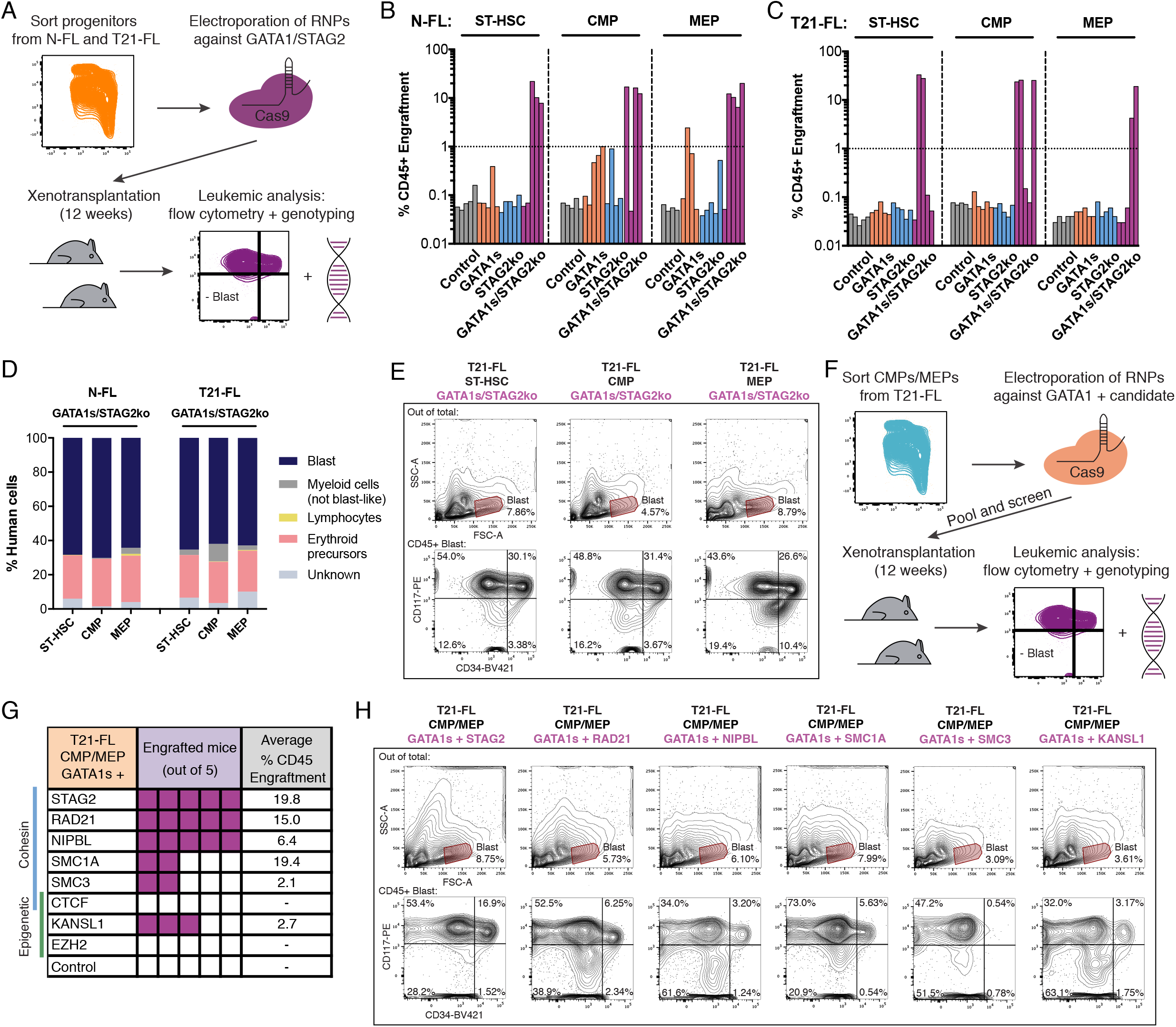
Combined GATA1s and STAG2ko drive leukemic progression in progenitors. **(A)** Experimental overview of sorting N-FL and T21-FL derived progenitor cells for CRISPR/Cas9 editing and transplanting into NSGW41 mice. **(B)** Engraftment levels of N-FL ST-HSC, CMP and MEP grafts in NSGW41 mice (all mice are shown regardless of CD45+ engraftment, n = 4-5 mice per condition). **(C)** Engraftment levels as described in (B) for T21-FL (n = 5 mice per condition). **(D)** Quantification of cell morphology of human cytospinned cells in N-FL and T21-FL GATA1s/STAG2ko grafts in NSGW41 mice from (B) and (C) (n = 400 cells per condition). **(E)** Flow cytometry plots depicting the blast population out of total cells in primary xenografts of T21-FL GATA1s/STAG2ko grafts in NSGW41 mice, as described in (C). The CD34/CD117 profiles out of the CD45+ blast populations are depicted below. **(F)** Experimental overview of sorting T21-FL CMPs and MEPs to conduct a loss-of-function screen to identify genes that endow leukemic progression in combination with GATA1s. **(G)** Result of screen described in (F) showing the number of mice with leukemic phenotypes based on average CD45+ engraftment in bone marrow and blast appearance (>1% CD45 in bone marrow, n = 5 mice per condition). **(H)** Flow cytometry plots of blast populations out of T21-FL CMPs and MEPs edited with GATA1s and candidate gene gRNAs as described in (F).

In order to investigate the mechanism by which GATA1s and STAG2 deficiency contribute to leukemogenesis, specifically within the propagating CD34/CD117 cell fractions from primary xenografts, we carried out transcriptional and epigenetic profiling by RNAseq and ATACseq. CIBERSORTx was utilized to computationally infer the cell lineage contribution from bulk ATAC-Seq (*30*). For this, a signature matrix was generated from normalized read counts over a set of sites unique to individually sorted N-FL HSPC subpopulations (**Fig. S1D and fig. S7, A and B**), including F1, F2 and F3 subgroups of MEPs and CMPs sorted based on CD71 and BAH-1 expression (*24*). Remarkably, engrafting fractions of T21-FL GATA1s LT-HSCs and N-FL and T21-FL GATA1s/STAG2ko LT-HSCs exhibited an increased MEP-like signature compared to control, with the MEP F3 subgroup being the most prominent (**Fig. 3, E and F**). Furthermore, these fractions showed enrichment of GATA-binding motifs at promoters (**fig. S5C and table S3**) associated with an increase in gene expression at these sites (**fig. S5D**). Gene set enrichment analysis of differentially expressed genes between preleukemic versus control and leukemic versus control populations (**table S4**) revealed up-regulation of pathways implicated in translation, ribosome biogenesis and interferon signaling in preleukemic and leukemic populations (**fig. S5E and F, and table S5**). Up-regulated genes were enriched in Down syndrome leukemia blasts from patient samples whereas down-regulated genes were enriched in the stem cell-rich CD34+CD38– fraction of human FL (**fig. S5G**) (*31*). Our results demonstrate that the preleukemic and leukemic propagating populations possess an open chromatin landscape that is largely driven by GATA1s-mediated transcriptional activation. Both of these propagating populations are identifiable by CD117 expression, which suggests that it could serve as a therapeutic target.

### Combined GATA1s and STAG2ko drive leukemic progression in progenitors

The originating cell type in leukemogenesis is increasingly recognized as playing an essential role in the resulting malignancy (*32-34*). To determine whether progeny downstream of LT-HSCs are able to initiate preleukemic or leukemic transformation, we introduced GATA1s and/or STAG2ko into ST-HSCs, CMPs and MEPs and transplanted them into NSGW41 mice at a dose of 1,000 cells (**Fig. 4A**). No consistent human CD45+ engraftment was detected after 12 weeks in mice transplanted with control, GATA1s or STAG2ko cells from either N-FL or T21-FL progenitors (**Fig. 4, B and C**), although limited engraftment of early erythroid lineage cells was observed in mice transplanted with N-FL GATA1s cells (**fig. S8, A and B**). This is consistent with the limited self-renewal potential of progenitors, but also highlights the inability of T21-FL GATA1s progenitor cells to initiate preleukemia in contrast to LT-HSCs. By contrast, human CD45+ engraftment was observed in mice transplanted with N-FL or T21-FL GATA1s/STAG2ko cells, regardless of the differentiation stage of the progenitors (**Fig. 4, B and C**). All grafts generated by N-FL and T21-FL GATA1s/STAG2ko progenitors contained high proportions of CD117+ blasts (**Fig. 4, D and E, and fig. S8C**) accompanied by other phenotypic marker expression typical of Down syndrome leukemia (**fig. S8, D and E**). Finally, cells harvested from grafts generated by N-FL and T21-FL GATA1s/STAG2ko progenitors were able to propagate the leukemia in secondary recipients (**fig. S8, F and G**). Taken together our results suggest that the GATA1s preleukemia-initiating event must occur in LT-HSCs and not downstream progenitors. However, subsequent *STAG2* mutations are not limited to LT-HSCs, but can also be acquired further downstream in the expanded pool of GATA1s-primed progenitor cells, highlighting that the preleukemic and leukemic events can occur in distinct cells of origin.

To explore whether mutations in genes other than *STAG2* can drive leukemic transformation, we carried out a focused loss-of-function screen to evaluate the effects of knocking-out 7 additional genes in T21-FL GATA1s CMPs and MEPs, including 4 additional cohesin genes and 3 genes encoding epigenetic regulators that are frequently mutated in Down syndrome leukemia (*10, 11*). For each gene, 4 gRNAs were individually introduced into T21-FL progenitor cells together with GATA1s, pooled after CRISPR/Cas9 editing and transplanted at a dose of 20,000 cells into NSGW41 mice (**Fig. 4F and table S6**). After 12 weeks, all 5 cohesin gene mutations drove leukemic engraftment in mice (average level of CD45+ engraftment 2-20%), with *STAG2, RAD21* and *NIPBL* being the most potent (**Fig. 4G**). Of the 3 targeted epigenetic regulators, mutations in only *KANSL1* drove leukemic transformation, implying that additional events are needed in the case of *CTCF* and *EZH2* mutations. As expected, control edited T21-FL progenitor cells with GATA1s did not produce any CD45+ grafts. All leukemic grafts contained CD117+ blasts with varying degrees of CD34+ expression (**Fig. 4H**). Interestingly, the blast immunophenotype in the leukemic grafts was similar regardless of the underlying mutation (**fig. S8H**) suggesting that the mutations converge on a common pathway for leukemic transformation. In contrast, a previous loss-of-function screen in a murine model of TAM did not identify cohesin mutations as drivers of Down syndrome leukemogenesis (*10*), implying significant differences between mouse and human systems towards their susceptibility of particular mutations. Altogether, our results show that specific cell types are susceptible to GATA1s-induced pre-leukemia and GATA1s- and cohesin mutations-induced leukemia, underscoring the importance of the cellular context during leukemogenesis.

### Chromosome 21 miRNAs predispose to preleukemia

To investigate the mechanism underlying the synergy between T21 and GATA1s in driving preleukemia development, we analyzed the binding occupancy of GATA1. To do this, we performed Cut&Run assays (*35*) to profile genome-wide GATA1 binding sites and also to quantify binding changes upon GATA1s editing in N-FL and T21-FL CD34+ enriched HSPCs. GATA1s retained many of the binding sites of full length GATA1 as evidenced by the large number of shared peaks (**fig. S9A**), consistent with previously reported findings in a murine cell line (*36*). GATA-binding motifs were highly enriched in these peaks, as were motifs for ETS family members (**fig. S9B**), suggesting binding cooperativity with GATA1. Interestingly, pathway enrichment analysis of GATA1s-specific peaks in T21-FL compared to either control-edited full length GATA1 peaks in T21-FL or to GATA1s peaks in N-FL revealed 13-fold enrichment of promoter sites of genes involved in miRNA loading (**fig. S9, C and D, and table S7**), which was confirmed by gene expression of *AGO1, AGO2, TARBP2* and *ADAR* (**fig. S9E**). These results suggest that GATA1s binding to these miRNA biogenesis genes may increase the potency of miRNA-mediated silencing and post-transcriptional regulation specifically in the T21 context.

In order to explore this idea further, we investigated miRNA expression in T21-FL. We profiled miRNAs from N-FL and T21-FL CD34+ enriched HSPCs by next-generation sequencing. Differential expression of miRNAs on chromosome 21 was not observed, except for let-7c-3p, which showed slightly higher expression in T21-FL CD34+ cells compared to N-FL (**fig. S9, F and G**). However, when chromosome 21 miRNAs were profiled by qPCR in the preleukemic cellular origin of LT-HSCs, as compared to the analysis of bulk CD34+ cells, significant differences were found (**Fig. 5A**). Of particular interest, T21-FL LT-HSCs exhibited up-regulation of miR-99a, miR-125b-2, miR-155 and let-7c compared to N-FL, with the first three having the greatest differential expression.

**Fig. 5.**
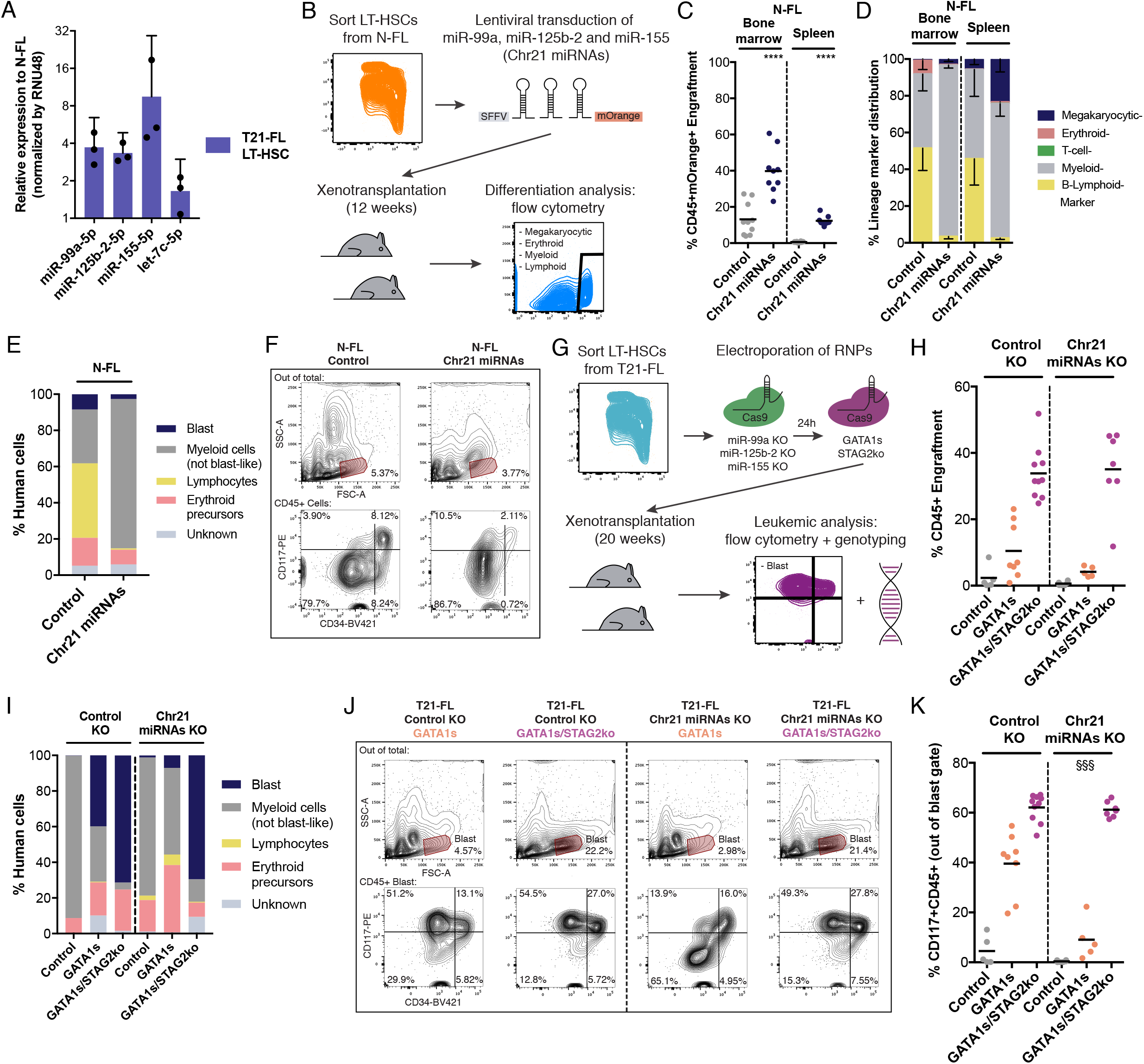
Chromosome 21 miRNAs predispose to preleukemia. **(A)** Relative expression of chromosome 21 miRNAs in T21-FL LT-HSCs compared to N-FL as measured by RT-qPCR (n = 3 replicates per condition). **(B)** Overview of lentiviral mediated overexpression of Chr21 miRNAs in N-FL LT-HSCs used for primary xenotransplantation into NSG and NSGW41 mice. **(C)** Engraftment levels of N-FL transduced control and Chr21 miRNAs LT-HSC grafts in NSG mice (only mice with >1% CD45+ cells in bone marrow were analyzed, n = 9-10 mice per condition). **(D)** Lineage marker distribution based on cell surface markers of engrafted NSG mice in (C). **(E)** Quantification of cell morphology of human cytospinned cells in grafts described in (C) (n = 400 cells per condition). **(F)** Flow cytometry plots showing absence of blast populations out of total cells in primary xenografts described in (C). **(G)** Experimental overview of sorting T21-FL LT-HSCs for CRISPR/Cas9 editing with miR-99a, miR-125b-2 and miR-155 gRNAs and subsequently with GATA1s and STAG2 gRNAs for primary xenotransplantation into NSG mice. **(H)** Engraftment levels of T21-FL control KO and Chr21 miRNAs KO LT-HSC transplanted mice. Each subgroup was additionally edited with control, GATA1s, and GATA1s/STAG2 gRNAs (only mice with >1% CD45+ cells in bone marrow and >90% CRISPR/Cas9 efficiency are depicted, n = 5-10 mice per condition). **(I)** Quantification of cell morphology of human cytospinned cells in transplanted NSG mice described in (H) (n = 400 cells per condition). **(J)** Flow cytometry plots showing blast populations out of total cells in primary xenografts described in (H). **(K)** Quantification of CD117+CD45+ blast of transplanted NSG mice described in (H) (§§§ indicate significance in relation to GATA1s control KO). Unpaired t test: *p < 0.05; **p < 0.01; ***/§§§p < 0.001; ****p < 0.001; error bars represent standard deviation.

To investigate whether our observed T21-specific phenotypes could be recapitulated upon enforced expression of these differentially-expressed chromosome 21 miRNAs, we used lentiviral transduction to overexpress miR-99a, miR-125b-2 and miR-155 (Chr21 miRNAs) together with the fluorescent marker mOrange in N-FL LT-HSCs. Transduced cells were transplanted into NSG and NSGW41 mice and lineage output was assessed at 12 weeks (**Fig. 5B and fig. S9H**). Cells transduced with Chr21 miRNAs generated 2-fold higher engraftment in bone marrow and spleen compared to control-transduced cells (**Fig. 5C**). Chr21 miRNA grafts displayed a strong bias towards increased myeloid, decreased lymphoid and a trend towards increased megakaryocyte differentiation in the transplanted bone marrow (**Fig. 5D and fig. S10, A to E**), similar to the lineage output of control CRISPR/Cas9-edited T21-FL cells in transplanted mice (**Fig. 2D**). Similar grafts were seen in NSGW41 recipients of Chr21 miRNAs LT-HSCs (**fig. S10, F and G**). No abnormal blast populations were detected in any of these grafts by morphologic or flow cytometric analysis (**Fig. 5, E and F, and fig. S10H**). These results demonstrate that simultaneous overexpression of miR-99a, miR-125b-2 and miR-155 in N-FL LT-HSCs is sufficient to generate a T21-like differentiation state.

Next, to examine the role of Chr21 miRNAs in preleukemic initiation and leukemic transformation, Chr21 miRNAs were first knocked out in T21-FL LT-HSCs and then followed by CRISPR/Cas9 editing for GATA1s, with or without STAG2ko, and transplanted into NSG mice (**Fig. 5G and fig. S10, I to K**). Interestingly, knock-out of Chr21 miRNAs combined with GATA1s resulted in a significant reduction in the blast population, including CD117+CD45+ blasts, at 20 weeks post-transplantation (**Fig. 5, H to K**). However, leukemic engraftment or blast accumulation in mice transplanted with T21-FL GATA1s/STAG2ko combined with Chr21 miRNA knock-out were similar to control. Taken together, these results suggest that chromosome 21 miRNAs play an important role in preleukemic initiation but are dispensable for leukemic progression, and raise the possibility that GATA1s directly enhances the potency of these miRNAs.

### CD117/KIT inhibition targets preleukemic-initiating cells and inhibits leukemic progression

Currently, there are no effective treatments to prevent progression from pre-leukemia to leukemia in individuals with Down syndrome. The development of a targeted approach to eliminate preleukemic cells could thus have significant clinical impact for these patients. Our results indicate that CD117/KIT expression marked the cells that mediated the propagation of the GATA1s-induced preleukemia and GATA1s/STAG2ko-induced leukemia. Thus, it is possible that both the preleukemia and leukemia are dependent on KIT signaling for maintenance and progression.

CD117/KIT is a receptor tyrosine kinase that regulates HSC proliferation, maintenance and survival after binding to its ligand, stem cell factor (*37*). To analyze KIT expression in normal hematopoiesis, FL-, umbilical cord blood (CB)- and bone marrow (BM)-derived LT-HSCs were immunophenotypically profiled for CD117 expression. Interestingly, N-FL and T21-FL LT-HSCs contained distinct CD117-low and CD117-high populations (**Fig. 6A**). In contrast, LT-HSCs from N-CB and N-BM showed a single population of cells with a continuum of low to high CD117 expression. Following transplantation at limiting cell dose into NSG mice, only the CD117-high population and not the CD117-low population from N-FL and T21-FL LT-HSCs was able to generate grafts at 20 weeks (**Fig. 6, B and C**). By contrast, both CD117-low and CD117-high N-CB LT-HSCs were able to generate long-term engraftment, albeit with different lineage outputs (**Fig. 6D**). These results suggest that KIT signaling plays an essential role in LT-HSC function during fetal development but may not be required, at least transiently, for LT-HSC function after birth.

**Fig. 6.**
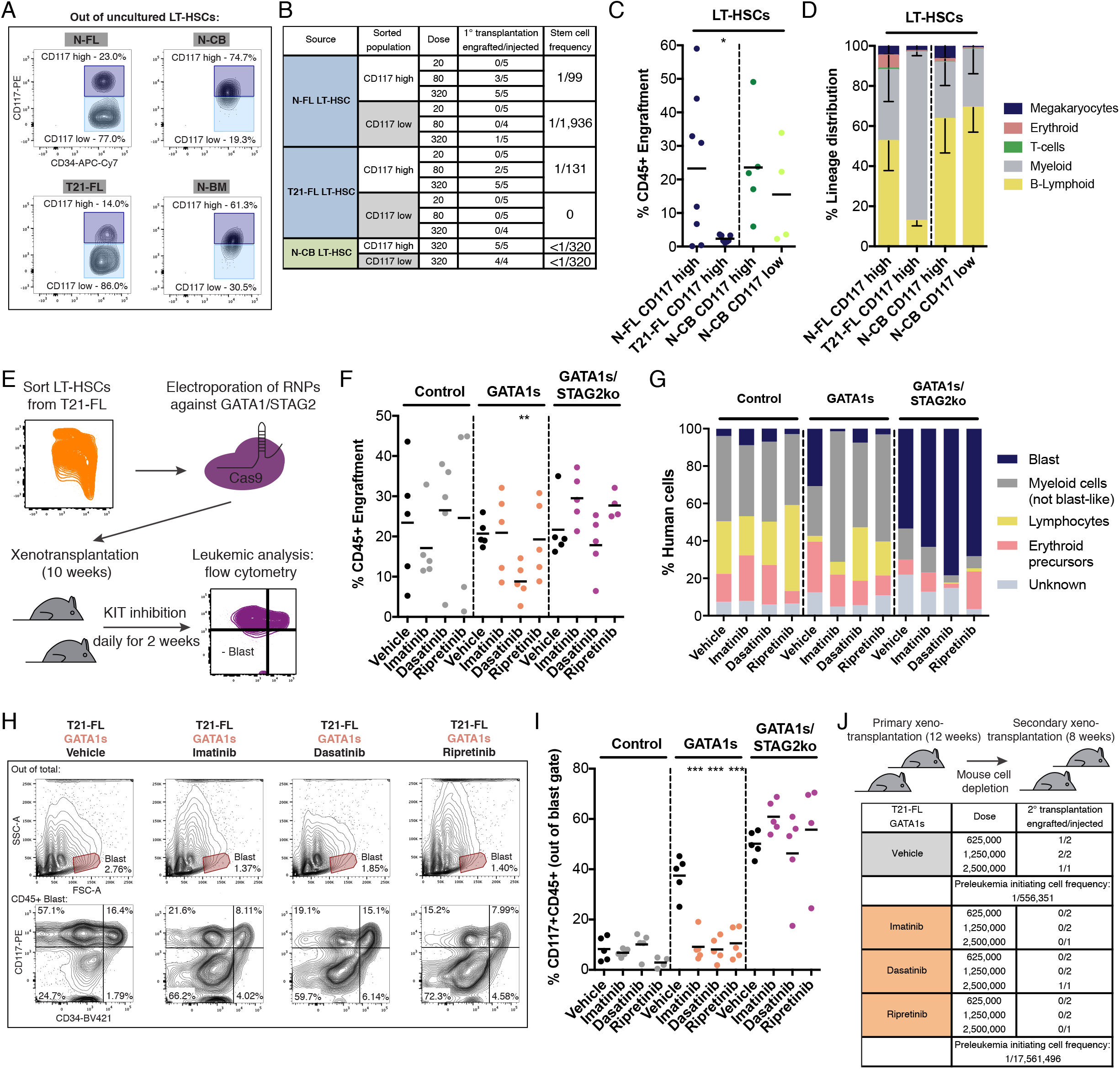
KIT inhibition targets preleukemic-initiating cells and inhibits leukemic progression. **(A)** Immunophenotypic profile of CD117 and CD34 expression of isolated LT-HSCs from N-FL and T21-FL, normal disomic cord blood (N-CB) and normal disomic bone marrow (N-BM). **(B)** CD117-low and CD117-high LT-HSCs were transplanted at defined doses into NSG mice for 20 weeks. Resulting stem cell frequencies are depicted (n = 4-5 mice per condition, in total 67 mice). **(C)** Engraftment levels of N-FL CD117-high, T21-FL CD117-high, N-CB CD117-high and N-CB CD117-low LT-HSCs transplanted NSG mice (>0.1% CD45+ cells in bone marrow, n = 4-8 mice per condition). **(D)** Lineage marker distribution based on cell surface markers of engrafted NSG mice from (C). **(E)** Experimental overview of control-, GATA1s- and GATA1s/STAG2ko-edited T21-FL LT-HSCs transplanted NSG mice, which were subsequently treated twice daily with small molecule inhibitors against KIT for 2 weeks. **(F)** Engraftment levels of T21-FL LT-HSC transplanted NSG mice treated with vehicle, imatinib, dasatinib and ripretinib (n = 4-5 mice per condition). **(G)** Quantification of cell morphology of human cytospinned cells in transplanted NSG mice described in (F). (n = 400 cells per condition). **(H)** Flow cytometry plots showing blast populations out of total cells in primary xenografts described in (F). **(I)** Quantification of CD117+CD45+ blasts in transplanted NSG mice described in (F). **(J)** Secondary xenotransplantations of T21-FL GATA1s grafts from (F). for 8 weeks (>0.1% CD45+ cells in bone marrow, n = 1-2 mice per condition, in total 20 mice). Unpaired t test: *p < 0.05; **p < 0.01; ***p < 0.001; error bars represent standard deviation.

To investigate whether pharmacological inhibition of KIT can target and eliminate preleukemic and leukemic blasts, mice engrafted with 1,300 T21-FL control, GATA1s or GATA1s/STAG2ko LT-HSCs were treated with 1^st^, 2^nd^ and 3^rd^ generation KIT inhibitors (50mg/kg imatinib, 20mg/kg dasatinib or 7.5mg/kg ripretinib) starting 10 weeks posttransplantation with twice-daily dosing for 2 weeks (**Fig. 6E**) (*38-40*). KIT inhibition did not have a significant effect on the overall level of CD45+ engraftment for any group, except for dasatinib-treated mice bearing T21-FL GATA1s preleukemic grafts (**Fig. 6F**). KIT inhibition had no effect on the blast population of leukemic grafts generated by T21-FL GATA1s/STAG2ko cells (**Fig. 6G**). Strikingly, GATA1s preleukemic grafts from mice treated with any of the KIT inhibitors contained significantly lower proportions of blasts compared to vehicle-treated mice, with reduction of CD117+CD45+ blast populations to levels seen in controls (**Fig. 6, H and I, and fig. S10L**). Because some residual CD117+ blasts remained detectable in mice with preleukemic GATA1s grafts (**Fig. 6H**), cells harvested from primary mice were serially transplanted at defined doses into secondary NSG recipients. Cells from KIT inhibitor-treated mice showed a 32-fold lower ability to generate secondary grafts at 8 weeks compared to cells from vehicle-treated mice (**Fig. 6J**). Our results demonstrate that KIT inhibition effectively blocks preleukemic expansion in this pre-clinical setting (**fig. S10M**), supporting further clinical evaluation of this approach as a means to prevent progression to leukemia in Down syndrome newborns.

## Discussion

Our study provides insight into the cellular and molecular mechanism of Down syndrome leukemogenesis, from atypical hematopoiesis associated with T21 to preleukemia initiation and ultimately to leukemic progression. To our knowledge, this study is the first to study T21-FL HSPC subpopulations *in vivo* to model human leukemogenesis within fetal stages. We confirmed that the T21-FL hematopoietic system exhibits an altered phenotypic HSPC hierarchy as previously described (*23, 41, 42*). Although earlier reports have speculated that T21 enhances self-renewal *in vitro* (*42*), our functional studies revealed the opposite; individual T21 HSPC subpopulations exhibited reduced proliferation *in vitro* and generated smaller grafts in xenotransplanted mice with myeloid and megakaryocytic bias and reduced serial transplant ability. These are likely cell-autonomous effects, and may be the basis for the higher incidence of hematopoietic abnormalities such as isolated cytopenias, myelodysplasia and bone marrow failure seen in adults with Down syndrome (*43*). Despite the reduced proliferative capacity of T21 LT-HSCs, our data clearly demonstrate that preleukemia is initiated exclusively in this cellular compartment, contrary to previous hypotheses that megakaryocytic-erythroid progenitor cells are the cell of origin for preleukemia, a prediction derived from their expansion in the HSPC hierarchy of T21-FL (*44*). In fact, the reduced proliferative capacity of T21 LT-HSCs is offset by the acquisition of *GATA1* mutations, providing a possible explanation for the observed selection of *GATA1* mutations in the context of T21. However, the increased function provided by GATA1 mutation comes at a cost including the development of preleukemia and a block in erythrocyte maturation; a result consistent with the lethal anemia seen in GATA1-deficient mice (*45*). In contrast to the LT-HSC origin for preleukemia, leukemic progression can occur in multiple types of downstream progenitors in addition to LT-HSCs. Indeed, the overall pool of progenitors is vastly expanded due to GATA1s-priming, providing a large reservoir for acquisition of secondary mutations in genes such as *STAG2* and thereby increasing the probability of leukemic progression. Enhanced self-renewal mediated by STAG2 deficiency could explain why it is subsequently selected for during leukemic evolution. Selection could also arise from the fact that STAG2 deficiency results in a temporary increase in erythroid output, as seen in our *in vitro* single cell differentiation assays. Erythroid cells make up the vast majority of the numerical daily output of the blood system, raising the possibility that there may be strong evolutionary pressure favoring GATA1s-mutated clones that re-acquire erythroid potential through additional *STAG2* mutation. Overall, our study reveals how critical it is to understand the identity of the cell type that acquires genetic drivers during the development of leukemia.

Our findings establish that initiation of GATA1s-induced preleukemia is dependent on Trisomy 21, which exerts its effects at least partly through up-regulation of chromosome 21 miRNAs, specifically miR-99a, miR-125b-2, and miR-155, exclusively within the LT-HSC compartment. This result refines previous suggestions that deregulated expression of chromosome 21 genes contributes to Down syndrome leukemogenesis and extends it to noncoding RNAs (*46*). Interestingly, these three miRNAs are highly expressed in leukemia-initiating populations of adult acute myeloid leukemias (*47*) and miR-125b-2 has been shown to be a potential oncomiR (*48*). Although preleukemic initiation is dependent on Trisomy 21, we made the surprising finding that progression to leukemia is independent of Trisomy 21 and can be induced by deficiency of STAG2 in combination with GATA1s. We found that preleukemic and leukemic populations were similar with respect to expression of lineage markers on blasts, enrichment of GATA1-binding sites at their promoters and up-regulated pathways compared to control. Nevertheless, the addition of STAG2ko to GATA1s-bearing cells led to enhanced selfrenewal, as evident in the increased frequency of leukemia initiating cells in CD117+ blast populations. Similar leukemic transformation was observed upon induced deficiency of other cohesin genes in combination with GATA1s and thus, it is possible that the effects of these mutations converge on an increase in self-renewal and stemness programs in general. Indeed, the fact that the LSC17 stemness signature is strongly associated with survival across a wide spectrum of acute myeloid leukemia patients irrespective of genetic drivers (*49*) lends support to the concept of convergence. Finally, although our results demonstrate that Down syndrome leukemogenesis can be modeled by a sequence of cell intrinsic mechanisms, we cannot exclude an additional role for the T21 microenvironment on leukemic evolution in Down syndrome individuals.

Through analysis of individual preleukemic and leukemic cell fractions isolated based on primitive stem cell markers and propagated in serial transplants, we identified CD117/KIT as a marker of disease-driving cells. This finding in turn opened up a novel approach for therapeutic targeting in the preleukemic stage where KIT inhibitors are used to block preleukemic stem cells from clonal expansion, limiting the pool of GATA1s-primed progenitors that can acquire additional mutations and ultimately preventing progression to ML-DS. This preclinical study, together with sensitive detection methods for preleukemia (*50*), provides proof of concept that leukemia prevention is feasible. Our findings not only provide novel insight into Down syndrome leukemogenesis, but also have implications for pediatric leukemia in general. Sequencing data from newborns found that the first genetic alterations for many subtypes of childhood leukemia occur during fetal development (*51-53*). Our results potentially suggest that the cellular origin of preleukemic mutations in other pediatric leukemias are also long-term hematopoietic stem cells, which is supported by the fact that it can take years after birth until leukemia is diagnosed (*54*). Early targeting of preleukemia during the newborn phase could become a new clinical paradigm for pediatric leukemia. Lastly, numerical and structural alterations of chromosome 21 are extremely common in hematological malignancies in both children and adults (*55*). In fact, gains of chromosome 21 are seen in up to 35% of cases in several types of acute leukemia (*56*). Thus, it will be important to identify mechanisms including but not limited to dysregulation of chromosome 21 miRNAs that are shared among those cases.

## Acknowledgments

We thank D. Curovic, R. Kelly and J. Law at the Research Centre for Women’s and Infants’ Health Biobank (Mount Sinai Hospital) for sample coordination; B. Chow at the Pathology and Laboratory Medicine (Mount Sinai Hospital) for assistance with pathology; M. DSouza and R. Lopez at the Animal Resources Centre (UHN) for support with mouse work; M. Bergeret, S. Boddeda, S. Ng, A. Srinath, O. Subedar, A. AuYeung and S. Zhao at the SickKids-UHN Flow and Mass Cytometry Facility for assistance with flow cytometry; B. Apresto at tThe Centre for Applied Genomics (SickKids) for Sanger sequencing; K. Ho at the Centre for Applied Genomics (SickKids) for next generation sequencing; A. Smith at the Cancer Cytogenetics Laboratory (UHN) for karyotyping; K. Asoyan, C. Cimafranca, J. Mouatt, M. Peralta and Y. Yang at the Pathology Research Program (UHN) and N. Law at the Sttarr Innovation Centre (UHN) for assistance with histology.; M. Bartolini at the Advanced Optical Microscopy Facility (UHN) for slide scanning; J. Moffat, K. Brown and C. Ross at the Donnelly Centre for supplying the gRNA sequences for the *in vivo* screen; S. Henikoff for supplying the Protein A-Micrococcal nuclease fusion protein for the Cut&Run assay; L. Shultz at the Jackson Laboratory for providing NSGW41 mice; the labs of S. Chan and F. Notta for sharing equipment. We thank C. Jones, A. Tikhonova and members of the Dick lab for comments on the manuscript.

## Author contributions

E.W., J.E.D. and E.R.L. conceived the project, supervised research and wrote the paper. J. A., O.I.G., G.K. and J.C.Y.W. edited the paper. E.W., J.A., O.I.G. and E.R.L analyzed experiments, E.W., J.A., S.K.C., M.A., S.A.S. performed *in vitro* and *in vivo* experiments. O.I.G and E.R.L assisted with mouse work. J.A. performed morphological analysis. O.I.G. assisted with single cell assays. A.M. analyzed ATACseq and RNAseq data. G.K. performed western blot assays, Cut&Run assays and prepared miRNA libraries. J.L.M. assisted with intrafemoral injections. S.A.M. performed Cut&Run analysis. M.G. analyzed miRNA sequencing data. M.C. performed CRISPR/Cas9 off-target analysis. L.G.P. assisted with ATACseq library preparations. K. C., M.R., P.S. and D.C. coordinated patient consent and sample collection. J.C.Y.W. and J.K.H. provided study consultation. J.E.D. secured funding for this study.

## Data and materials availability

Raw sequence data will be available at European Genome-phenome Archive and processed data will be available at Gene Expression Omnibus. All other data is available in the main text or the supplementary materials.

## Materials and Methods

### Animal studies

All mouse experiments were approved by the University Health Network (UHN) Animal Care Committee and we confirm that all experiments conform to the relevant regulatory and ethical standards. All xenotransplantations were performed in 8-to 12-week-old female *NOD.Cg-Prkdc^scid^Il2rg^tm1Wjl^Kit^em1Mvw^/SzJ* (NSG) mice (JAX) that were sublethally irradiated with 225cGy, 24 hours before transplantation, or in 8-to 12-week-old female *NOD.Cg-Prkdc^scid^Il2rg^tm1Wjl^Kir^m1Mm^7SzJ* (NSGW41) mice that were not irradiated. NSGW41 mice used in this study were a kind gift from Leonard D. Shultz. NSGW41 mice were utilized because they support long-term engraftment of human hematopoietic cells without the need for sub-lethal irradiation and provide better support for erythroid and megakaryocytic differentiation compared to NSG mice. Female littermates were randomly assigned to experimental groups.

### Human patient samples

Human fetal liver samples were obtained from elective pregnancy terminations at Mount Sinai Hospital with informed consent in accordance to guidelines approved by the Mount Sinai Hospital Research Ethics Board and the University Health Network (UHN) Research Ethics Board. Fetal liver samples of normal disomic karyotype and trisomy 21 were collected at 16 to 19 weeks gestation from either sex. For the single cell *in vitro* assays, only male samples were used since GATA1 and STAG2 are located on the X-chromosome. For the *in vivo* assays, both male and female samples were utilized.

### Fetal liver CD34+ isolation

Fetal liver samples were obtained from Mount Sinai Hospital and processed within 1-3 hours. For this, the fetal liver sample was finely minced using razor blades (VWR) and was then put into two 50ml Falcon tubes, each with 40ml of pre-warmed IMDM media (Thermo Fisher), 5 ml of Collagenase IV (Stem Cell Technologies) and 25μl of 10 mg/ml DNAse I (Roche). The 50ml tubes were incubated on a shaker at 37°C for 30min. The dissociated sample was then filtered through a 40μm cell strainer (Corning) and the remaining tissue pieces were pushed through the strainer using the black rubber end of a 5ml syringe (BD). Subsequently, the cells were spun at 350xg for 10min and the red blood cells were lysed with 5ml Ammonium chloride (StemCell Technologies) per 50ml tube for 5min. Lysis was stopped with IMDM media and cells were spun at 350xg for 10min. The remaining cell pellets were combined and re-suspended in 500μl MACS auto running buffer (Miltenyi Biotec). Subsequently, CD34+ cells were enriched using the human CD34 MicroBead kit (Miltenyi Biotec) according to the manufacturer’s protocol. In total, 12 LS columns (Miltenyi Biotec) were used per fetal liver sample. CD34+ fetal liver cells were viably stored in 90% FBS (GE Healthcare) and 10% DMSO (FisherScientific) at −150°C.

### Fetal liver sex determination

For each fetal liver sample, the sex was determined using a PCR-based approach as previously described (*57*). For this, a small aliquot of CD34-cells from each fetal liver was utilized. The sample was denatured at 95°C for 5min and then centrifuged at 12,000xg for 5min. Subsequently, the supernatant was used as input at several dilutions (undiluted, 1:2 and 1:5). Each PCR reaction contained 5μl of undiluted or diluted crude genomic DNA, 1μl of forward and reverse primer (10μM, IDT), 18μl nuclease-free H_2_O (IDT) and 25μl of AmpliTaq Gold 360 master mix (Thermo Fisher). The PCR program was: 95°C for 10min, followed by 95°C for 30s, 56°C for 30s and 72°C for 1min (35 cycles) and then 72°C for 10min. Two sets of PCR primers were used: SRY primers to detect the Y-chromosome and ATL1 primers to detect the X-chromosome(s). PCR products were run on a 1.5% Agarose gel (Thermo Fisher) and the sex was determined based on the observance of positive PCR bands (250bp for SRY and 300bp for ATL1).

### Trisomy 21 verification

Fetal liver samples were clinically identified with trisomy 21 through prenatal screening. In order to confirm trisomy 21 status, genomic DNA was isolated from CD34+ cells using the Agencourt GenFind V3 kit (Beckman Coulter) and subjected to droplet digital PCR (ddPCR, Bio-Rad) to assess chromosome 21 copy number variations. For this, two sets of probes were used separately. In the first set NCAM2 (FAM) was utilized to detect chromosome 21 and ASTN1 (HEX) used as a reference to detect chromosome 1. In the second set RUNX1 (FAM) was used to detect chromosome 21 and SMAD4 (HEX) used as a reference to detect chromosome 18. Each ddPCR reaction contained 1.6μl of genomic DNA (15ng/μl), 1μl HaeIII (2U/μl in 1x CutSmart Buffer, NEB), 1μl of HEX primers/probe assay, 1μl of FAM reference primers/probe assay, 5.4μl nuclease-free H_2_O (IDT) and 10μl ddPCR Supermix (no dUTP, Bio-Rad). The ddPCR was carried out using the QX200 ddPCR system according to manufacturer’s protocol. The ddPCR program was: 95°C for 10min, followed by 94°C for 30s, 60°C for 1min (45 cycles) and then 98°C for 10min. Copy number variation was determined using the QuantaSoft ddPCR software (Bio-Rad) using ASTN1 or SMAD4 as the reference.

### Fetal liver sorting

CD34+ fetal liver cells were thawed via slow dropwise addition of thawing media, made up with X-VIVO 10 media (Lonza) with 50% FBS (GE Healthcare) and DNase I (100 μg/ml, Roche). Cells were spun at 350xg for 10min and then re-suspended in PBS (Thermo Fisher) + 2.5% FBS. For all *in vitro* and *in vivo* experiments, the full stem and progenitor hierarchy was utilized to sort LT-HSCs, ST-HSCs, CMPs and MEPs (*24*). For this, CD34+ fetal liver cells were resuspended in 100μl PBS + 2.5% FBS per 1×10^6 cells and stained in two subsequent rounds for 20min each at room temperature. For the first round, the following antibodies were used (volume per 1×10^6 cells): CD45RA FITC (5μl, BD), CD49f PE-Cy5 (3.5μl, BD), CD10 BV421 (4μl, BD), CD19 V450 (4μl, BD) and FLT3 CD135 biotin (12μl, BD). Afterwards, cells were washed and the following antibodies were used for the second round (volume per 1×10^6 cells): CD45 V500 (4μl, BD), CD34 APC-Cy7 (3μl, BD), CD38 PE-Cy7 (2.5μl, BD), CD90 APC (4μl, BD), CD7 AF700 (10μl, BD) and Streptavidin Conjugate Qdot 605 (3μl, Thermo Fisher). Cells were sorted on the FACSAria III (BD). LT-HSCs were sorted as CD45+CD34+CD38-CD45RA-CD90+CD49f+, ST-HSCs as CD45+CD34+CD38-CD45RA-CD90-CD49f-, CMPs as CD45+CD34+CD38+CD10/19-CD7-CD45RA-FLT3+ and MEPs as CD45+CD34+CD38+CD10/19-CD7-CD45RA-FLT3-. Cell sorting purity checks were performed after each sorted sample (>95%).

### CRISPR/Cas9 RNP electroporation

Sorted LT-HSCs, ST-HSCs, CMPs and MEPs were cultured for 36-48 hours in serum-free X-VIVO 10 media (Lonza) supplemented with 1% Bovine Serum Albumin Fraction V (Roche), 1x L-Glutamine (Thermo Fisher), 1x Penicillin-Streptomycin (Thermo Fisher) and the following cytokines (Miltenyi Biotec): FLT3 Ligand (100ng/mL), G-CSF (10ng/mL), SCF (100ng/mL), TPO (15ng/mL) and IL-6 (10ng/mL). Cells were cultured in 96 well round-bottom plates (Corning). CRISPR/Cas9 RNP electroporations were carried out using chemically synthesized gRNAs (IDT), recombinant Cas9 nuclease (IDT) and the 4D-Nucleofector (Lonza) as previously described (Wagenblast et al., 2019). For this, Alt-R CRISPR/Cas9 crRNA and tracrRNA (IDT) were re-suspended to 200μM with TE Buffer (IDT), mixed 1:1 and annealed in a thermocycler at 95°C for 5min, then cooled to room temperature. If using two gRNAs to target the same gene such as for GATA1s, both cRNAs were annealed to the tracrRNA in a single tube at a 1:1:2 ratio. For each electroporation reaction, 1.2μl crRNA:tracrRNA, 1.7μl Cas9 protein and 2.1μl PBS (Thermo Fisher) were combined in a low-binding Eppendorf tube (Axygen) and incubated for 15min at room temperature. If two genes were targeted in the same electroporation reaction such as for GATA1s/STAG2ko, 1μl of each crRNA:tracrRNA, 1.7μl Cas9 protein and 1.4μl PBS were combined into the same Eppendorf tube. After incubation, 1μl of 100μM electroporation enhancer (IDT) was added. Pre-cultured cells were washed with pre-warmed PBS and spun down at 350xg for 10min. Between 1×10^3 – 5×10^4 cells were re-suspended in 20μl Buffer P3 (Lonza) per reaction and quickly added to the Eppendorf tube containing the CRISPR/Cas9 gRNA RNP complex. This mixture was mixed briefly by pipetting up and down and transferred to the electroporation chamber (Lonza). The cells were electroporated using the 4D-Nucleofector with the program DZ-100. Immediately afterwards, 180μl of pre-warmed X-VIVO 10 media (as described above) was added to the electroporation chamber and transferred to a 96 well roundbottom plate. The electroporated cells were recovered overnight at 37°C in the incubator.

### gRNA sequences

The gRNA sequences for control and GATA1s were previously described (*21*). For control, two gRNAs were utilized that target the olfactory receptor OR2W5. The OR2W5 and STAG2 gRNAs were predicted using the CRoatan algorithm (*58*). For GATA1s, two gRNAs were utilized that target the 5’ and 3’ end of exon 2. Simultaneous use of both gRNAs enabled complete excision of exon 2, resulting in GATA1s.. The gRNA sequences for miR-99a, miR-125b-2 and miR-155 were designed with Benchling. For this, gRNA sequences were considered optimal that were proximal to the 5p strand of each miRNA. For control, two gRNAs against the olfactory receptor OR10A4 were used, which were predicted from the CRoatan algorithm. Finally, gRNA sequences for the loss of function *in vivo* screen were kindly provided by Jason Moffat using their algorithm (Table S6) (*59, 60*).

### CRISPR/Cas9 off-target analysis

To assess CRISPR/Cas9 off-targets, CRISPR/Cas9-edited N-FL LT-HSCs were expanded in X-VIVO 10 media (as described above for CRISPR/Cas9 RNP electroporation) for 7 days. As a reference, non-electroporated LT-HSCs were used. Genomic DNA was isolated using the Agencourt GenFind V3 kit (Beckman Coulter) and next-generation sequencing libraries were prepared using the KAPA HyperPrep kit (Roche). Subsequently, the libraries were sequenced at 30X coverage on the NovaSeq S4 using 150bp paired-end sequencing. Targeted gRNA sequences for each sample were blasted against the GRCh38 genome using blastn to identify regions of the genome with similar sequences (*61*). Whole genome sequencing data was aligned to GRCh38 using bwa mem (*62*) and the data was collapsed using picard. Variants, specifically indels, were then called using bcftools within 25bp flanking regions of each location identified by blastn. Default parameters were used for all tools.

### Karyotyping analysis

CRISPR/Cas9-edited CD34+CD38-enriched stem cells were expanded in IMDM (Thermo Fisher), supplemented with 10% FBS (GE Healthcare), 1x L-Glutamine (Thermo Fisher), 1x Penicillin-Streptomycin (Thermo Fisher) and the following cytokines (all from Miltenyi Biotec): FLT3L (100ng/mL), G-CSF (10ng/mL), SCF (100ng/mL), TPO (15ng/mL) and IL-6 (10ng/mL) for 10 days. Karyotyping of chromosomes was prepared according to standard procedures. Briefly, metaphase slides were prepared, banded with trypsin and stained with Leishman’s stain. The G-banded slides were scanned and metaphases were analyzed using the imaging system Ikaros (Meta Systems). For each condition, 20 metaphases were analyzed by G-banded karyotyping to assess numerical and structural abnormalities.

### Single cell *in vitro* assay

Setup of single cell assay: Single cell *in vitro* assays were performed as previously described (*21*). For this, three days prior to the single cell assay, Nunc 96-well flat bottom plates (Thermo Fisher) were treated with 50μl 0.2% Gelatin solution (Sigma-Aldrich) per well for one hour. After the Gelatin solution was removed, MS-5 murine stromal cells (*63*) were seeded at a density of 1,500 cells per well in 100μl H5100 media (Stem Cell Technologies). One day prior to the single cell assay, the H5100 media was replaced with 100μl of erythro-myeloid promoting media, which consisted of StemPro-34 SFM media (Thermo Fisher) with the provided supplement, 1x L-Glutamine (Thermo Fisher), 1x Penicillin-Streptomycin (Thermo Fisher), 0.02% Human LDL (Stem Cell Technologies) and the following cytokines (Miltenyi Biotec unless stated otherwise): FLT3L (20ng/mL), GM-CSF (20ng/mL), SCF (100ng/mL), TPO (100ng/mL), EPO (3ng/mL, Eprex), IL-2 (10ng/mL), IL-3 (10ng/mL), IL-6 (50ng/mL), IL-7 (20ng/mL) and IL-11 (50ng/mL). On the day of the single cell assay, electroporated cells were stained with 1:2000 Sytox Blue (Thermo Fisher) in PBS + 2.5% FBS and viable single cells were sorted and deposited directly onto the MS-5 seeded 96-well plates using the FACSAria II (BD). Single cells were cultured for 17 days and 100μl of fresh erythro-myeloid promoting media was added at day 8.

Flow cytometry of single cell assay: Individual wells with human cells were marked and these numbers were used to calculate single cell colony efficiencies. Cells were removed from each well and transferred to a 40μm 96-well filter plate (Pall) to remove MS-5 murine stromal cells. The filter plate was put on top of a 96-well round bottom plate (Corning) and centrifuged at 300xg for 7min. The supernatant was removed by inverting the round bottom plate, leaving around 20μl of liquid per well. Afterwards, cell pellets were re-suspended by adding 30μl of PBS + 2.5% FBS. 25μl was transferred to a PCR plate (Eppendorf) and stored at −80°C for subsequent genotyping. The remaining 25μl of cells were mixed with 25μl of antibody mix and stained for 45min at 4°C. The following antibodies were used: CD45 APC (1:100, BD), CD34 APC-Cy7 (1:250, BD), CD33 BV421 (1:50, Biolegend), CD71 FITC (1:100, BD), CD41 PE-Cy5 (1:200, Beckman Coulter) and GlyA PE (1:200, Beckman Coulter). Finally, 100μl of PBS + 2.5% FBS was added to each well and 100ml was analyzed on the FACSCelesta with a high throughput sampler (HTS, BD). All flow cytometry analysis was performed using FlowJo (BD) in a blinded manner. Generally, greater than 10 cells were required to call a positive lineage. Erythroid cells were identified by GlyA expression and CD71+ cells were identified by CD71 expression and lack of GlyA expression. For cell proliferation, absolute numbers of CD45+ and CD41+ cells were counted and multiplied by 3 to obtain total number of cells per well. Colonies containing only GlyA+ erythroid colonies were excluded in the CD45+ proliferation analysis.

Genotyping of single cell assay: Genomic DNA was isolated from each single cell colony using the Agencourt GenFind V3 kit. The isolation was performed using 96-well PCR plates (Eppendorf) and a magnetic stand (Thermo Fisher) according to manufacturer’s protocol, but with modified volumes: 25μl lysis buffer, 1.2μl of Proteinase K (Zymo Research), 50μl magnetic particles, 200μl wash buffer 1, 125μl wash buffer 2 and 60μl TE buffer (IDT) for elution. PCR was used to amplify the CRISPR/Cas9 modified genomic locus and each PCR reaction contained 23μl of genomic DNA, 1μl of forward and reverse primer (10μM, IDT) and 25μl of AmpliTaq Gold 360 master mix (Thermo Fisher). The PCR program was: 95°C for 10min, followed by 95°C for 30s, 56°C for 30s and 72°C for 1min (40 cycles) and then 72°C for 7min. To identify colonies with control (OR2W5) gRNA mediated heterozygous and homozygous cleavage, PCR products were run on a 1.5% Agarose gel and were screened for a shift from 700bp to 500bp. Both, heterozygous and homozygous control gRNA colonies were used in the analysis. To identify colonies with the GATA1s genotype, PCR products were run on a 1.5% Agarose gel (Thermo Fisher) and were screened for a shift in size from 1000bp to 550bp. Lastly, to identify colonies with STAG2 knock out genotypes, PCR products were purified and then sent to Sanger sequencing using the reverse PCR primer. Chromatograms were inspected to identify colonies that showed a frameshift mutation. For the single cell *in vitro* assays, only male samples were used since both GATA1 and STAG2 are located on the X-chromosome, resulting in only one allele needing to be edited. CRISPR/Cas9 editing efficiencies were calculated based on the number of positive colonies among all samples that displayed a band in the gel electrophoresis or a signal in the chromatogram.

### Cell cycle analysis

N-FL and T21-FL were utilized to sort LT-HSCs, ST-HSCs, CMPs and MEPs by flow cytometry in a two-step staining protocol. For the first round, the following antibodies were used (all from BD, unless stated otherwise): CD45RA FITC (1:50, BD), CD90 BV650 (1:50, BD), CD34 APC-Cy7 (1:200), CD38 PE-Cy7 (1:100, BD), CD49f PE-Cy5 (1:50) and FLT3 CD135 biotin (1:50, BD). Cells were washed and the following antibody was used for the second round: Streptavidin Conjugate Qdot 605 (1:200, Thermo Fisher). Around 1-3×10^4 cells were obtained from each subpopulation and cells were subsequently cultured in X-VIVO 10 media overnight (as described above for CRISPR/Cas9 RNP electroporation) and then pulsed with EdU for 2 hours using the Click-iT EdU Pacific Blue Flow Cytometry Assay Kit according to manufacturer’s protocol. After fixation and the Click-iT reaction, cells were stained with Ki67 APC (1:30, BD) overnight at 4°C. The next day, cells were stained with Propidium Iodide (1:500) and DNAse-free RNAseA (10μg/ml, Ambion) for 15 min and then analyzed on the FACSCelesta (BD).

### Western blot assay

CMPs and MEPs were sorted by flow cytometry, then combined and edited with CRISPR/Cas9 (as described in CRISPR/Cas9 RNP electroporation). After electroporation, the cells were cultured in erythroid-myeloid differentiation media (as described above for single cell *in vitro* assay) for 3 days. 2×10^5 cells were lysed in RIPA buffer (Thermo Fisher) with protease and phosphatase inhibitors (Thermo Fisher). Then samples were centrifuged at 12,000xg for 10min at 4°C and the supernatants were subsequently used. Western blot assays were performed on the automated Simple Western capillary platform (Wes, Protein Simple) using 12-230kDa capillary cartridges according to manufacturer’s protocol. Antibodies were first titrated on CD34+ fetal liver lysates and subsequently used at the following dilutions: Anti-GATA1 (Cell Signaling, 1:5), anti-STAG2 (Cell Signaling, 1:50) and anti-GAPDH (Cell Signaling, 1:300).

### Near-clonal xenotransplantation

Setup of xenotransplantation: Xenotransplantations were carried out as intrafemoral injections (*64*). For this, mice were anesthetized with isoflurane and the right knee was secured in a bent position and a hole was drilled into the right femur with a 27gauge needle. This was followed by injection of CRISPR/Cas9-edited cells in 30μl PBS using a 28gauge ½ cc syringe (Becton Dickinson, 329461). Primary xenotransplantations of CRISPR/Cas9-edited LT-HSCs transplanted mice were sacrificed at week 20, whereas CRISPR/Cas9-edited ST-HSCs, CMPs and MEPs transplanted mice were sacrificed at week 12. For this, mice were sacrificed to obtain the injected right femur (bone marrow). Bones were flushed with 1ml PBS + 2.5% FBS. In addition, spleens were isolated and crushed using the end of the plunger of a 3 ml syringe (BD) and subsequently pushed through a 5ml tube with a 35μm cell strainer (Corning). All collected cells were centrifuged at 350xg for 10min. Then cells were re-suspended in 500μl and counted in 500μl ammonium chloride (Stem Cell Technologies) using the Vicell XR (Beckman Coulter). Subsequently, 25μl of cells were transferred to a PCR plate (Eppendorf) and stored at −80°C for subsequent genotyping using the same protocol as described for single cell *in vitro* assay.

Analysis of xenotransplantation: For flow cytometry, 50μl of cells were mixed with 50μl of antibody mix and stained for 60min at 4°C. The following antibodies were used (all from BD, unless stated otherwise): CD45 AF700 (1:100), CD33 APC (1:100), CD19 V450 (1:100), CD41 PE-Cy5 (1:200, Beckman Coulter), GlyA PE (1:100, Beckman Coulter), CD71 BV650 (1:100), CD3 FITC (1:100) and CD34 APC-Cy7 (1:100). For experiments that evaluated the CD117 blast burden, the following antibodies were used (all from BD, unless stated otherwise): CD45 AF700 (1:100), CD33 BV786 (1:100), CD19 V450 (1:100), CD41 PE-Cy5 (1:200, Beckman Coulter), GlyA PE (1:100, Beckman Coulter), CD71 BV650 (1:100), CD3 FITC (1:100), CD34 APC-Cy7 (1:100) and CD117 APC (1:100). Cells were analyzed on the FACSCelesta (BD) with a high throughput sampler (HTS). Remaining unstained cells from each femur were viably frozen and stored at −150°C. All flow cytometry analysis was performed using FlowJo (BD) in a blinded manner.

Genotyping of xenotransplantation: For each CRISPR/Cas9 electroporation, a small subset of cells was cultured in X-VIVO 10 media (as described above for CRISPR/Cas9 RNP electroporation) for 5-7 days to validate CRISPR/Cas9 efficiency. In order to assess CRISPR/Cas9 efficiency in each xenograft, genomic DNA was isolated from bulk cells of bone marrow. The CRISPR/Cas9 modified genomic locus was PCR amplified (as described above for the single cell *in vitro* assay). Sanger sequencing was carried out using the reverse PCR primer and chromatograms were analyzed using TIDE (*65*) to verify CRISPR/Cas9 editing in control, GATA1s, STAG2ko, miR-99a, miR-125b-2 and miR-155 edited bulk cells. For control and GATA1s-edited xenografts, where a large deletion of 200bp and 400bp is expected, CRISPR/Cas9 efficiency was determined based on the percentage of aberrant sequences after the gRNA cut site.

Only mice that showed a CRISPR/Cas9 efficiency of >90% and a CD45+ engraftment level in the bone marrow of >1% were utilized in the analysis. For the CD45+ engraftment level graphs, all corresponding spleen samples were shown, regardless of engraftment. For the subsequent cell differentiation graphs, only spleen samples with CD45+ engraftment levels of >1% were included in the analysis. For the xenotransplantations, one female T21-FL sample was used in each NSG and NSGW41 cohort and the reaming samples were male.

In xenotransplantations of CRISPR/Cas9-edited male fetal liver samples, clonal engraftment was visible in GATA1s-, STAG2ko- and GATA1s/STAG2ko-edited mice. CRISPR/Cas9-edited mice with more than one clone were included in the analysis as long as the CRISPR/Cas9 efficiency of >90% and engraftment criteria of >1% were satisfied.

### Limiting dilution *in vivo* assay

For limiting dilution primary xenotransplantations, cells were transplanted at defined doses into 8-to 12-week old female NSG or NSGW41 mice. Cell numbers were based on the number of cells sorted at flow cytometry. Cell frequencies were estimated using ELDA (*66*).

### Methylcellulose colony formation assay

For methylcellulose colony formation assays, 2×10^4 cells from bone marrow of xenotransplanted mice were transferred to 2ml of MethoCult H4034 optimum methylcellulose medium (Stem Cell Technologies) and plated onto two 35mm dishes. Only human-specific colonies were able to grow on methylcellulose. After 12 days, individual colonies were picked in order to genotype CRISPR/Cas9-edited loci. For this, 12 individual colonies per sample were transferred to a 96-well PCR plate (Eppendorf), washed with PBS and stored at −80°C. The number of individual unique CRISPR/Cas9 edits were determined after genomic DNA isolation, PCR amplification and Sanger sequencing with the reverse PCR primer to assess overall clonality (as described for single cell *in vitro* assay).

### Cellular morphology

In primary and secondary xenotransplantations, multiple femur of engrafted mice were combined for each condition and murine cells were depleted using the Mouse cell depletion kit (Miltenyi Biotec). Cells were stained for 15min at 4°C and then transferred to an LS column (Miltenyi Biotec). Human cells were then cytospinned with Shandon cytofunnels (Thermo Fisher) onto glass slides using the CytoSpin 4 instrument (Thermo Fisher) at 112xg for 10min with medium acceleration. Slides were air-dried overnight and then stained for standard Giemsa staining (Sigma). All quantifications were performed in a blinded manner.

### Immunohistochemistry

During primary xenotransplantations, the right humerus of each mouse was isolated and fixed in a 10% Formalin solution (Sigma) for 24 hours at room temperature. Subsequently, the bones were decalcified using a 12% EDTA pH 8.0 solution (Sigma) for 3-4 days. Serial 5μm sections were then prepared. One section was stained with a standard haematoxylin and eosin protocol. The other sections were used for immunohistochemistry against human CD45 and CD61. For CD45, slides were treated with Tris EDTA pH 9.0 and for CD61, slides were treated with Pepsin to retrieve antigens. Subsequently, anti-CD45 and anti-CD61 were used in a 1:2000 dilution for 1 hours and 1:50 dilution overnight, respectively. CD45 and CD61 diaminobenzidine-stained (DAB) and haematoxylin stainings were scanned on the AT2 Slide Scanner (Aperio) and visualized using QuPath (*67*). CD61 DAB and haematoxylin stainings were quantified using ImageJ (*68*). For this, images were colour deconvoluted (*69*) and the mean intensity of CD61 positive staining was measured. All quantifications were performed in a blinded manner.

### Blast panel analysis

In primary, secondary and tertiary xenotransplantations, multiple femur of engrafted mice were combined for each condition and stained with four different antibody panels. For this, 50μl cells were mixed with 50μl of antibody mix and stained for 60min at 4°C. The following antibodies were used. Panel 1 to stain against erythroid makers (all from BD, unless stated otherwise): CD45 AF700 (1:100), CD34 BV421 (1:100), CD41 APC-Cy7 (1:100, Biolegend), CD117 PE (1:100), CD71 BV650 (1:100), CD36 APC (1:100), GlyA PC5 (1:100) and CD33 BV786 (1:100). Panel 2 to stain against megakaryocytic and aberrant makers (all from BD, unless stated otherwise): CD45 AF700 (1:100), CD34 BV421 (1:100), CD41 APC-Cy7 (1:100, Biolegend), CD61 FITC (1:100), CD42b APC (1:100), CD7 PE (1:100), CD4 PC5 (1:100, Beckman Coulter) and CD56 BV605 (1:100). Panel 3 to stain against myeloid and aberrant makers (all from BD, unless stated otherwise): CD45 AF700 (1:100), CD34 BV421 (1:100), CD41 APC-Cy7 (1:100, Biolegend), CD117 PE (1:100), CD33 BV786 (1:100), CD13 APC (1:100), HLA-Dr PC5 (1:100) and CD14 BV605 (1:100). Lastly, panel 4 to stain against myeloid makers (all from BD, unless stated otherwise): CD45 AF700 (1:100), CD34 BV421 (1:100), CD41 APC-Cy7 (1:100, Biolegend), CD117 PE (1:100), CD33 BV786 (1:100), CD13 APC (1:100), CD14 BV605 (1:100) and CD11b PC5 (1:100, Beckman Coulter). Cells were analyzed on the FACSCelesta (BD).

### Secondary transplantation

Setup of secondary xenotransplantation: Individual femur of primary xenotransplantations, which were validated for CRISPR/Cas9 edits, were thawed (as described in fetal liver sorting). On average, 8 femur samples were thawed per condition. Then, the femur samples were combined and murine cells were depleted using the Mouse cell depletion kit (Miltenyi Biotec). Cells were stained for 15min at 4°C and then transferred to LS columns (Miltenyi Biotec). Subsequently, human cells were re-suspended in 100μl per 1×10^6 cells and stained for 60min at 4°C. The following antibodies were used (volume per 1×10^6 cells): CD34 APC-Cy7 (2μl, BD), CD117 PE (2μl, BD). Afterwards, the cells were stained with 1:2000 Sytox Blue (Thermo Fisher) in PBS + 2.5% FBS. For injections into NSGW41 secondary recipients, CD34+ and/or CD117+ cells were viably sorted using the FACSAria III (BD). For injections into NSG secondary recipients, three populations were sorted: CD34-CD117+, CD34+CD117+ and CD34+CD117-. Subsequently, cells were transplanted at defined doses via intrafemoral injections. Secondary xenotransplantations were sacrificed at week 12 and the injected right and left femur were isolated. Engraftment was considered positive if CD45+ engraftment level was >0.1% in the bone marrow.

Analysis of secondary xenotransplantation: For flow cytometry, 50μl of cells were mixed with 50μl of antibody mix and stained for 60min at 4°C. The following antibodies were used: CD45 AF700 (1:100), CD33 BV786 (1:100), CD19 V450 (1:100), CD41 PE-Cy5 (1:200, Beckman Coulter), GlyA PE (1:100, Beckman Coulter), CD71 BV650 (1:100), CD3 FITC (1:100), CD34 APC-Cy7 (1:100) and CD117 APC (1:100). Cells were analyzed on the FACSCelesta (BD) with a high throughput sampler (HTS). Remaining unstained cells from each right and left femur were viably frozen and stored at −150°C. All flow cytometry analysis was performed using FlowJo (BD) in a blinded manner.

### Tertiary transplantation

At the time of analysis of the secondary transplantations, femurs of engrafted mice were combined for each condition and murine cells were depleted using the Mouse cell depletion kit (Miltenyi Biotec). Afterwards, human cells were subsequently transplanted via intrafemoral injections. Tertiary xenotransplantations were sacrificed at week 12. Engraftment was considered positive if CD45+ engraftment level was >0.1% in the bone marrow.

### Leukemic transformation screen

CMPs and MEPs from T21-FL were sorted by flow cytometry and then combined (as described in fetal liver sorting). Only progenitor cells were utilized in this screen as control or GATA1s CMPs and MEPs did not generate any CD45+ engraftment after 12 weeks post-transplantation, enabling an easy read-out of leukemic transformation. In addition, higher cell numbers can be utilized for each xenograft, thus maintaining a high complexity setting for the CRISPR/Cas9 screen. Progenitor cells were individually CRISPR/Cas9-edited with a single gRNA against a candidate gene and GATA1s (as described in fetal liver sorting and CRISPR/Cas9 RNP electroporation, see Table S6 for gRNA sequences). Four gRNAs per gene were utilized and upon CRISPR/Cas9 editing, cells were pooled according to candidate gene and subsequently transplanted via intrafemoral injections. 2×10^4 progenitor cells were transplanted per mouse and primary xenotransplantations were sacrificed at week 12. Mice were considered leukemic if they showed CD45+ engraftment and blast accumulation (all of the mice that showed CD45+ engraftment were positive for blast accumulation).

### KIT inhibition study

LT-HSCs from T21-FL were sorted by flow cytometry (as described in fetal liver sorting) and CRISPR/Cas9-edited (as described in fetal liver sorting and CRISPR/Cas9 RNP electroporation). 1,300 T21-FL derived LT-HSCs were transplanted per mouse via intrafemoral injections. After 10 weeks, mice were treated twice daily with imatinib (50mg/kg), dasatinib (20mg/kg) and ripretinib (7.5mg/kg) for 2 weeks via oral gavage.

### Real-time quantitative RT-PCR

For GATA1 expression quantification, 1-5×10^4 cells from individual HSPC subpopulations from N-FL and T21-FL were sorted by flow cytometry (as described in fetal liver sorting) and subsequently process using the TaqMan Gene Expression Cells-to-CT Kit according to manufacturer’s protocol. Likewise, for *AGO1, AGO2, TARBP2* and *ADAR* expression quantification, 5×10^5 CD34+ cells from N-FL and T21-FL were sorted, edited for control and GATA1s (as described in CRISPR/Cas9 RNP electroporation) and cultured for 5 days in X-VIVO 10 media (as described above for CRISPR/Cas9 RNP electroporation). qPCR was performed on the Lightcycler 480 instrument II (Roche) using primer/probe sets. All signals were quantified using the ΔCt method and were normalized to the levels of GAPDH. Similarly, for miRNA expression quantification, LT-HSCs from N-FL and T21-FL were sorted by flow cytometry (as described in fetal liver sorting). 2×10^3 LT-HSCs were sorted per sample and subsequently processed using the TaqMan MicroRNA Cells-to-CT kit according to manufacturer’s protocol. qPCR was performed on the Lightcycler 480 instrument II (Roche) using TaqMan primer/probe sets. All signals were quantified using the ΔCt method and were normalized to the levels of RNU48.

### Genomic drop-out analysis by ddPCR

To determine genomic drop-outs of the 50kB region between miR-99a and miR-125b-2 on chromosome 21 upon CRISPR/Cas9 editing of miR-99a and miR-125b-2, droplet digital PCR (ddPCR, Bio-Rad) was carried out. A FAM primer/probe set detecting a 700bp miR-99a-125b-2 fusion and a HEX primer/probe set detecting a 700bp wild-type miR-99a genomic region were designed. Genomic DNA was isolated from CRISPR/Cas9-edited LT-HSCs and subjected to ddPCR (as described in trisomy 21 verification), with the exception that 1μl of genomic DNA (15 and 30ng/μl) and no HaeIII was used. The ddPCR program was: 95°C for 10min, followed by 94°C for 30s, 60°C for 2min (40 cycles) and then 98°C for 10min.

### Virus production and transduction

The chromosome 21 miRNAs miR-99a, miR-125b-2 and miR-155 were overexpressed together with the fluorescent marker mOrange from the SFFV promoter and compared to the empty mOrange control vector. The lentiviral vectors were packaged using 293FT cells (Thermo Fisher). Cells were transfected using the CalPhos Mammalian Transfection Kit (Clontech) according to manufacturer’s protocol. Virus was collected, concentrated using ultracentrifugation and stored at −80°C for subsequent use. Sorted LT-HSCs were cultured for 24 hours in serum-free X-VIVO 10 media (as described above for CRISPR/Cas9 RNP electroporation) and subsequently transduced with lentivirus for 48 hours.

### RNAseq library generation

CRISPR/Cas9-edited cells from primary xenotransplantations were sorted for CD34-CD117+, CD34+CD117+ and CD34+CD117-(as described in setup of secondary xenotransplantation). On average, 1-5×10^4 cells were sorted per sample. Total RNA was purified and DNase treated using the RNeasy Micro kit (Qiagen). RNA integrity was measured using the RNA 6000 Pico kit (Agilent) and showed RNA integrity scores >8. RNAseq libraries were generated using total RNA as input with the SMART-Seq V4 Ultra Low Input RNA kit (Clontech). The cDNA libraries were confirmed on the Bioanalyzer using the High Sensitivity DNA kit (Agilent). Subsequently, next-generation sequencing libraries were generated using the Nextera XT DNA Library Preparation kit (Illumina) and sequenced on the HiSeq 2500 using 125bp paired-end sequencing. Three primary xenograft replicates were used per condition.

### RNAseq and microarray analysis

STAR was used to align raw sequencing reads and to obtain hg38 and count reads over gencode known transcripts. EdgeR was used to normalize read counts and fit a glm model to compute differential expression. Unstranded read counts were used as input for GSEA. Enrichment map in Cytoscape was used to explore GSEA enrichment results. Down syndrome leukemia microarray data was downloaded using GEOquery, quantile normalized and limma was used to calculate differential expression.

### ATACseq library generation

CRISPR/Cas9-edited cells from primary xenotransplantations were sorted for CD34-CD117+, CD34+CD117+ and CD34+CD117- (as described in setup of secondary xenotransplantation). On average, 1-5×10^4 cells were sorted per sample. For constructing the full lineage hierarchy, NFL were processed and sorted (as described in fetal liver sorting), but with the addition of BAH-1 PE (8ml per 1×10^6 cells) in the first staining round and CD71 BV650 (3ml per 1×10^6 cells) in the second staining round. CMPs and MEPs were further divided into F1, F2 and F3 subgroups based on CD71 and BAH-1 expression (F1: BAH-1-CD71-, F2: BAH1-CD71+, F3: BAH1+CD71+) (*24*). On average, 1-10×10^3 cells were sorted per sample. ATACseq libraries were generated as previously described with slight modifications (*70, 71*). For this, cells were lysed in 100μl of lysis buffer (100μl 1M Tris-HCl, 20μl 5M NaCl, 30μl 1M MgCl_2_, 10μl IGEPAL, 0.1% Tween in 10ml H_2_O) for 5min at 4°C. The resulting crude nuclei preparation was centrifuged at 650xg for 10min. This was followed by a transposition reaction using 2.5μl Tagment DNA TDE1 transposase (Illumina), 25μl buffer, 22.5μl nuclease-free H_2_O (IDT) and 0.5μl Tween for 30min at 37°C in a heat block. Following transposition, DNA was purified using the MinElute PCR Purification kit (Qiagen) and eluted in 14μl nuclease-free H2O (IDT). The transposed DNA fragments were subsequently PCR amplified. For this, the number of PCR cycles was pre-determined based on real-time quantitative PCR using 1μl of 1:2 diluted transposed DNA, 1.25μl forward and reverse primer (10μM, IDT), 0.5μl of 1:1000 SYBR Gold (Thermo Fisher), 1μl nuclease-free H2O, 5μl Phusion High-Fidelity master mix (NEB). The sample was run on the CFX96 real-time PCR system (Bio-Rad) with the program: 72°C for 5min, 98°C for 30s, followed by 98°C for 10s, 63°C for 30s and 72°C for 1min (30 cycles). The number of cycles was chosen based on the Ct value at an intensity threshold of 1000. On average, the necessary number of PCR cycles for these samples were between 10-15. Thus, the final PCR amplification was performed using the remaining 12.5μl transposed DNA, 6.25μl forward and barcoded reverse primer (10μM, IDT) and 25μl Phusion High-Fidelity master mix (NEB). The PCR program was: 72°C for 5min, 98°C for 30s, followed by 98°C for 10s, 63°C for 30s and 72°C for 1min (pre-determined number of cycles) and then 72°C for 2.5min. Afterwards, the amplified library was purified using the MinElute PCR Purification kit (Qiagen). Each library was analyzed on the Bioanalyzer using the High Sensitivity DNA kit (Agilent), a primer cleanup was performed using Ampure XP beads (Beckman Coulter) and finally sequenced on the HiSeq 2500 using 50bp paired-end sequencing. Three primary xenograft replicates were used per condition.

### ATACseq analysis

Raw sequencing reads were mapped against the hg38 human reference genome using bwa with default parameters. All duplicate reads, and reads mapped to mitochondria, chromosome Y, an ENCODE blacklisted region or an unspecified contig were removed. MACS was used to call peaks in downsampled reads. The catalogue of all peaks called in any population was produced by merging all called peaks which overlapped by at least one base pair using bedtools. The MACS bdgcmp function was used to compute the fold enrichment over background for all populations, and the bedtools map function was used to identify the maximum fold enrichment observed at each peak in the catalogue in each population. Peaks present in at least two thirds of replicates were compared to identify peaks gained and lost in each contrast. Homer was used to identify enriched motifs within these peaks for each signature, using default parameters, and the catalogue of all called peaks as a background.

### CIBERSORTx analysis

Chromatin accessibility peaks unique to any single N-FL subpopulation sample were determined by comparing the peaks common to all populations of each type. 500bp windows, overlapping by 250bp were identified over these unique peaks, and read counts per sample over these windows were obtained with HTSeq. EdgeR was used to normalize these read counts, and CIBERSORTx was used to create a signature matrix. Counts over windows for each sample to be deconvoluted were analogously obtained and CIBERSORTx was used to infer population proportions.

### Cut&Run Assay

CD34+ cells from N-FL and T21-FL were sorted by flow cytometry (as described in fetal liver sorting) and subsequently 2×10^5 cells were CRISPR/Cas9-edited per condition (as described in fetal liver sorting and CRISPR/Cas9 RNP electroporation). Edited cells were expanded for 5 days in X-VIVO 10 media (as described above for CRISPR/Cas9 RNP electroporation). Cells were washed in cold PBS (Thermo Fisher) and spun down at 350xg for 10min at 4°C. Cut&Run (cleavage under targets and release under nuclease) assay was performed as previously described (*72*). Protein A-Micrococcal nuclease (pA-MN) fusion protein was obtained as a gift from S. Henikoff’s lab. Briefly, cells were washed twice with wash buffer and activated Concanavalin A beads were added dropwise while vortexing the samples. Wash buffer was removed by placing the samples on a magnet and antibody buffer containing 0.0125% of digitonin and GATA1 antibody (Abcam, 1:100) or rabbit gG control antibody (Thermo Fisher, 1:300) was added to the beads. Samples were incubated on a rotator for 2h at 4°C, followed by the addition and activation of pA-MN and isolation of soluble DNA as previously described (*72*). DNA was extracted with the MinElute PCR Purification kit (Qiagen) and DNA libraries were prepared with NEBNext Ultra II DNA Library Prep Kit for Illumina (NEB) and NEBNext Multiplex Oligos for Illumina (NEB). Subsequently, libraries were size selected for 150-400bp range with the Pippen HT and size verified with the Bioanalyzer. Samples were sequenced on Illumina NextSeq500 using 75bp paired-end reads.

### Cut&Run Analysis

Cut&Run paired-end data was trimmed using fastp to remove adapters and low quality base pairs. A base pair quality score of 30 and minimum length cut-off of 35bp were used in the trimming process. Trimmed reads were aligned to the hg38 genome using bowtie2. The same alignment setting was used as previously described (*72*). Unaligned reads and discordantly aligned reads were eliminated and only primary aligned loci were kept. Duplicated reads were kept for the downstream analysis. In order to perform spike-in normalization, the data was aligned to the S. cerevisiae yeast genome (sacCer3) using the same bowtie2 options as previously described (*72*).

Peaks were called using MACS2. Instead of normalizing the samples by the library size, they were normalized by the number of spikein reads that came from aligning the samples to the yeast genome. Peaks from different samples were intersected in a pairwise manner to determine the shared and specific peaks between two samples. ChipPeakAnnot library in the R package was used to determine the overlap between peak sets and determine the shared and specific peaks. Motif analysis was performed on called peaks using homer. Normalized enrichment score (NES) for each motif was calculated as the fold change of the target to the background percentage. If the target percentage is less than 5, the pseudo count 1 was added to the target and the background percentages before calculating the fold change to attenuate the fold change of motifs with low target percentages. Peaks were annotated to determine the overlap with the genomic features such as promoters, 5’UTR, 3’UTR, introns, exons and intergenic regions using the Refseq genome annotation. Promoters were defined as the genomic regions starting 2Kb upstream of the transcriptional start site until 500bp downstream of the transcriptional start site. Gene ontology analysis was performed on the list of genes that have peaks bound to their promoters. PANTHER was used for gene ontology analysis.

### miRNA profiling

CD34+ cells from N-FL and T21-FL were sorted by flow cytometry (as described in fetal liver sorting). Total RNA was isolated from sorted 2-3×10^5 CD34+ cells using the mirVANA miRNA isolation kit (Thermo Fisher) and subsequently processed using the Small RNA Library Prep kit (Norgen Biotek). Subsequently, libraries were size excised at 130-177bp using the Pippin Prep and sequenced on the Illumina HiSeq2500 using single-end reads of 50bp.

### miRNA profiling analysis

The fastq files were processed using a pipeline previously described (*73*). Briefly, the fastq files were trimmed using cutadapt and sequences below 15 nucleotides were removed followed by annotation and alignment using bwa and a custom perl script attached to the pipeline accessible from github. Minor modifications to ensure compatibility with perl5 and MariaDB were made. miRNA v.22b was used as reference from miRbase and the alignment was conducted to hg38 as accessed from UCSC. Count data was then analyzed in edgeR using standard parameters.

### Statistical analysis

Statistical significance was calculated using two-tailed unpaired student’s t-test. For motif enrichment analyses, cumulative binomial tests were used to calculate p-values and for GO term analyses binomial tests were used. Sample size was chosen to give sufficient power for calling significance with standard statistical tests.

## Materials table

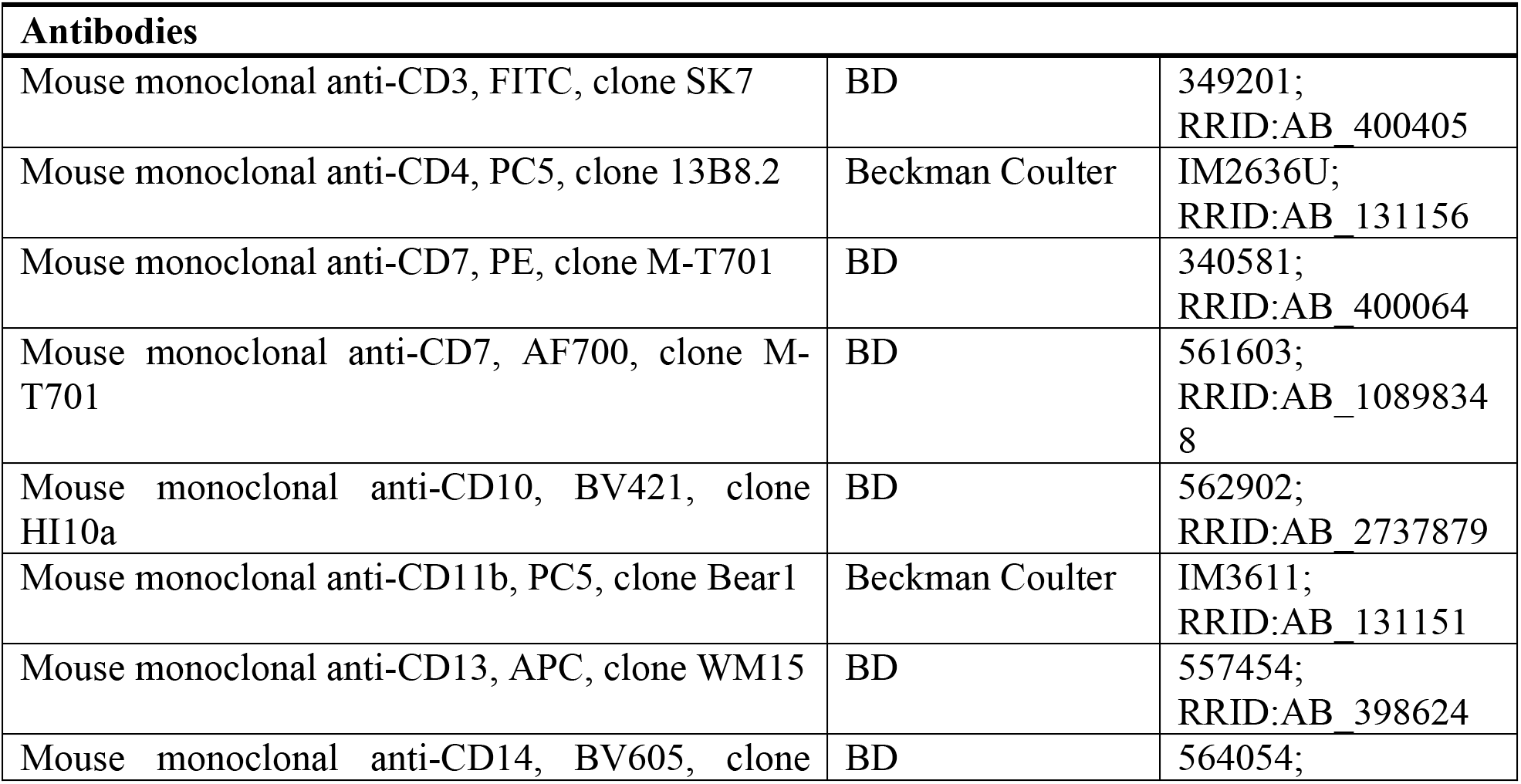
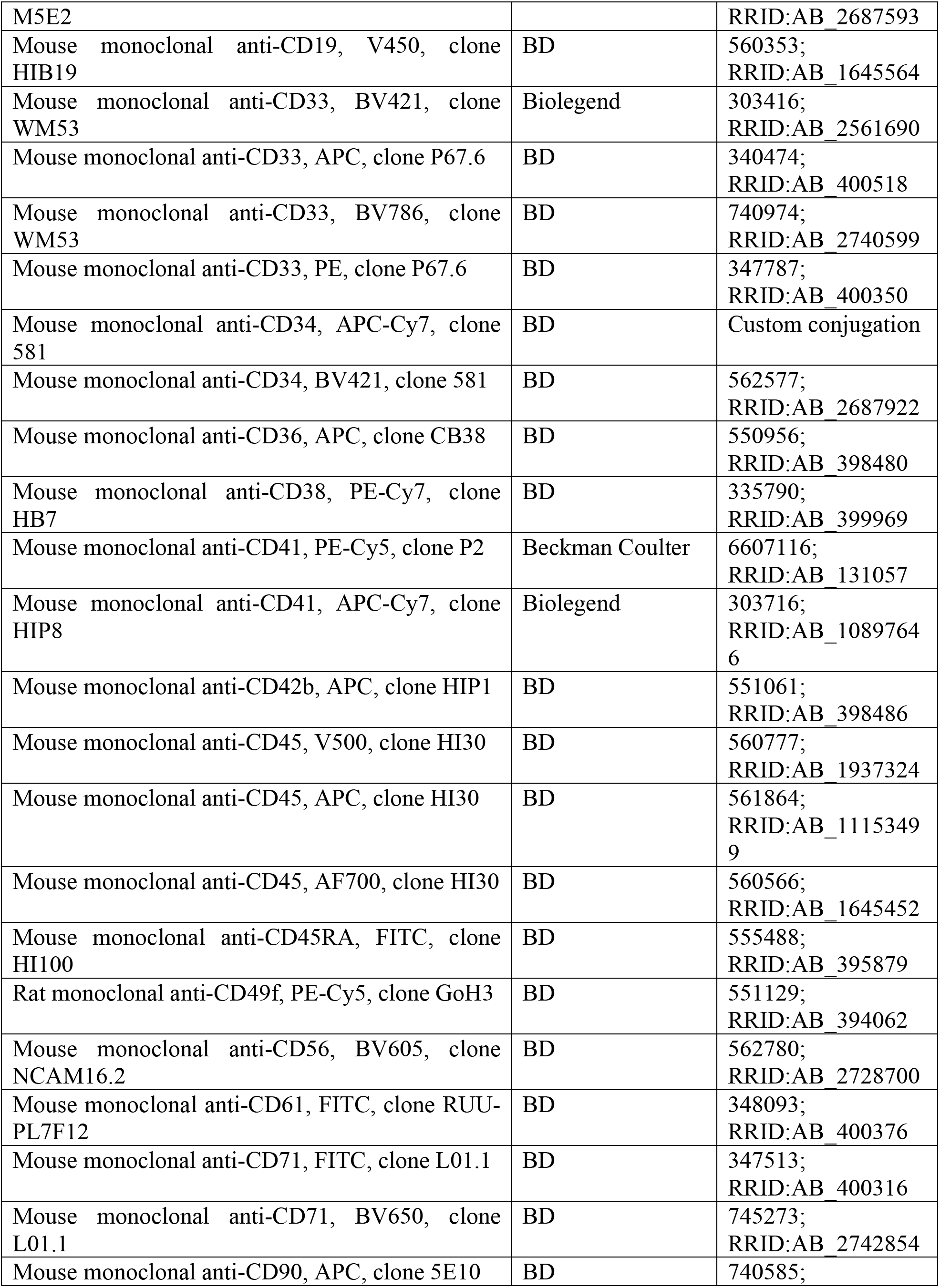
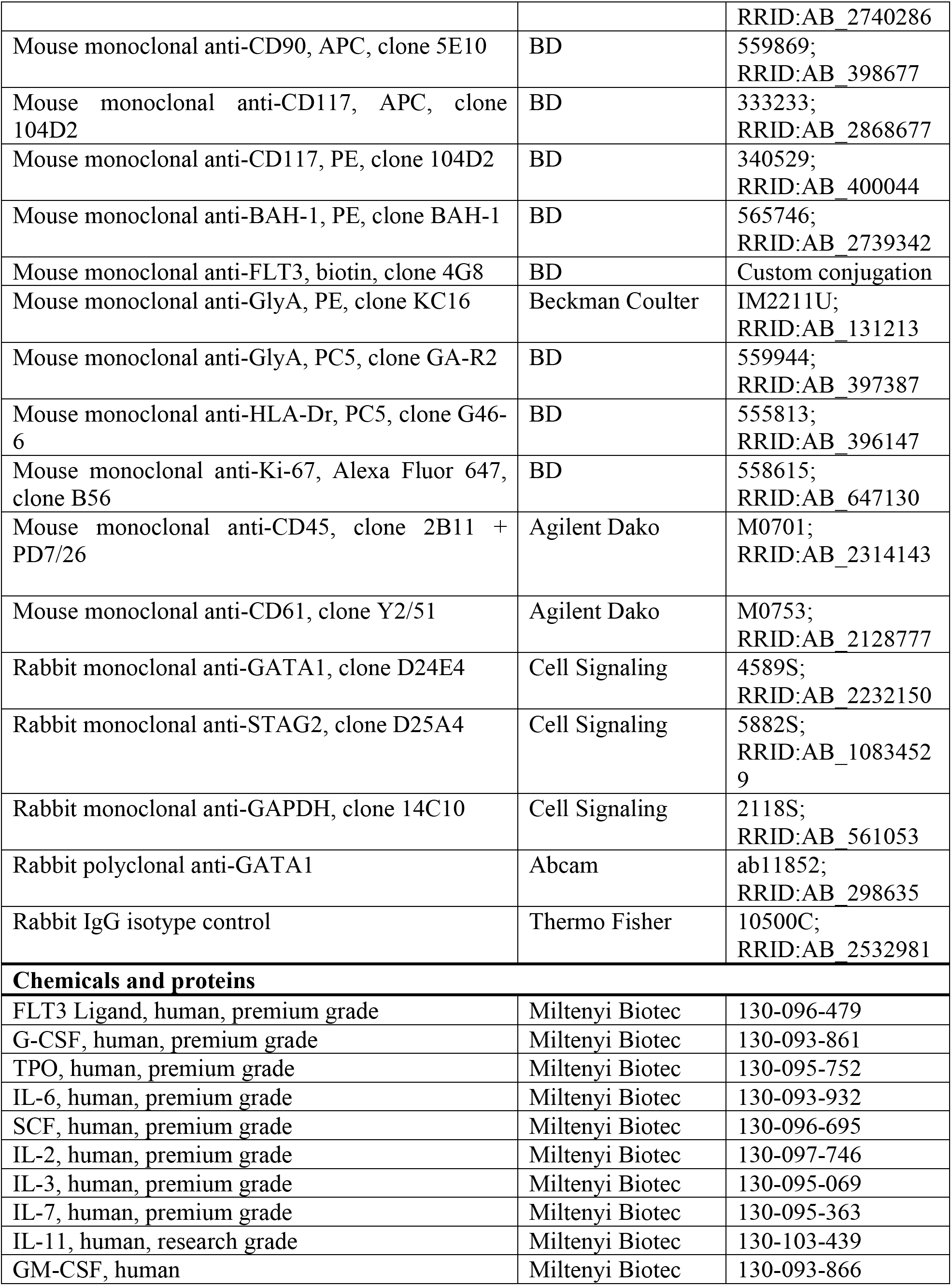
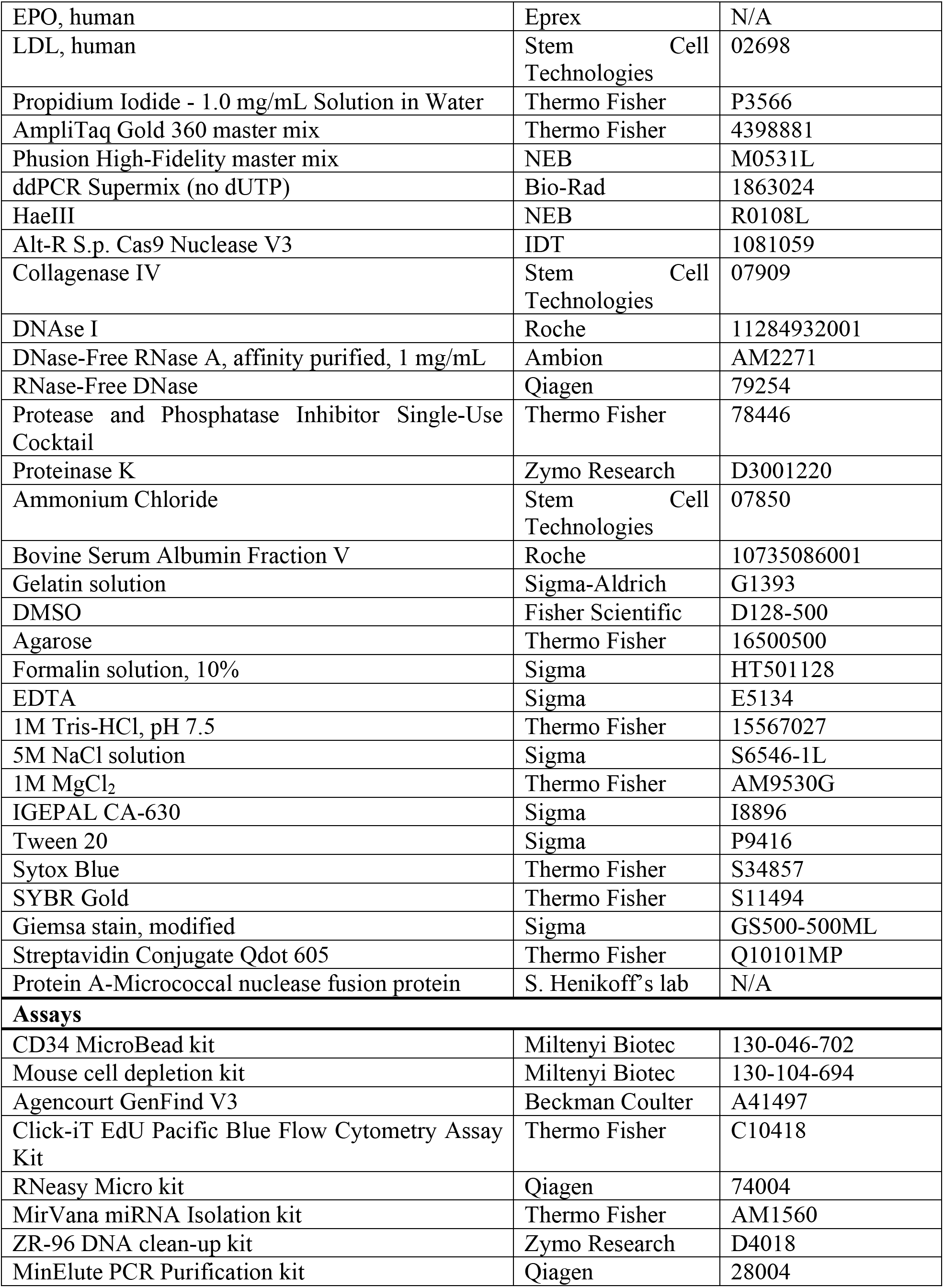
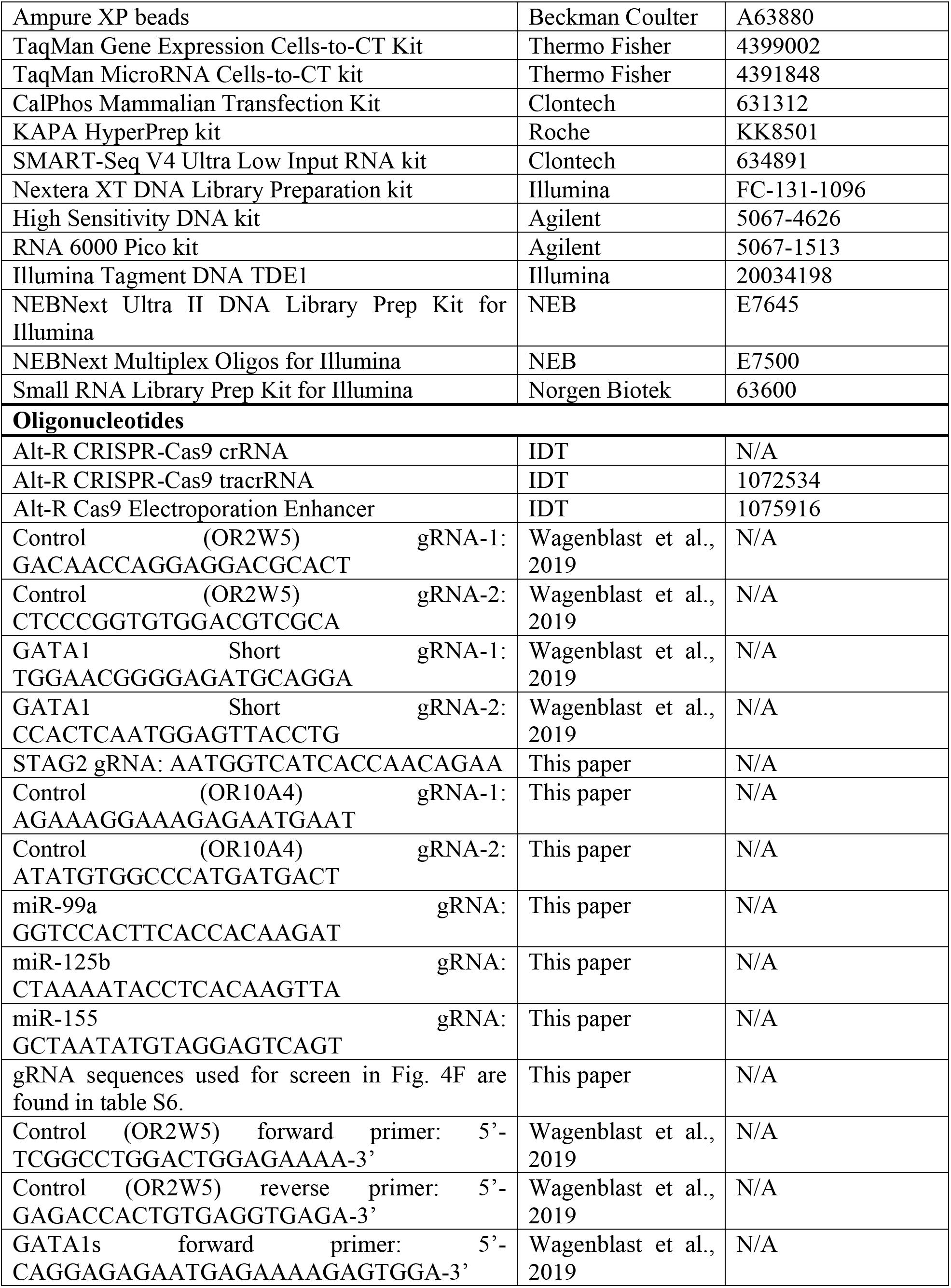
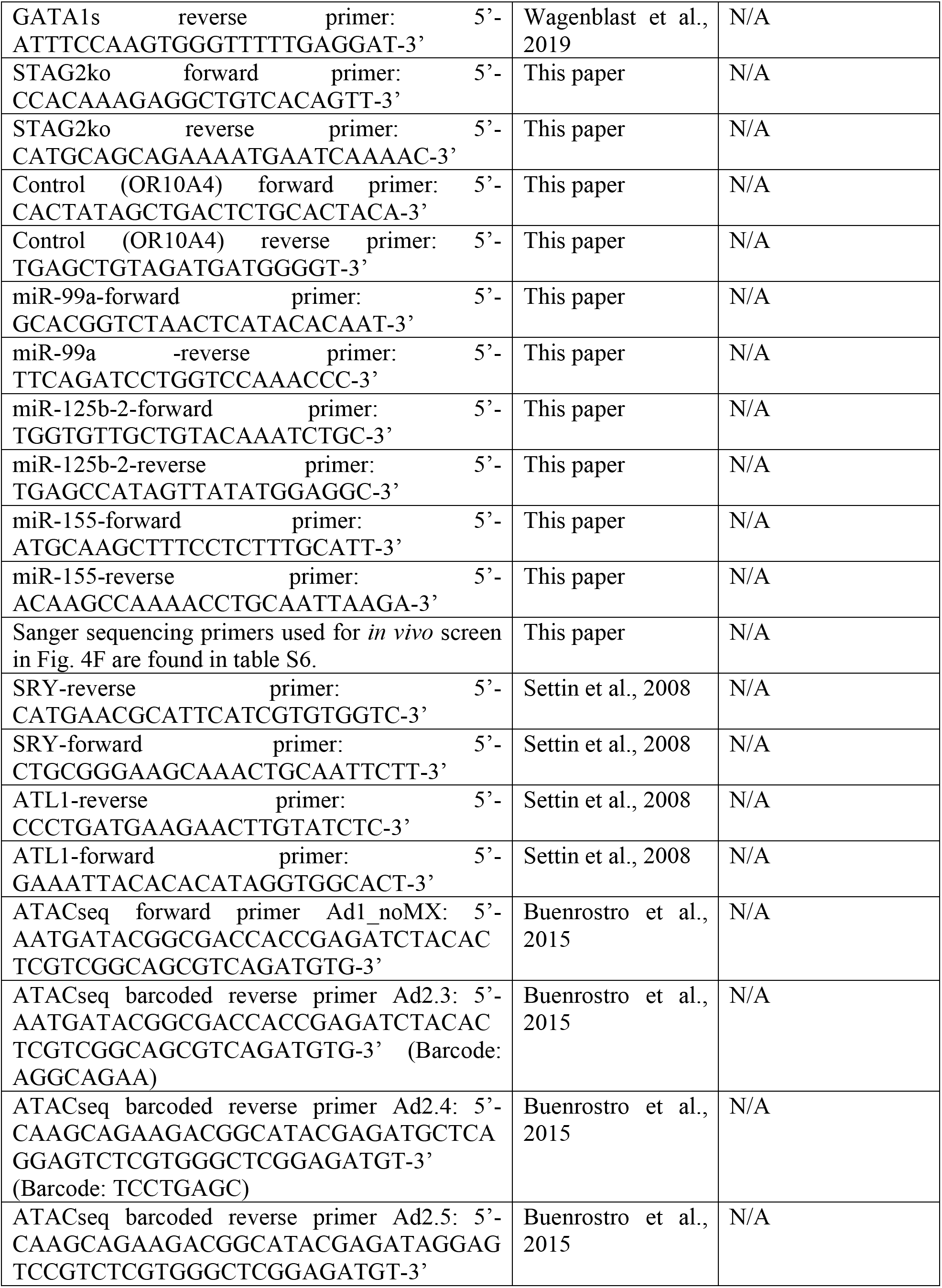
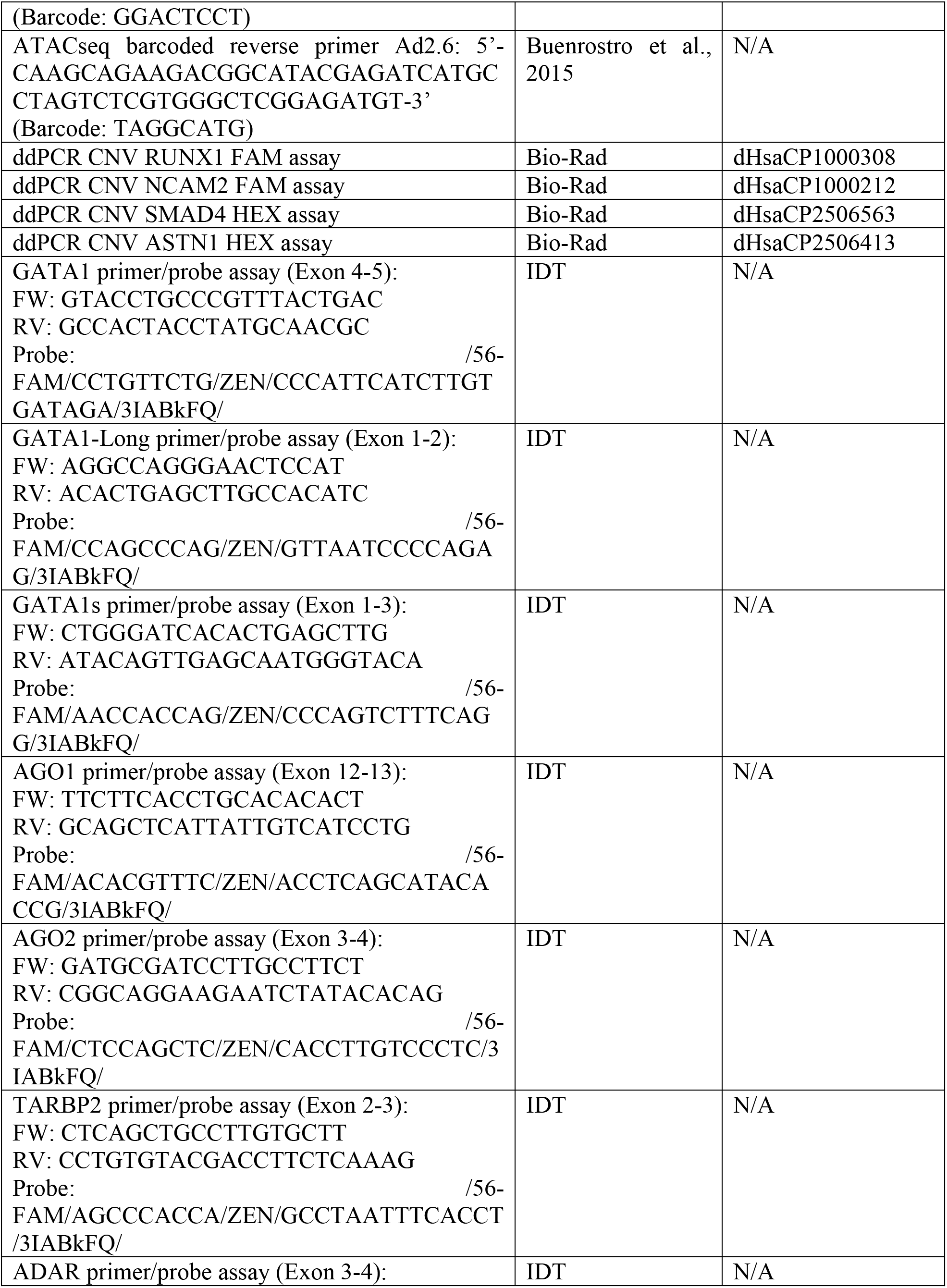
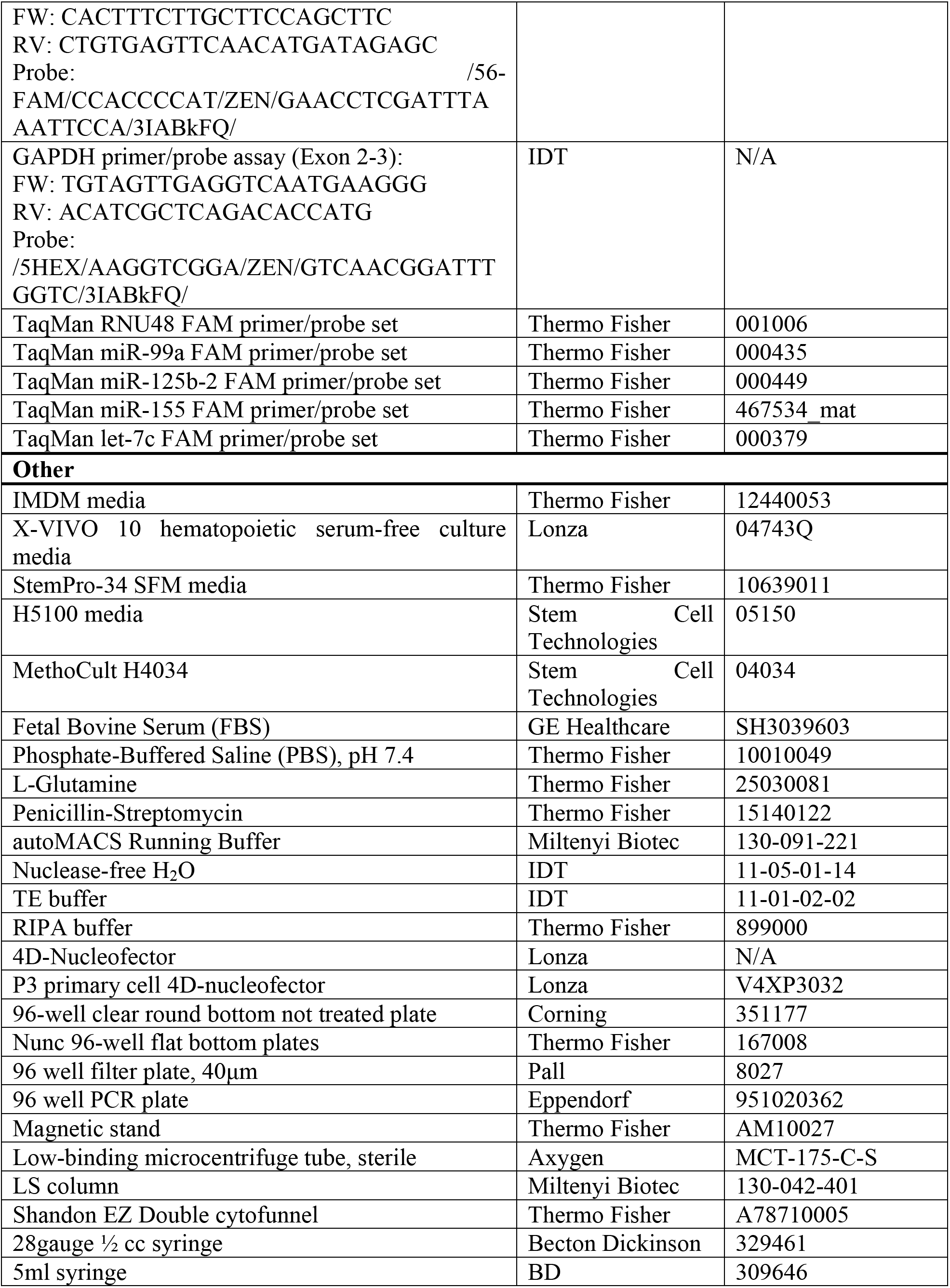
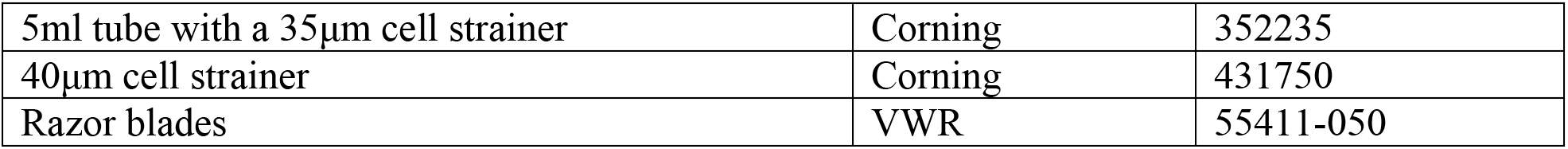

## References and Notes

1. H. Hasle, I. H. Clemmensen, M. Mikkelsen, Risks of leukaemia and solid tumours in individuals with Down’s syndrome. Lancet. 355, 165–169 (2000).

2. M. F. Greaves, A. T. Maia, J. L. Wiemels, A. M. Ford, Leukemia in twins: lessons in natural history. Blood. 102, 2321–2333 (2003).

3. J. K. Hitzler, J. Cheung, Y. Li, S. W. Scherer, A. Zipursky, GATA1 mutations in transient leukemia and acute megakaryoblastic leukemia of Down syndrome. Blood. 101, 4301–4304 (2003).

4. I. Roberts et al., GATA1-mutant clones are frequent and often unsuspected in babies with Down syndrome: identification of a population at risk of leukemia. Blood. 122, 3908–3917 (2013).

5. J. Wechsler et al., Acquired mutations in GATA1 in the megakaryoblastic leukemia of Down syndrome. Nat. Genet. 32, 148–152 (2002).

6. S. Hoeller et al., Morphologic and GATA1 sequencing analysis of hematopoiesis in fetuses with trisomy 21. Hum Pathol. 45, 1003–1009 (2014).

7. M. Uffmann et al., Therapy reduction in patients with Down syndrome and myeloid leukemia: the international ML-DS 2006 trial. Blood. 129, 3314–3321 (2017).

8. J. W. Taub et al., Improved outcomes for myeloid leukemia of Down syndrome: a report from the Children’s Oncology Group AAML0431 trial. Blood. 129, 3304–3313 (2017).

9. A. S. Gamis et al., Natural history of transient myeloproliferative disorder clinically diagnosed in Down syndrome neonates: a report from the Children’s Oncology Group Study A2971. Blood. 118, 6752–9– quiz 6996 (2011).

10. M. Labuhn et al., Mechanisms of Progression of Myeloid Preleukemia to Transformed Myeloid Leukemia in Children with Down Syndrome. Cancer Cell (2019), doi:10.1016/j.ccell.2019.06.007.

11. K. Yoshida et al., The landscape of somatic mutations in Down syndrome-related myeloid disorders. Nat. Genet. 45, 1293–1299 (2013).

12. J.-H. Klusmann et al., Treatment and prognostic impact of transient leukemia in neonates with Down syndrome. Blood. 111, 2991–2998 (2008).

13. A. D. Sorrell et al., Favorable survival maintained in children who have myeloid leukemia associated with Down syndrome using reduced-dose chemotherapy on Children”s Oncology Group trial A2971: a report from the Children”s Oncology Group. Cancer. 118, 4806–4814 (2012).

14. M. Flasinski et al., Low-dose cytarabine to prevent myeloid leukemia in children with Down syndrome: TMD Prevention 2007 study. Blood Adv. 2, 1532–1540 (2018).

15. T. Taga et al., Clinical characteristics and outcome of refractory/relapsed myeloid leukemia in children with Down syndrome. Blood. 120, 1810–1815 (2012).

16. J. K. Hitzler et al., Outcome of transplantation for acute myelogenous leukemia in children with Down syndrome. Biol Blood Marrow Transplant. 19, 893–897 (2013).

17. J. T. Caldwell, Y. Ge, J. W. Taub, Prognosis and management of acute myeloid leukemia in patients with Down syndrome. Expert Rev Hematol. 7, 831–840 (2014).

18. K. A. Alford et al., Perturbed hematopoiesis in the Tc1 mouse model of Down syndrome. Blood. 115, 2928–2937 (2010).

19. G. Kirsammer et al., Highly penetrant myeloproliferative disease in the Ts65Dn mouse model of Down syndrome. Blood. 111, 767–775 (2008).

20. S. Malinge et al., Increased dosage of the chromosome 21 ortholog Dyrk1a promotes megakaryoblastic leukemia in a murine model of Down syndrome. J. Clin. Invest. 122, 948–962 (2012).

21. E. Wagenblast et al., Functional profiling of single CRISPR/Cas9-edited human long-term hematopoietic stem cells. Nature Communications. 10, 4730–11 (2019).

22. F. Notta et al., Evolution of human BCR-ABL1 lymphoblastic leukaemia-initiating cells. Nature. 469, 362–367 (2011).

23. A. Roy et al., Perturbation of fetal liver hematopoietic stem and progenitor cell development by trisomy 21. Proc. Natl. Acad. Sci. U.S.A. 109, 17579–17584 (2012).

24. F. Notta et al., Distinct routes of lineage development reshape the human blood hierarchy across ontogeny. Science. 351, aab2116–aab2116 (2016).

25. C. Langebrake, U. Creutzig, D. Reinhardt, Immunophenotype of Down syndrome acute myeloid leukemia and transient myeloproliferative disease differs significantly from other diseases with morphologically identical or similar blasts. Klin Padiatr. 217, 126–134 (2005).

26. L. Wang et al., Acute megakaryoblastic leukemia associated with trisomy 21 demonstrates a distinct immunophenotype. Cytometry B Clin Cytom. 88, 244–252 (2015).

27. WHO, WHO Classification of Tumours of Haematopoietic and Lymphoid Tissues (World Health Organization, 2008).

28. N. Bhatnagar, L. Nizery, O. Tunstall, P. Vyas, I. Roberts, Transient Abnormal Myelopoiesis and AML in Down Syndrome: an Update. Curr Hematol Malig Rep. 11, 333–341 (2016).

29. A. D. Viny et al., Cohesin Members Stag1 and Stag2 Display Distinct Roles in Chromatin Accessibility and Topological Control of HSC Self-Renewal and Differentiation. Cell Stem Cell. 25, 682–696.e8 (2019).

30. A. M. Newman et al., Determining cell type abundance and expression from bulk tissues with digital cytometry. Nat. Biotechnol. 37, 773–782 (2019).

31. A. Schwarzer et al., The non-coding RNA landscape of human hematopoiesis and leukemia. Nature Communications. 8, 218–17 (2017).

32. A. V. Krivtsov et al., Transformation from committed progenitor to leukaemia stem cell initiated by MLL-AF9. Nature. 442, 818–822 (2006).

33. Y. Wang et al., The Wnt/beta-catenin pathway is required for the development of leukemia stem cells in AML. Science. 327, 1650–1653 (2010).

34. M. Ye et al., Hematopoietic Differentiation Is Required for Initiation of Acute Myeloid Leukemia. Cell Stem Cell. 17, 611–623 (2015).

35. P. J. Skene, S. Henikoff, An efficient targeted nuclease strategy for high-resolution mapping of DNA binding sites. Elife. 6, 576 (2017).

36. T. M. Chlon, M. McNulty, B. Goldenson, A. Rosinski, J. D. Crispino, Global transcriptome and chromatin occupancy analysis reveal the short isoform of GATA1 is deficient for erythroid specification and gene expression. Haematologica. 100, 575–584 (2015).

37. J. Domen, I. L. Weissman, Hematopoietic stem cells need two signals to prevent apoptosis; BCL-2 can provide one of these, Kitl/c-Kit signaling the other. J. Exp. Med. 192, 1707–1718 (2000).

38. H. Kantarjian et al., Dasatinib versus imatinib in newly diagnosed chronic-phase chronic myeloid leukemia. N. Engl. J. Med. 362, 2260–2270 (2010).

39. B. D. Smith et al., Ripretinib (DCC-2618) Is a Switch Control Kinase Inhibitor of a Broad Spectrum of Oncogenic and Drug-Resistant KIT and PDGFRA Variants. Cancer Cell. 35, 738–751. e9 (2019).

40. M. C. Heinrich et al., Inhibition of c-kit receptor tyrosine kinase activity by STI 571, a selective tyrosine kinase inhibitor. Blood. 96, 925–932 (2000).

41. S. T. Chou et al., Trisomy 21 enhances human fetal erythro-megakaryocytic development. Blood. 112, 4503–4506 (2008).

42. O. Tunstall-Pedoe et al., Abnormalities in the myeloid progenitor compartment in Down syndrome fetal liver precede acquisition of GATA1 mutations. Blood. 112, 4507–4511 (2008).

43. S. McLean, C. McHale, H. Enright, Hematological abnormalities in adult patients with Down’s syndrome. Ir J Med Sci. 178, 35–38 (2009).

44. B. Liu, S. Filippi, A. Roy, I. Roberts, Stem and progenitor cell dysfunction in human trisomies. EMBO Rep. 16, 44–62 (2015).

45. L. Gutiérrez et al., Ablation of Gata1 in adult mice results in aplastic crisis, revealing its essential role in steady-state and stress erythropoiesis. Blood. 111, 4375–4385 (2008).

46. I. Roberts, S. Izraeli, Haematopoietic development and leukaemia in Down syndrome. Br. J. Haematol. 167, 587–599 (2014).

47. E. R. Lechman et al., miR-126 Regulates Distinct Self-Renewal Outcomes in Normal and Malignant Hematopoietic Stem Cells. Cancer Cell. 29, 214–228 (2016).

48. J.-H. Klusmann et al., miR-125b-2 is a potential oncomiR on human chromosome 21 in megakaryoblastic leukemia. Genes Dev. 24, 478–490 (2010).

49. S. W. K. Ng et al., A 17-gene stemness score for rapid determination of risk in acute leukaemia. Nature. 540, 433–437 (2016).

50. D. Cruz Hernandez et al., Sensitive, rapid diagnostic test for transient abnormal myelopoiesis and myeloid leukemia of Down syndrome. Blood. 136, 1460–1465 (2020).

51. C. M. McHale et al., Prenatal origin of childhood acute myeloid leukemias harboring chromosomal rearrangements t(15;17) and inv(16). Blood. 101, 4640–4641 (2003).

52. C. M. McHale et al., Prenatal origin of TEL-AML1-positive acute lymphoblastic leukemia in children born in California. Genes Chromosom. Cancer. 37, 36–43 (2003).

53. J. L. Wiemels et al., In utero origin of t(8;21) AML1-ETO translocations in childhood acute myeloid leukemia. Blood. 99, 3801–3805 (2002).

54. Y. Wang et al., Impact of age on the survival of pediatric leukemia: an analysis of 15083 children in the SEER database. Oncotarget. 7, 83767–83774 (2016).

55. P. H. G. Duijf, N. Schultz, R. Benezra, Cancer cells preferentially lose small chromosomes. Int. J. Cancer. 132, 2316–2326 (2013).

56. A. P. Laurent, R. S. Kotecha, S. Malinge, Gain of chromosome 21 in hematological malignancies: lessons from studying leukemia in children with Down syndrome. Leukemia. 34, 1984–1999 (2020).

57. A. Settin, E. Elsobky, A. Hammad, A. Al-Erany, Rapid sex determination using PCR technique compared to classic cytogenetics. Int J Health Sci Qassim). 2, 49–52 (2008).

58. N. Erard, S. R. V. Knott, G. J. Hannon, A CRISPR Resource for Individual, Combinatorial, or Multiplexed Gene Knockout. Mol. Cell. 67, 348–354. e4 (2017).

59. M. Aregger, M. Chandrashekhar, A. H. Y. Tong, K. Chan, J. Moffat, Pooled Lentiviral CRISPR-Cas9 Screens for Functional Genomics in Mammalian Cells. Methods Mol Biol. 1869, 169–188 (2019).

60. T. Hart et al., High-Resolution CRISPR Screens Reveal Fitness Genes and Genotype-Specific Cancer Liabilities. Cell. 163, 1515–1526 (2015).

61. S. F. Altschul, W. Gish, W. Miller, E. W. Myers, D. J. Lipman, Basic local alignment search tool. . Mol. Biol 215, 403–410 (1990).

62. H. Li, R. Durbin, Fast and accurate short read alignment with Burrows-Wheeler transform. Bioinformatics. 25, 1754–1760 (2009).

63. K. Itoh et al., Reproducible establishment of hemopoietic supportive stromal cell lines from murine bone marrow. Exp. Hematol. 17, 145–153 (1989).

64. F. Mazurier, M. Doedens, O. I. Gan, J. E. Dick, Rapid myeloerythroid repopulation after intrafemoral transplantation of NOD-SCID mice reveals a new class of human stem cells. Nat. Med. 9, 959–963 (2003).

65. E. K. Brinkman, T. Chen, M. Amendola, B. van Steensel, Easy quantitative assessment of genome editing by sequence trace decomposition. Nucleic Acids Res. 42, e168–e168 (2014).

66. Y. Hu, G. K. Smyth, ELDA: extreme limiting dilution analysis for comparing depleted and enriched populations in stem cell and other assays. J. Immunol. Methods. 347, 70–78 (2009).

67. P. Bankhead et al., QuPath: Open source software for digital pathology image analysis. Sci Rep. 7, 16878–7 (2017).

68. C. A. Schneider, W. S. Rasband, K. W. Eliceiri, NIH Image to ImageJ: 25 years of image analysis. Nat. Methods. 9, 671–675 (2012).

69. A. C. Ruifrok, D. A. Johnston, Quantification of histochemical staining by color deconvolution. Anal. Quant. Cytol. Histol. 23, 291–299 (2001).

70. J. D. Buenrostro, B. Wu, H. Y. Chang, W. J. Greenleaf, ATAC-seq: A Method for Assaying Chromatin Accessibility Genome-Wide. Curr Protoc Mol Biol. 109, 21.29.1–21.29.9 (2015).

71. M. R. Corces et al., An improved ATAC-seq protocol reduces background and enables interrogation of frozen tissues. Nat. Methods. 14, 959–962 (2017).

72. P. J. Skene, J. G. Henikoff, S. Henikoff, Targeted in situ genome-wide profiling with high efficiency for low cell numbers. Nat Protoc. 13, 1006–1019 (2018).

73. A. Chu et al., Large-scale profiling of microRNAs for The Cancer Genome Atlas. Nucleic Acids Res. 44, e3–e3 (2016).

