## Supplementary Figures 1-10 for "Mapping the Cellular Origin and Early Evolution of Leukemia in Down Syndrome"

Wagenblast et al., Supplementary Figure 1

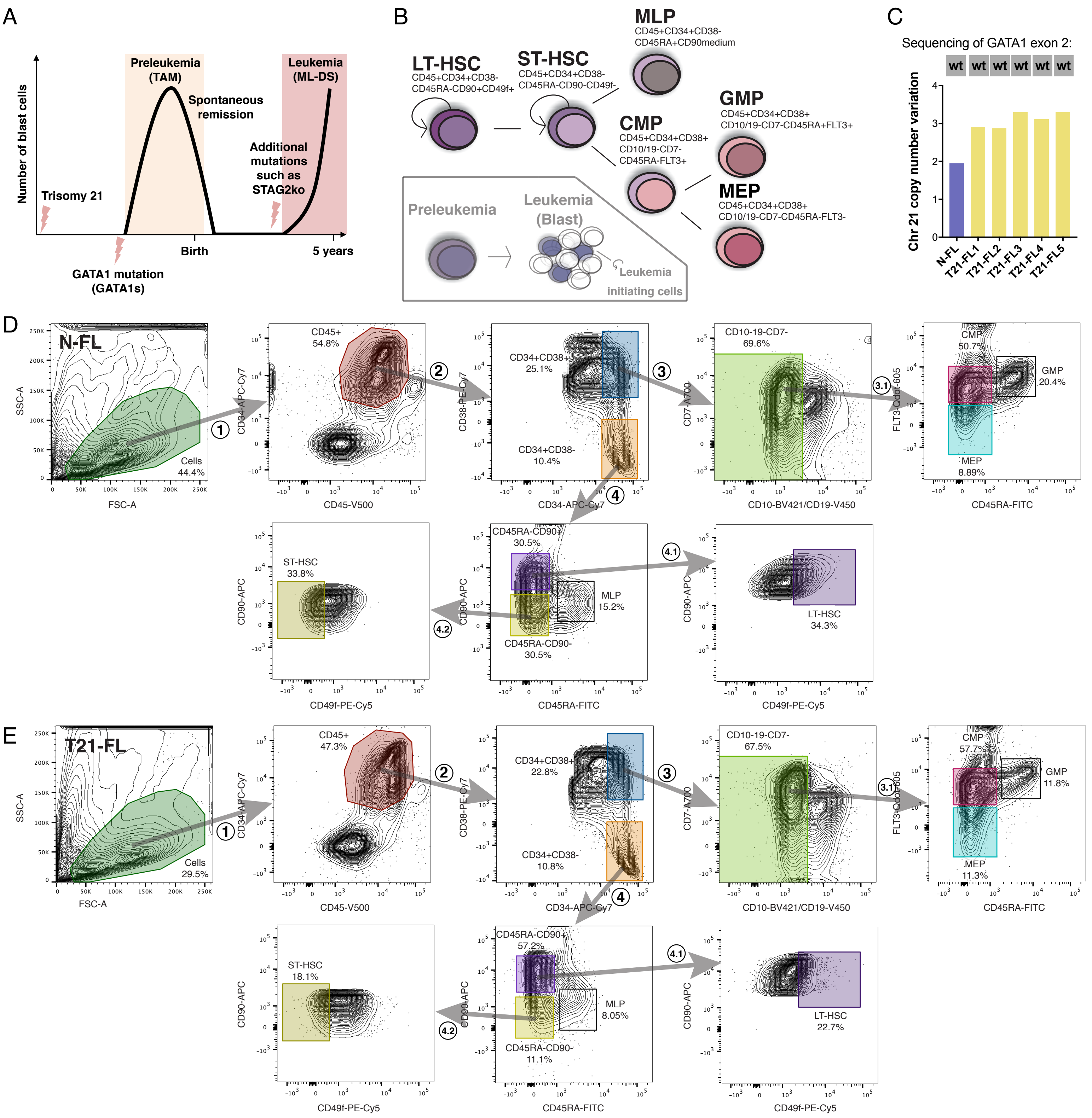


**Fig. S1 Lineage hierarchy of normal disomic and trisomic 21 fetal liver hematopoietic stem cells.**
**(A)** Schematic overview of disease initiation and progression of Down syndrome preleukemia and leukemia. **(B)** Schematic of stem and progenitor lineage hierarchy of human hematopoietic stem cells, which are defined by cell surface markers. On the malignant side, preleukemic cells represent evolutionary ancestors of leukemia. **(C)** Copy number variation of chromosome 21 of all five T21-FL samples used in this study, as determined by droplet digital PCR. **(D)** Flow cytometry sorting scheme of N-FL. Fetal liver-derived CD34+ enriched stem and progenitor cells were utilized to sort LT-HSCs as CD45+CD34+CD38–CD45RA–CD90+CD49f+, ST-HSCs as CD45+CD34+CD38–CD45RA–CD90–CD49f–, CMPs as CD45+CD34+CD38+CD10–CD19–CD7–CD45RA–FLT3+ and MEPs as CD45+CD34+CD38+CD10–CD19–CD7–CD45RA–FLT3–. **(E)** Flow cytometry sorting scheme of T21-FL. Gates are based on normal disomic fetal liver in (D).

Wagenblast et al., Supplementary Figure 2

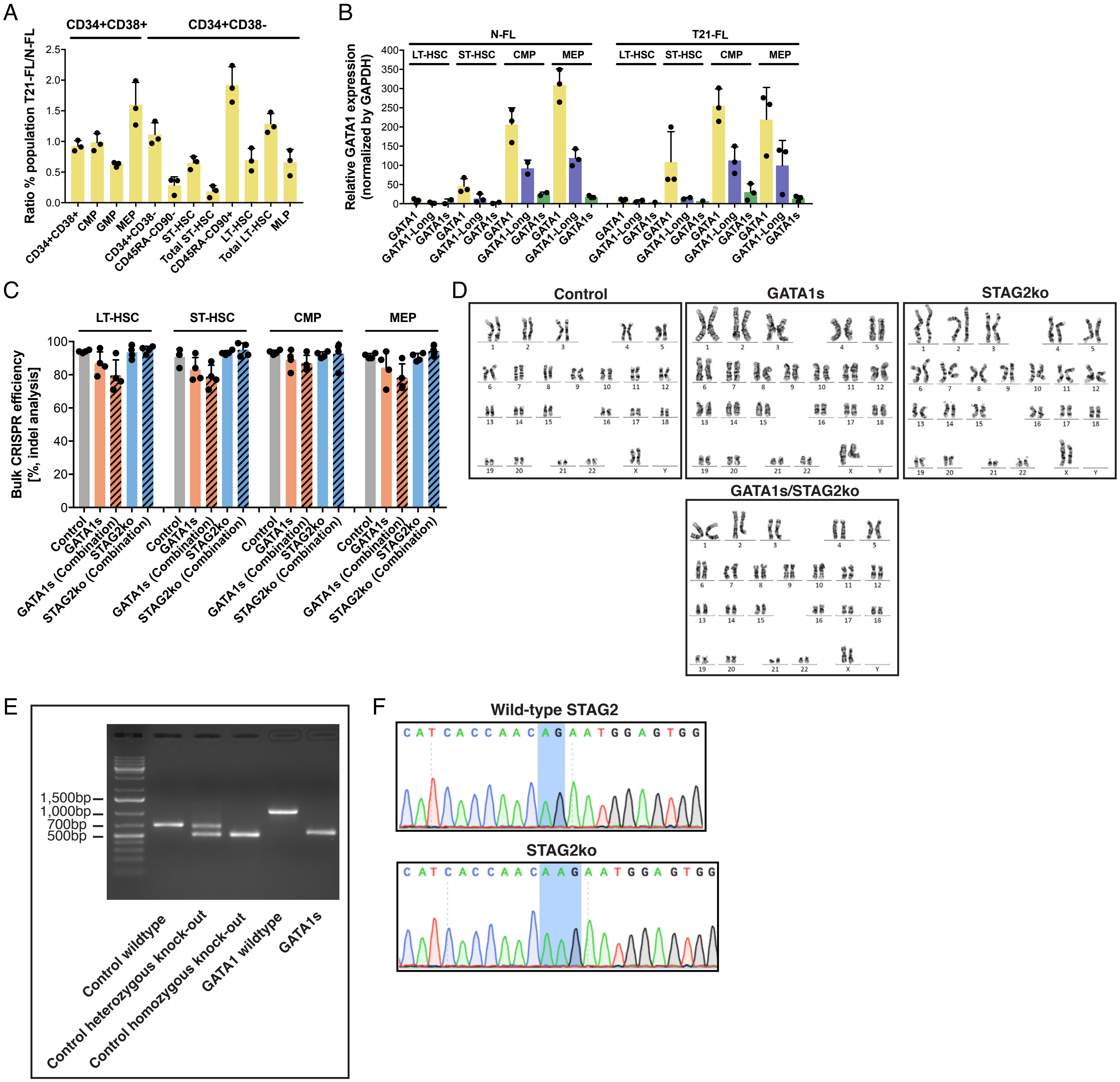


**Fig. S2 Lineage hierarchy and CRISPR/Cas9-editing in fetal liver hematopoietic stem cells.**
**(A)** Percent ratio of HSPC subpopulations between N-FL and T21-FL using the same gates as in fig. S1, D and E. (n = 3 replicates per condition). **(B)** Relative expression of total GATA1, the long isoform of GATA1 (GATA1-Long) and GATA1s in N-FL and T21-FL HSPC subpopulations as determined by RT-qPCR (n = 2-3 replicates per HSPC subpopulation). **(C)** CRISPR/Cas9 efficiency in bulk HSPC subpopulations, as determined by Sanger sequencing and indel analysis. Of note, this analysis does not discriminate between cases where only a single gRNA or both gRNAs cut in control or GATA1s (n = 4 replicates per condition). **(D)** G-banding karyotype analysis on N-FL CD34+CD38– enriched stem cells (n = 20 metaphases per condition). **(E)** Gel electrophoresis analysis of single cell-derived *in vitro* colonies that were CRISPR/Cas9-edited with control and GATA1s. Successfully edited colonies display a shift in size of the PCR product. For the single cell *in vitro* assays, only male samples were used since GATA1 and STAG2 are located on the X-chromosome, resulting in only one allele needing to be edited (total numbers of successfully edited control and GATA1s colonies that were used for analysis were 802 and 1407, respectively). **(F)** Sanger sequencing analysis of a single cell derived colony that was CRISPR/Cas9-edited with STAG2ko (total number of successfully edited STAG2ko colonies that was used for analysis was 946). Error bars represent standard deviation.

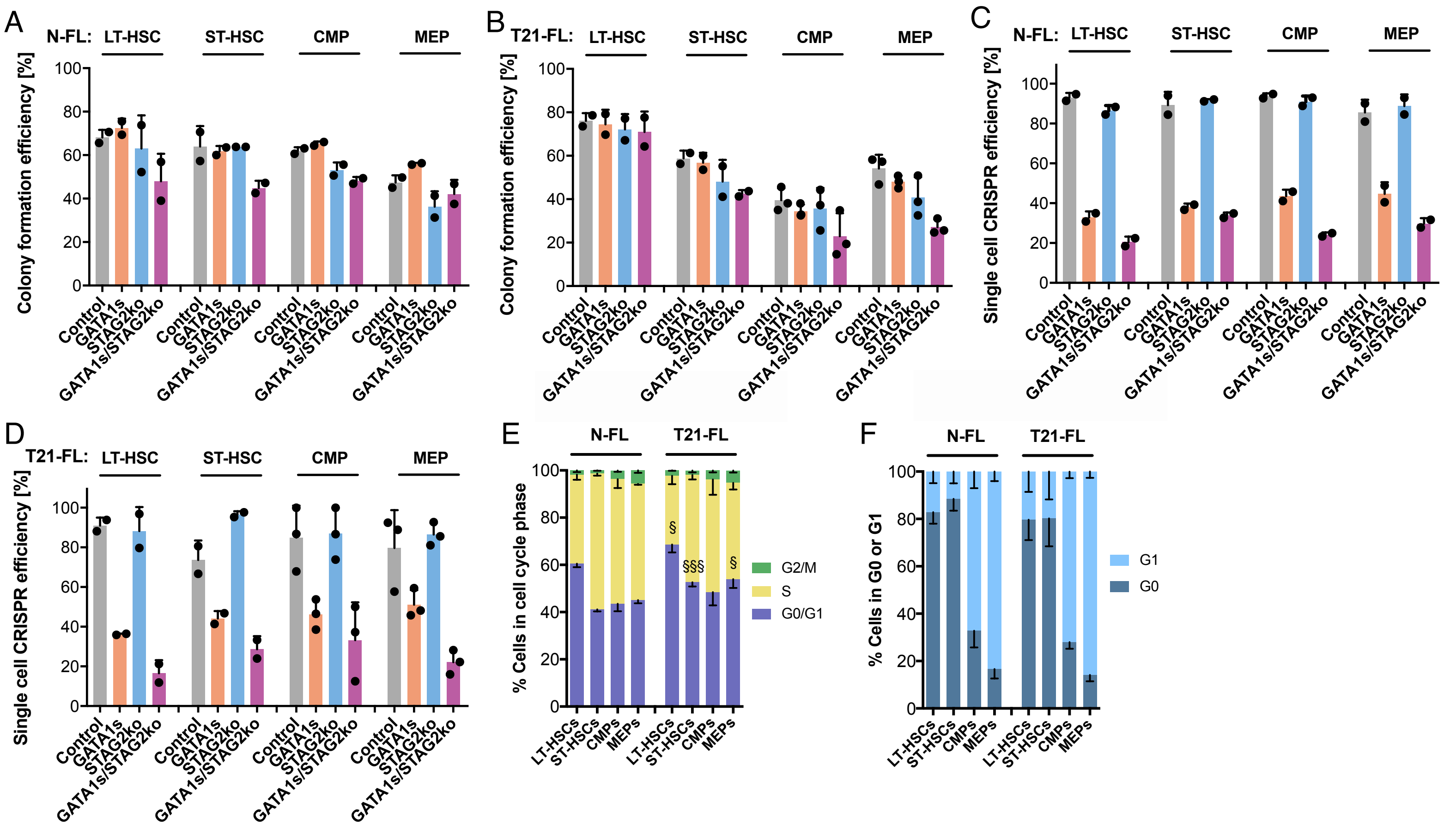


G

| N-FL | | % Colonies containing (relative to control) | | | |
|---|---|---|---|---|---|
| | | Meg | CD71 | E | M |
| LT-HSC | GATA1s | +6.3 | +11.7 | -29.1 | +0.4 |
| | STAG2ko | +2.5 | +0.0 | +10.9 | -2.2 |
| | GATA1s/STAG2ko | +37.1 | +3.8 | -31.2 | +0.4 |
| ST-HSC | GATA1s | +11.1 | +3.2 | -16.6 | +0.0 |
| | STAG2ko | +6.9 | +0.0 | +20.0 | -0.8 |
| | GATA1s/STAG2ko | +18.4 | +10.5 | -14.1 | +0.0 |
| CMP | GATA1s | +20.6 | +7.3 | -15.8 | +0.0 |
| | STAG2ko | +4.7 | +0.0 | +20.6 | -1.6 |
| | GATA1s/STAG2ko | +12.7 | +20.7 | -15.9 | +0.0 |
| MEP | GATA1s | +17.4 | +2.9 | -10.0 | +7.2 |
| | STAG2ko | +1.4 | +0.0 | +28.5 | -13.4 |
| | GATA1s/STAG2ko | +6.7 | +10.0 | -5.1 | +0.5 |

H

| T21-FL | | % Colonies containing (relative to control) | | | |
|---|---|---|---|---|---|
| | | Meg | CD71 | E | M |
| LT-HSC | GATA1-Short | +13.7 | +2.9 | -18.4 | +3.2 |
| | STAG2ko | +4.5 | +0.0 | +8.4 | +3.2 |
| | GATA1s/STAG2ko | +6.8 | +1.8 | -26.6 | +3.2 |
| ST-HSC | GATA1-Short | +16.5 | +13.3 | -29.0 | +18.4 |
| | STAG2ko | -1.3 | +0.0 | +21.4 | -29.2 |
| | GATA1s/STAG2ko | +7.8 | +9.1 | -26.1 | +13.9 |
| CMP | GATA1-Short | +17.8 | +2.0 | -9.2 | +29.9 |
| | STAG2ko | -2.6 | +0.0 | +23.5 | -6.2 |
| | GATA1s/STAG2ko | +6.5 | +6.1 | -21.9 | +31.0 |
| MEP | GATA1-Short | +13.9 | +1.4 | -16.9 | +38.4 |
| | STAG2ko | +0.0 | +0.0 | +19.4 | -15.0 |
| | GATA1s/STAG2ko | +15.2 | +0.0 | -19.8 | +29.1 |

I

| NSG | LT-HSC dose | 1° transplantation engrafted/injected | Stem cell frequency |
|---|---|---|---|
| Control | 25 | 0/10 | 1/279 (Range 173 - 449) |
| | 50 | 2/10 | |
| | 100 | 3/10 | |
| | 200 | 5/10 | |
| | 400 | 8/10 | |

J

| NSGW41 | LT-HSC dose | 1° transplantation engrafted/injected | Stem cell frequency |
|---|---|---|---|
| Control | 25 | 0/10 | 1/265 (Range 159 - 443) |
| | 50 | 2/10 | |
| | 100 | 4/9 | |
| | 200 | 4/8 | |
| | 400 | 6/8 | |

**Fig. S3 CRISPR/Cas9-editing and single cell analysis upon GATA1s and STAG2 knock-out.**
**(A)** Percentage of single cells that grew into a colony in the *in vitro* differentiation/proliferation assay for N-FL (n = 2 experiments). **(B)** Percentage of single cells as described in (A) for T21-FL (§p for combined T21-FL LT-HSCs versus N-FL LT-HSCs, n = 2-3 experiments). **(C)** Percentage of CRISPR/Cas9 efficiency in single cell derived *in vitro* colonies for N-FL (n = 2 experiments). **(D)** Percentage of CRISPR/-Cas9 efficiency as described in (C) for T21-FL (n = 2-3 experiments). **(E)** Cell cycle analysis of N-FL and T21-FL HSPC subpopulations based on EdU incorporation and propidium iodide staining (§ and §§§ indicate significance in relation to the respective N-FL condition, n = 3 replicates per condition). **(F)** Percent cells of quiescent G0 and G1 cells from (E) based on Ki67 staining (n = 3 replicates per condition). **(G)** Overall percentage of megakaryocytic, CD71+, erythroid and myeloid containing colonies from the N-FL *in vitro* assay described in Fig. 1G. Individual percentages are relative to control and are shaded based on a 2-color scale, normalized by each column, and this also applies to (H) **(H)** Overall percentage of colonies from the T21-FL *in vitro* assay described in Fig. 1H. **(I)** Stem cell frequency based on primary xenotransplantations of N-FL control LT-HSC transplanted NSG mice at defined doses for 20 weeks. Limiting dilution analysis was used to assess normal stem cell frequency (n = 10 mice per condition). **(J)** Stem cell frequency as described in (I) for NSGW41 mice (n = 8-10 mice per condition). Unpaired t test: */§p < 0.05; **p < 0.01; ***/§§§p < 0.001; error bars represent standard deviation.

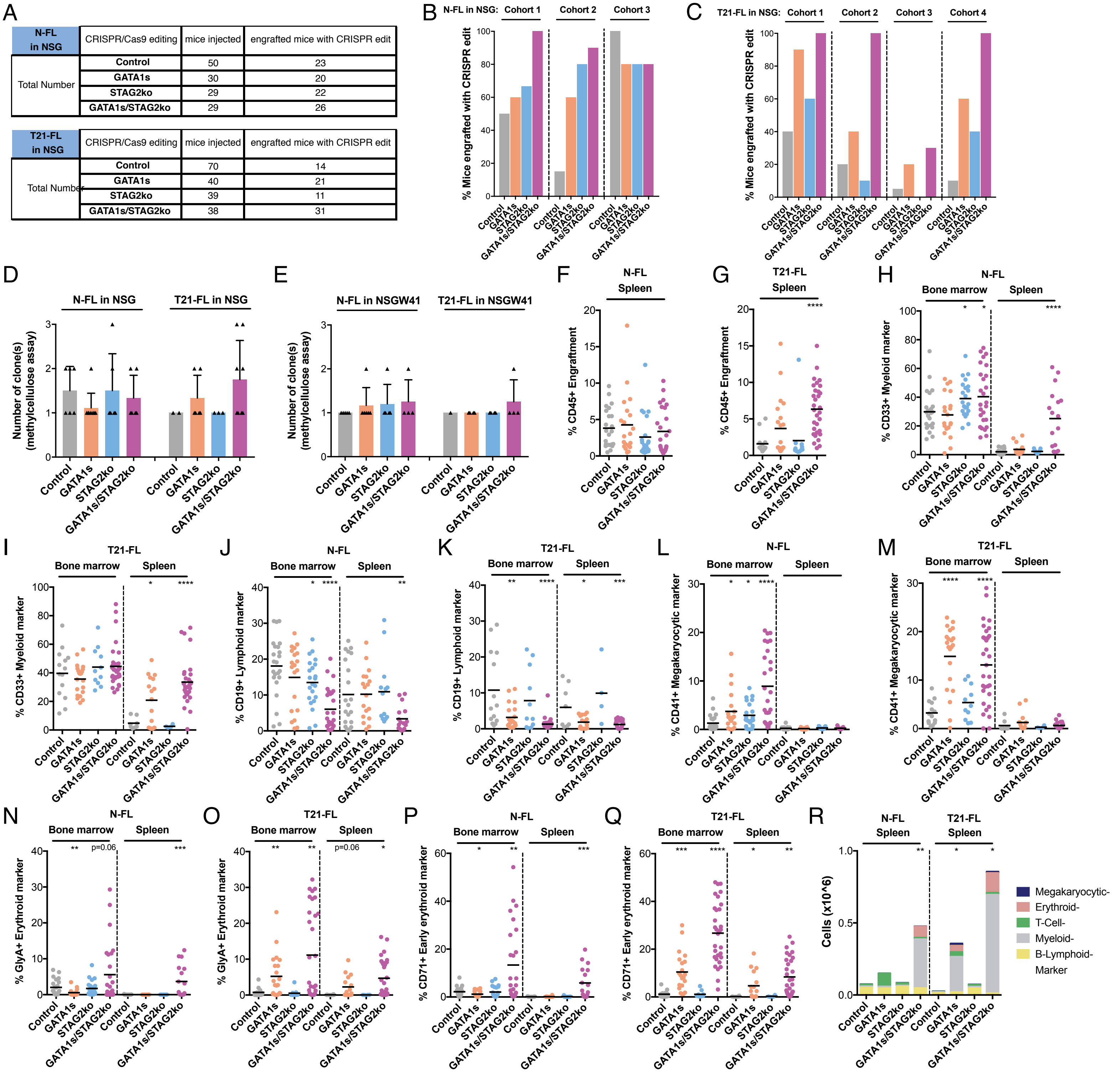


**Fig. S4 Lineage differentiation analysis upon GATA1s and STAG2 knock-out.**

**(A)** Number of transplanted mice in total for NSG mice and the number of mice used for analysis after validation of CRISPR/Cas9 edits (n = 3 cohorts for N-FL and n = 4 cohorts for T21-FL). **(B)** Percentage of NSG mice engrafted in each of the 3 cohorts using CRISPR/Cas9-edited N-FL LT-HSCs. **(C)** Percentage of NSG mice engrafted in each of the 4 cohorts using CRISPR/Cas9-edited T21-FL LT-HSCs (*p for T21-FL GATA1s/STAG2ko versus control and **p for combined N-FL and T21-FL GATA1s/STAG2ko versus control). **(D)** Number of clones in primary xenografts as detected by Sanger sequencing of individual methylcellulose colonies from grafts of NSG mice (n = 2-9 mice per condition, 12 colonies were screened per mouse, total numbers of colonies evaluated for N-FL and T21-FL were 834 and 856, respectively). **(E)** Number of clones in primary xenografts as described in (D) for NSGW41 mice (n = 1-6 mice per condition, total numbers of colonies evaluated for N-FL and T21-FL were 602 and 679, respectively). **(F)** Engraftment levels of N-FL LT-HSC grafts in spleen of NSG mice from Fig. 2A. **(G)** Engraftment levels of T21-FL LT-HSC grafts in spleen of NSG mice from Fig. 2B (n = 4 cohorts). **(H)** Percentage of CD33+CD45+ myeloid cells in N-FL LT-HSC grafts in NSG mice (only grafts with >1% CD45+ cells in spleen are depicted, n = 3 cohorts). **(I)** Percentage of myeloid cells as described in (H) for T21-FL (n = 4 cohorts). **(J)** Percentage of CD19+CD45+ lymphoid cells in N-FL grafts in NSG mice. **(K)** Percentage of lymphoid cells as described in (J) for T21-FL. **(L)** Percentage of CD41+CD45– megakaryocytic cells in N-FL grafts. **(M)** Percentage of CD41+CD45– megakaryocytic cells as described in (L) for T21-FL. **(N)** Percentage of GlyA+ erythroid cells in N-FL grafts in NSG mice. **(O)** Percentage of GlyA+ cells as described in (N) for T21-FL. **(P)** Percentage of CD71+ early erythroid cells in N-FL grafts in NSG mice. **(Q)** Percentage of CD71+ cells as described in (P) for T21-FL. **(R)** Absolute cell numbers in spleen of grafts in NSG mice. Unpaired t test: *p < 0.05; **p < 0.01; ***p < 0.001; ****p < 0.001; error bars represent standard deviation.

Wagenblast et al., Supplementary Figure 5

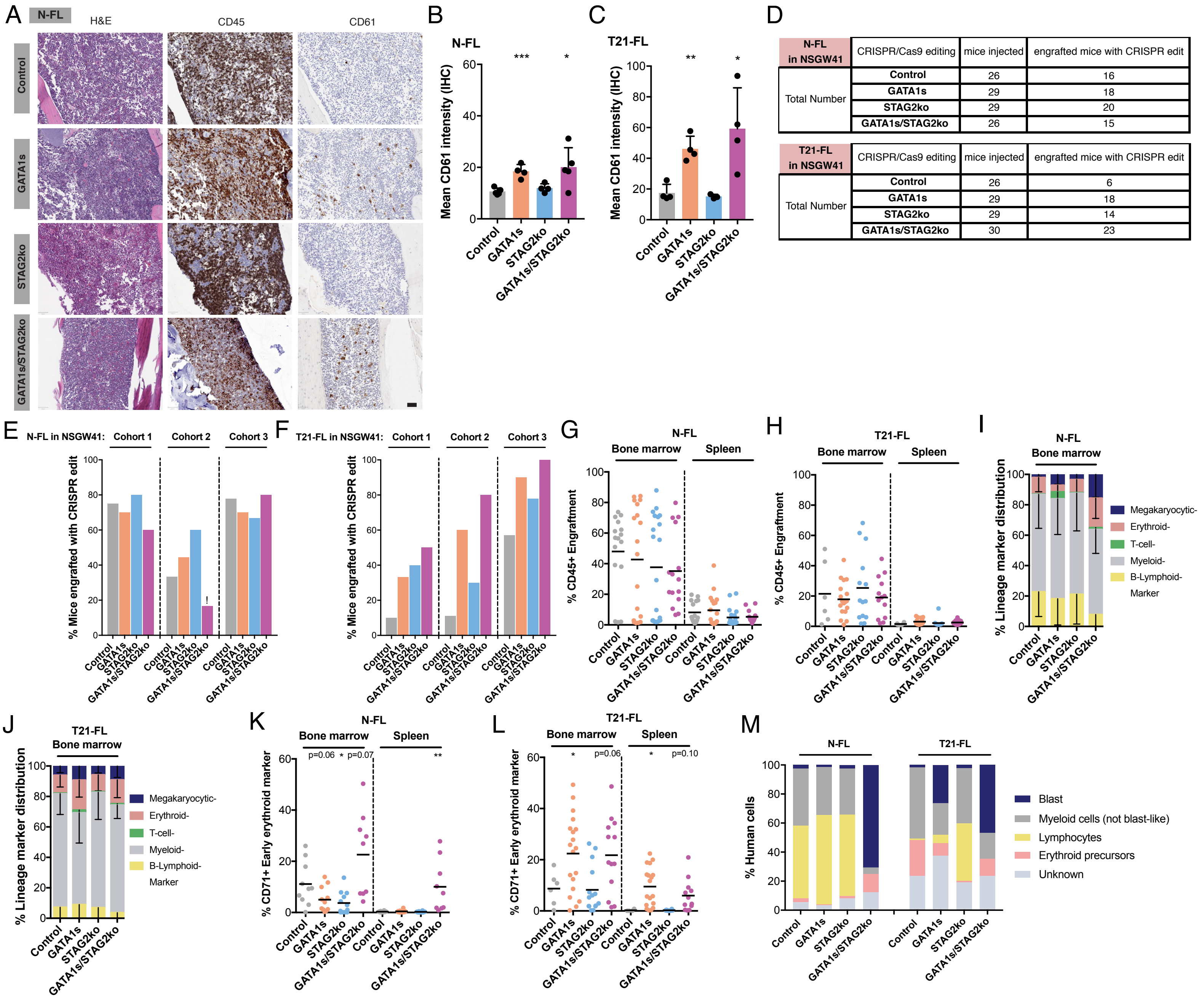


**Fig. S5 Lineage differentiation analysis and characterization of leukemic potential upon GATA1s and STAG2 knock-out.**
**(A)** H&E and IHC stainings for human CD45 and human megakaryocytic marker CD61 in humeri of N-FL grafts in NSG mice (scale 50μ m). **(B)** Quantification of percentage of human CD61 from (H) (n = 4-5 humeri per condition). **(C)** Quantification of percentage of human CD61 from Fig. 2E (n = 3-4 humeri per condition). **(D)** Number of transplanted mice in total for NSGW41 mice and the number of mice used for analysis after validation of CRISPR/Cas9 edits (n = 3 cohorts for N-FL and T21-FL). **(E)** Percentage of NSGW41 mice engrafted in each of the 3 cohorts using CRISPR/Cas9-edited N-FL LT-HSCs. The exclamation sign indicates 4 dead mice that could not be included in the analysis. **(F)** Percentage of NSGW41 mice engrafted in each of the 3 cohorts using CRISPR/Cas9-edited T21-FL LT-HSCs. **(G)** Engraftment levels of N-FL LT-HSC grafts in NSGW41 mice (n = 3 cohorts). **(H)** Engraftment levels as described in (G) for T21-FL (n = 3 cohorts). **(I)** Lineage marker distribution based on cell surface markers in N-FL grafts in NSGW41 mice. **(J)** Lineage marker distribution as described in (I) for T21-FL. **(K)** Percentage of CD71+ early erythroid cells in N-FL grafts in NSGW41 mice. **(L)** Percentage of CD71+ cells as described in (L) for T21-FL. **(M)** Quantification of morphological analysis of human cytospinned cells in N-FL grafts in NSGW41 mice (n = 400 cells per condition). Unpaired t test: *p < 0.05; **p < 0.01; error bars represent standard deviation.

Wagenblast et al., Supplementary Figure 6

A

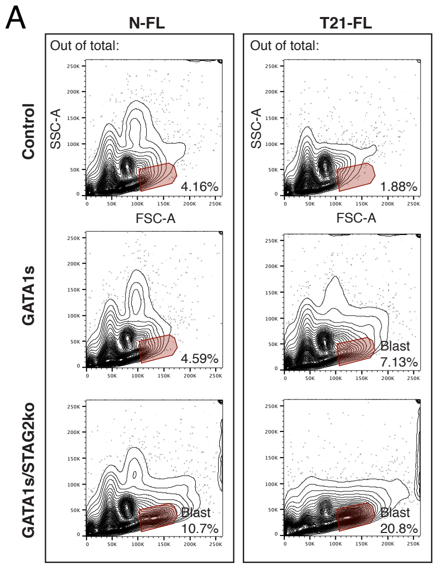


B

| Markers | out of CD45+: | N-FL in NSGW41 Control | GATA1s | GATA1s/STAG2ko | Pre- or Leukemia Patient |
|---|---|---|---|---|---|
| Stem- | % CD34+ | 13.8 | 9.5 | 15.0 | –/+ |
| | % CD117+ | 5.5 | 4.4 | 77.2 | + |
| Meg- | % CD41+ | 2.1 | 4.8 | 15.0 | + |
| | % CD42b+ | 2.6 | 2.3 | 11.2 | + |
| | % CD61+ | 1.1 | 2.6 | 6.9 | + |
| Erythroid- | % CD36+ | 7.7 | 8.3 | 44.5 | + |
| | % CD71+ | 5.6 | 2.7 | 76.6 | + |
| | % GlyA+ | 5.8 | 2.2 | 35.2 | – |
| Lymphoid- | % CD4+ | 19.5 | 15.9 | 83.6 | + |
| | % CD7+ | 10.1 | 7.8 | 73.7 | + |
| | % HLA-Dr+ | 79.3 | 75.4 | 37.0 | –/+ |
| | % CD56+ | 2.1 | 1.2 | 7.1 | –/+ |
| Myeloid- | % CD33+ | 22.4 | 19.2 | 87.3 | + |
| | % CD11b+ | 6.8 | 5.34 | 7.6 | + |
| | % CD13+ | 2.6 | 2.1 | 1.8 | –/+ |
| | % CD14+ | 5.5 | 4.8 | 3.3 | – |

C

| Markers | out of CD45+: | T21-FL in NSGW41 Control | GATA1s | GATA1s/STAG2ko | Pre- or Leukemia Patient |
|---|---|---|---|---|---|
| Stem- | % CD34+ | 12.0 | 41.3 | 38.4 | –/+ |
| | % CD117+ | 18.4 | 45.0 | 77.3 | + |
| Meg- | %CD41 | 10.3 | 43.7 | 10.7 | + |
| | %CD42b | 3.0 | 49.4 | 33.6 | + |
| | %CD61 | 2.1 | 22.7 | 2.8 | + |
| Erythroid- | %CD36 | 22.7 | 25.1 | 14.6 | + |
| | %CD71 | 19.0 | 65.6 | 83.4 | + |
| | %GlyA | 18.9 | 44.6 | 67.5 | – |
| Lymphoid- | %CD4 | 34.1 | 70.1 | 73.2 | + |
| | %CD7 | 20.4 | 70.3 | 83.3 | + |
| | %HLA-Dr | 69.6 | 7.6 | 0.5 | –/+ |
| | %CD56 | 44.2 | 25.7 | 20.4 | –/+ |
| Myeloid- | %CD33 | 51.9 | 78.5 | 80.5 | + |
| | %CD11b | 13.2 | 5.74 | 6.9 | + |
| | %CD13 | 3.9 | 3.0 | 1.1 | –/+ |
| | %CD14 | 1.1 | 0.7 | 0.5 | – |

D

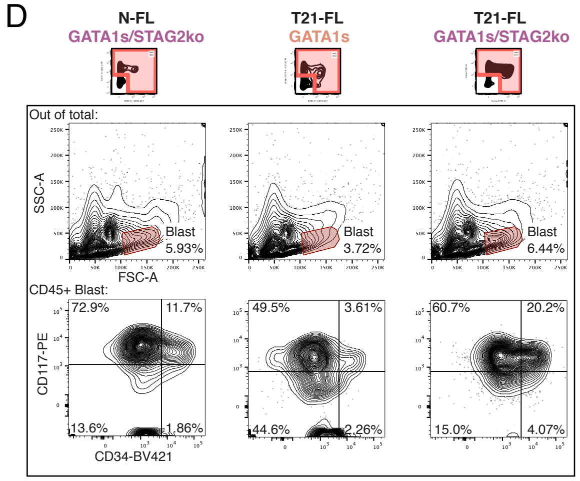


E

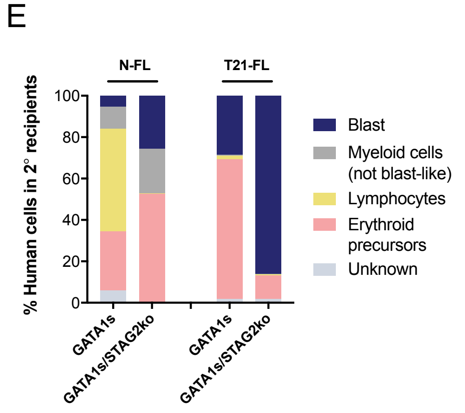


F

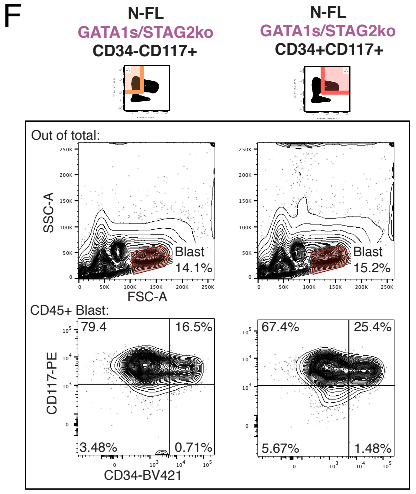


G

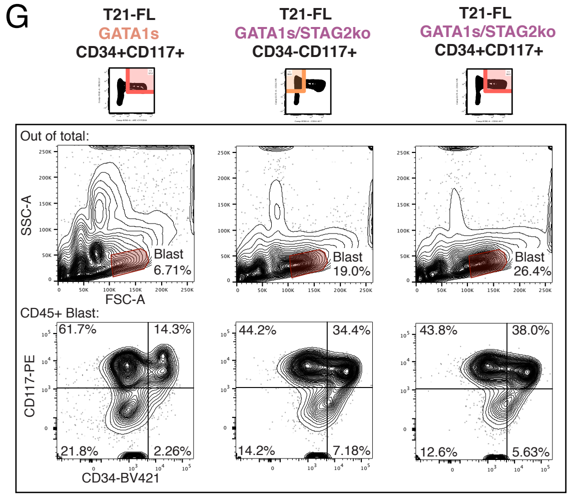


H

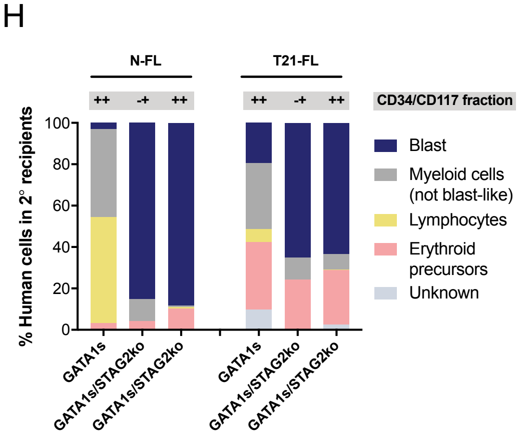


**Fig. S6 Leukemic potential and secondary engraftment upon GATA1s and STAG2 knock-out**
**(A)** Flow cytometry plots depicting the blast population out of total cells in primary xenografts of NSG mice. **(B)** Percent expression of cell surface markers within the CD45+ blast population in primary xenografts of N-FL grafts in NSGW41 mice. For reference, general expression status of each cell surface marker in patient blasts' of Down syndrome preleukemia and leukemia is indicated to the right, although expression is variable from patient to patient. Individual percentages are shaded based on a 2-color scale, normalized by the whole table and this also applies to (C) (data from pooled samples of multiple xenografts). **(C)** Percent expression of cell surface markers as described in (B) for T21-FL (data from pooled samples of multiple xenografts). **(D)** Flow cytometry plots of blast populations out of total cells in secondary xenografts of NSGW41 mice. **(E)** Quantification of morphological analysis of human cytospinned cells in secondary xenografts of N-FL and T21-FL grafts in NSGW41 mice (n = 400 cells per condition). **(F)** Flow cytometry plots of blast populations out of total cells in secondary NSG xenografts of N-FL. **(G)** Flow cytometry plots of blast populations as described in (F) for T21-FL. **(H)** Quantification of morphological analysis of human cytospinned cells in secondary xenografts of N-FL and T21-FL grafts in NSG mice (n = 400 cells per condition).

Wagenblast et al., Supplementary Figure 7

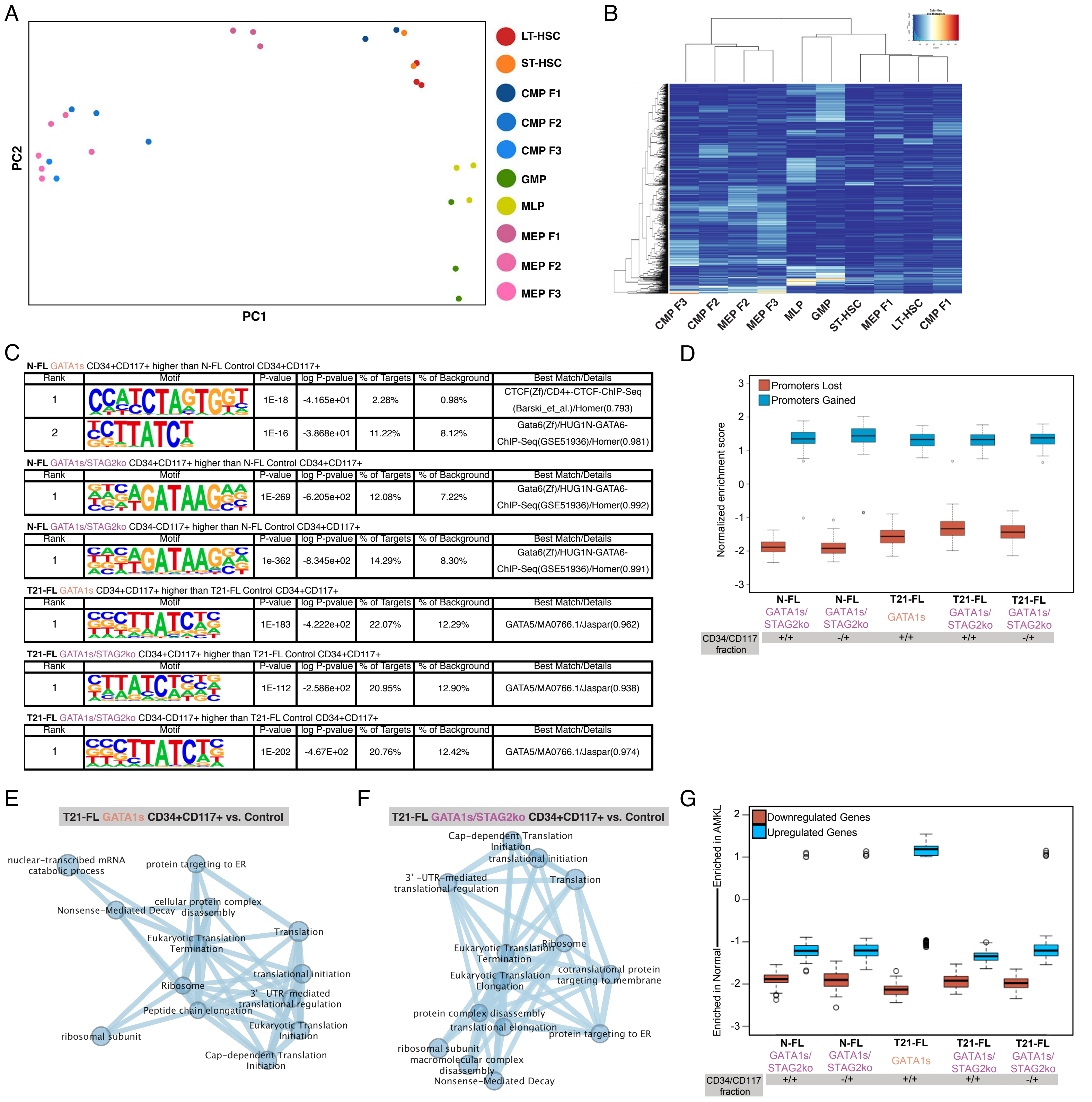


**Fig. S7 Transcriptional and epigenetic profiling of GATA1s and STAG2 knock-out grafts**
**(A)** Principal component analysis of ATACseq profiles of individually sorted N-FL HSPC subpopulations (n = 3 fetal livers). **(B)** Heatmap showing the CIBERSORTx signature matrix, which was generated from (A). **(C)** Motif enrichment analysis of open chromatin accessible promoters sites assessed through ATACseq. Algorithm can not differentiate between binding motifs of GATA1 to 6 (see table S3 for full list of motif enrichments, n = 3 replicates per condition). **(D)** Enrichment in gene expression assessed through RNAseq of genes with open chromatin accessible promoters gained and lost in N-FL and T21-FL GATA1s and GATA1s/STAG2ko LT-HSC primary xenografts (n = 3 replicates per condition). **(E)** Pathway enrichment analysis of T21-FL GATA1s-edited CD34+CD117+ fraction from primary xenografts compared to control (see table S5 for all pathways and other comparisons, n = 3 replicates per condition). **(F)** Pathway enrichment analysis of T21-FL GATA1s/STAG2-edited CD34+CD117+ fraction from primary xenografts compared to control. **(G)** Enrichment of down- and up-regulated genes of N-FL and T21-FL GATA1s and GATA1s/STAG2ko LT-HSC primary xenografts in gene expression profiles of patient derived samples of Down syndrome leukemia blasts and human samples of fetal liver derived CD34+CD38– enriched stem cells (n = 3 replicates per condition).

A

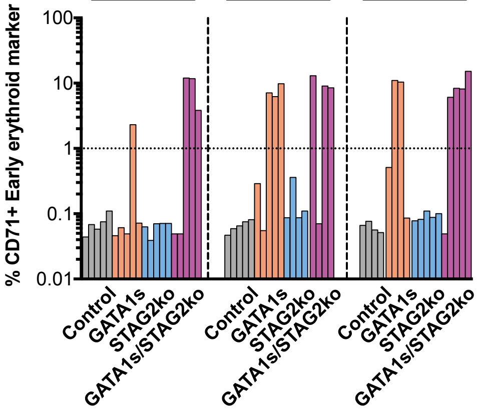


B

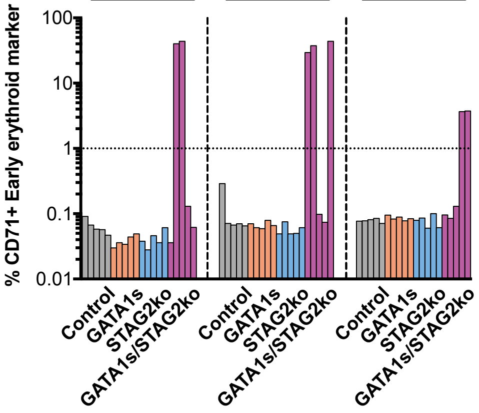


C

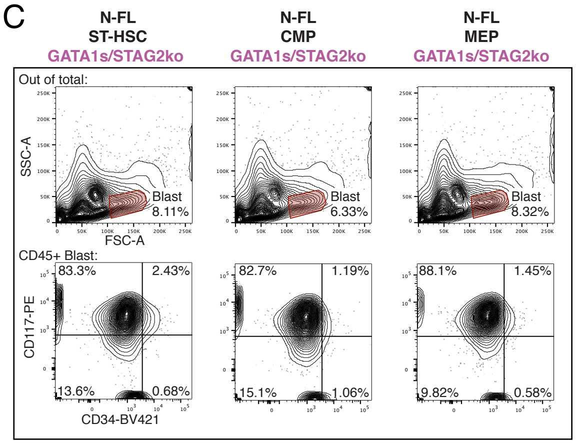


D

| Markers | out of CD45+: | N-FL ST-HSC GATA1s/STAG2ko | N-FL CMP GATA1s/STAG2ko | N-FL MEP GATA1s/STAG2ko | Pre- or Leukemia Patient |
|---|---|---|---|---|---|
| Stem- | % CD34+ | 3.1 | 2.3 | 2.0 | –/+ |
| | % CD117+ | 85.7 | 83.9 | 89.6 | + |
| Meg- | % CD41+ | 23.9 | 32.4 | 24.4 | + |
| | % CD42b+ | 5.6 | 7.1 | 4.7 | + |
| | % CD61+ | 3.6 | 5.5 | 2.1 | + |
| Erythroid- | % CD36+ | 40.5 | 43.9 | 30.5 | + |
| | % CD71+ | 83.7 | 80.3 | 79.8 | + |
| | % GlyA+ | 67.5 | 47.0 | 44.2 | – |
| Lymphoid- | % CD4+ | 89.4 | 88.9 | 87.7 | + |
| | % CD7+ | 76.4 | 74.9 | 71.1 | + |
| | % HLA-Dr+ | 13.8 | 9.2 | 9.9 | –/+ |
| | % CD56+ | 1.2 | 1.4 | 1.3 | –/+ |
| Myeloid- | % CD33+ | 89.9 | 88.2 | 92.8 | + |
| | % CD11b+ | 10.8 | 16.0 | 12.5 | + |
| | % CD13+ | 0.9 | 2.0 | 0.9 | –/+ |
| | % CD14+ | 0.3 | 0.6 | 0.3 | – |

E

| Markers | out of CD45+: | T21-FL ST-HSC GATA1s/STAG2ko | T21-FL CMP GATA1s/STAG2ko | T21-FL MEP GATA1s/STAG2ko | Pre- or Leukemia Patient |
|---|---|---|---|---|---|
| Stem- | % CD34+ | 33.5 | 35.1 | 37.0 | –/+ |
| | % CD117+ | 84.1 | 80.2 | 70.2 | + |
| Meg- | % CD41+ | 2.0 | 3.5 | 8.1 | + |
| | % CD42b+ | 28.6 | 23.3 | 19.9 | + |
| | % CD61+ | 0.1 | 0.1 | 0.5 | + |
| Erythroid- | % CD36+ | 29.5 | 30.2 | 19.9 | + |
| | % CD71+ | 65.9 | 65.6 | 35.5 | + |
| | % GlyA+ | 43.4 | 44.9 | 33.7 | – |
| Lymphoid- | % CD4+ | 78.7 | 75.8 | 79.2 | + |
| | % CD7+ | 95.5 | 93.3 | 95.7 | + |
| | % HLA-Dr+ | 8.4 | 9.3 | 11.5 | –/+ |
| | % CD56+ | 24.7 | 25.7 | 23.2 | –/+ |
| Myeloid- | % CD33+ | 94.8 | 91.5 | 94.2 | + |
| | % CD11b+ | 0.2 | 0.4 | 1.1 | + |
| | % CD13+ | 0.5 | 0.6 | 0.9 | –/+ |
| | % CD14+ | 0.6 | 1.8 | 0.3 | – |

F

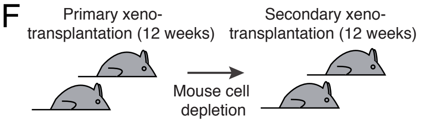


| N-FL | Dose | 2° transplantation engrafted/injected |
|---|---|---|
| ST-HSC GATA1s/STAG2ko | 200,000 | 5/5 |
| CMP GATA1s/STAG2ko | 200,000 | 2/5 |
| MEP GATA1s/STAG2ko | 200,000 | 5/5 |

| T21-FL | Dose | 2° transplantation engrafted/injected |
|---|---|---|
| ST-HSC GATA1s/STAG2ko | 300,000 | 5/5 |
| CMP GATA1s/STAG2ko | 300,000 | 2/2 |
| MEP GATA1s/STAG2ko | 300,000 | 3/3 |

G

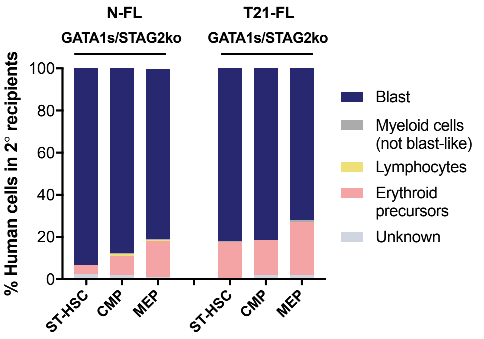


H

| Markers | out of CD45+: | T21-FL GATA1s + STAG2 | T21-FL GATA1s + RAD21 | T21-FL GATA1s + NIPBL | T21-FL GATA1s + SMC1A | T21-FL GATA1s + SMC3 | T21-FL GATA1s + KANSL1 | Pre- or Leukemia Patient |
|---|---|---|---|---|---|---|---|---|
| Stem- | % CD34+ | 18.4 | 8.6 | 4.4 | 6.2 | 1.3 | 4.9 | –/+ |
| | % CD117+ | 70.3 | 58.8 | 37.2 | 78.5 | 47.7 | 35.2 | + |
| Meg- | % CD41 | 23.4 | 28.9 | 20.6 | 51.3 | 32.0 | 42.8 | + |
| | % CD42b | 27.4 | 12.4 | 5.4 | 19.9 | 4.6 | 4.4 | + |
| | % CD61 | 0.0 | 0.2 | 0.2 | 0.3 | 0.1 | 0.5 | + |
| Erythroid- | % CD36 | 26.1 | 22.3 | 12.4 | 23.8 | 24.5 | 15.4 | + |
| | % CD71 | 90.8 | 84.2 | 87.7 | 89.0 | 97.0 | 87.0 | + |
| | % GlyA | 61.4 | 66.7 | 89.6 | 56.6 | 93.0 | 87.0 | – |
| Lymphoid- | % CD4 | 80.8 | 83.1 | 73.1 | 70.4 | 90.1 | 69.4 | + |
| | % CD7 | 80.1 | 80.8 | 60.3 | 88.5 | 68.7 | 35.3 | + |
| | % HLA-Dr | 0.4 | 3.4 | 8.3 | 2.4 | 2.0 | 17.6 | –/+ |
| | % CD56 | 32.8 | 20.0 | 19.6 | 14.5 | 11.3 | 23.8 | –/+ |
| Myeloid- | % CD33 | 85.4 | 85.3 | 81.2 | 87.7 | 89.1 | 85.0 | + |
| | % CD11b | 6.8 | 4.3 | 4.4 | 3.4 | 0.3 | 0.9 | + |
| | % CD13 | 0.3 | 0.7 | 0.7 | 0.6 | 0.2 | 2.1 | –/+ |
| | % CD14 | 2.0 | 1.1 | 1.0 | 1.1 | 0.8 | 1.6 | – |

**Fig. S8 Leukemic potential of progenitor cells upon GATA1s and STAG2 knock-out**

**(A)** Percentage of CD71+ early erythroid cells in N-FL ST-HSC, CMP and MEP grafts in NSGW41 mice (n = 4-5 mice per condition). **(B)** Percentage of CD71+ cells as described in (A) for T21-FL (n = 5 mice per condition). **(C)** Flow cytometry plots of blast populations out of total cells in primary NSGW41 xenografts of N-FL GATA1s/STAG2ko ST-HSC, CMP and MEP grafts. **(D)** Percent expression of cell surface markers within the CD45+ blast population in primary NSGW41 xenografts of GATA1s/STAG2ko-edited N-FL progenitors. Individual percentages are shaded based on a 2-color scale, normalized by the whole table and this also applies to (E) and (H) (data from pooled samples of multiple xenografts). **(E)** Percent expression of cell surface markers as described in (D) for T21-FL (data from pooled samples of multiple xenografts). **(F)** Secondary xenotransplantations of N-FL and T21-FL GATA1s/STAG2ko ST-HSC, CMP and MEP grafts in NSGW41 mice for 12 weeks (>0.1% CD45 in bone marrow, n = 2-5 mice per condition). **(G)** Quantification of morphological analysis of human cytospinned cells in secondary NSGW41 xenografts of GATA1s/STAG2ko N-FL and T21-FL progenitors (n = 400 cells per condition). **(H)** Percent expression of cell surface markers within the CD45+ blast population out of T21-FL CMPs and MEPs edited with GATA1s in combination with other candidate gene knock-outs and transplanted into NSGW41 mice (data from pooled samples of multiple xenografts).

Wagenblast et al., Supplementary Figure 9

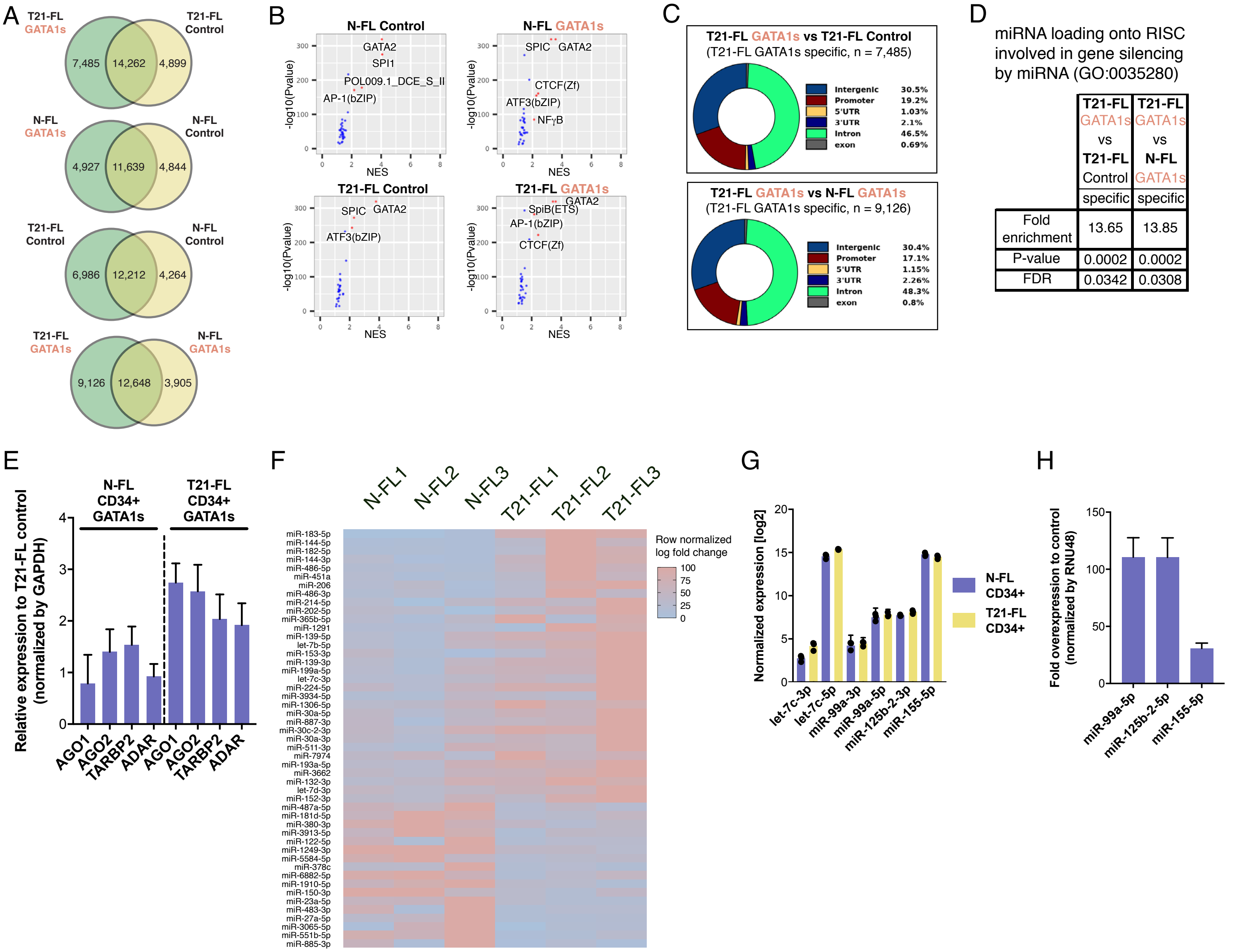


**Fig. S9 Analysis of GATA1 binding occupancy and miRNA profiling.**
**(A)** Venn diagrams of GATA1 binding sites as determined by Cut&Run assays in N-FL and T21-FL control and GATA1s CD34+ enriched HSPCs (n = 1 fetal liver per condition). **(B)** Motif enrichment analysis of GATA1 binding sites in N-FL and T21-FL control and GATA1s CD34+ cells. Algorithm can not differentiate between binding motifs of GATA1 to 6. **(C)** Distribution of GATA1 binding sites in the genome in N-FL and T21-FL control and GATA1s CD34+ cells. **(D)** Enrichment analysis of the GO term "miRNA loading onto RISC involved in gene silencing by miRNA" in GATA1 binding sites that are specific to T21-FL GATA1s CD34+ cells (see table S7 for full list of GO terms). **(E)** Relative expression of AGO1, AGO2, TARBP2 and ADAR in T21-FL GATA1s CD34+ cells compared to N-FL control as measured by RT-qPCR and normalized by GAPDH (n = 1 fetal liver with three technical replicates per condition). **(F)** Differentially expressed miRNAs between N-FL and T21-FL CD34+ enriched HSPCs. Values are row normalized from raw count (n = 3 replicates per condition). **(G)** Normalized expression of all detectable chromosome 21 miRNAs in N-FL and T21-FL CD34+ cells from the same dataset as (F). **(H)** Relative expression of chromosome 21 miRNAs upon lentiviral overexpression of miR-99a, miR-125b-2 and miR-155 in N-FL LT-HSCs as measured by RT-qPCR (n = 1 fetal liver with three technical replicates per condition). Error bars represent standard deviation.

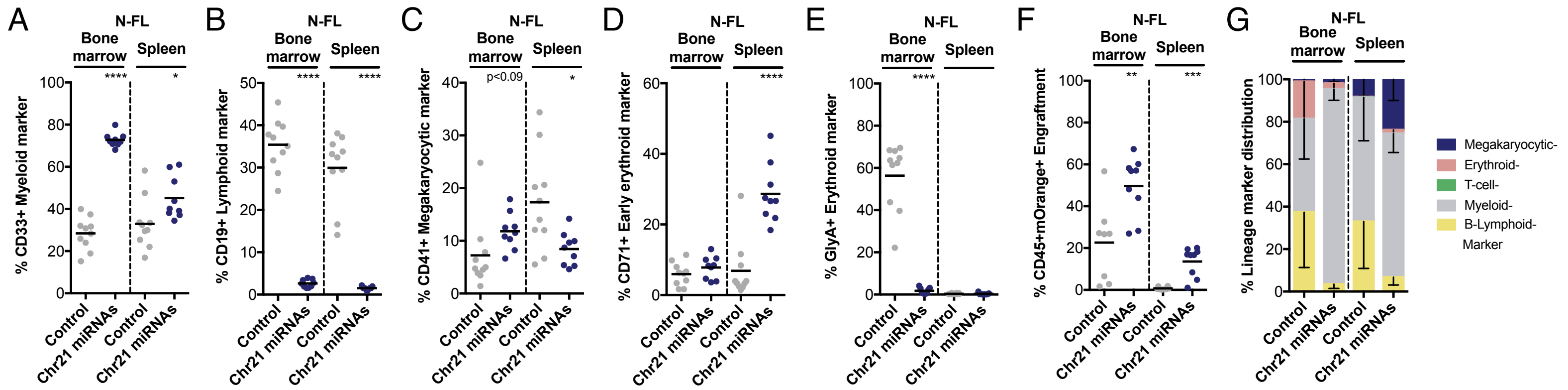


H

| Markers | out of CD45+: | N-FL Control | N-FL Chr21 miRNAs | Pre- or Leukemia Patient |
|---|---|---|---|---|
| Stem- | % CD34+ | 16.4 | 2.8 | –/+ |
| | % CD117+ | 12.0 | 12.6 | + |
| Meg- | % CD41+ | 5.5 | 7.2 | + |
| | % CD42b+ | 3.4 | 41.4 | + |
| | % CD61+ | 2.2 | 21.4 | + |
| Erythroid- | % CD36+ | 9.7 | 8.6 | + |
| | % CD71+ | 5.5 | 6.4 | + |
| | % GlyA+ | 5.5 | 5.3 | – |
| Lymphoid- | % CD4+ | 26.1 | 2.3 | + |
| | % CD7+ | 6.4 | 1.2 | + |
| | % HLA-Dr+ | 80.5 | 82.0 | –/+ |
| | % CD56+ | 8.8 | 5.4 | –/+ |
| Myeloid- | % CD33+ | 35.7 | 89.7 | + |
| | % CD11b+ | 5.0 | 75.8 | + |
| | % CD13+ | 7.7 | 13.3 | –/+ |
| | % CD14+ | 5.8 | 8.0 | – |

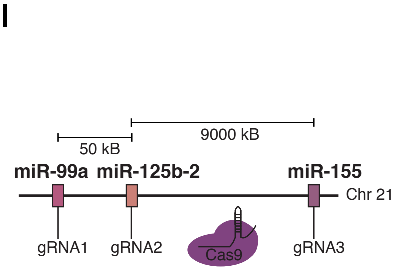


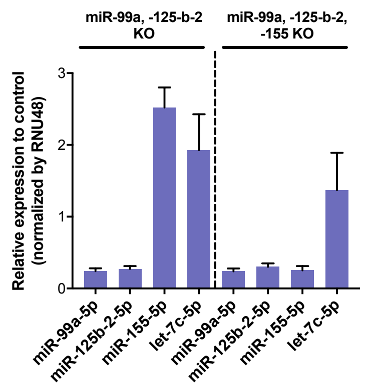


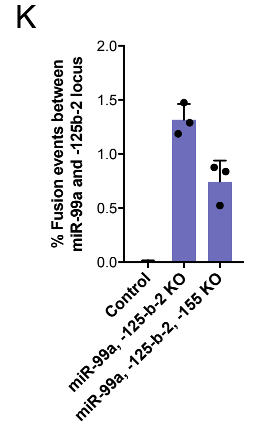


L

| Markers | out of CD45+: | T21-FL GATA1s Vehicle | T21-FL GATA1s Imatinib | T21-FL GATA1s Dasatinib | T21-FL GATA1s Ripretinib | Pre- or Leukemia Patient |
|---|---|---|---|---|---|---|
| Stem- | % CD34+ | 18.2 | 12.1 | 21.2 | 12.6 | –/+ |
| | % CD117+ | 73.5 | 29.7 | 34.2 | 23.2 | + |
| Meg- | % CD41 | 31.6 | 22.7 | 15.2 | 10.5 | + |
| | % CD42b | 20.4 | 10.8 | 12.2 | 7.4 | + |
| | % CD61 | 2.4 | 2.3 | 2.0 | 1.4 | + |
| Erythroid- | % CD36 | 18.0 | 22.6 | 19.4 | 18.3 | + |
| | % CD71 | 62.1 | 24.4 | 23.7 | 16.1 | + |
| | % GlyA | 44.6 | 21.1 | 10.1 | 8.8 | – |
| Lymphoid- | % CD4 | 67.2 | 57.0 | 40.4 | 43.2 | + |
| | % CD7 | 82.5 | 51.8 | 60.9 | 51.7 | + |
| | % HLA-Dr | 12.1 | 54.3 | 59.5 | 66.7 | –/+ |
| | % CD56 | 22.4 | 30.5 | 33.0 | 30.7 | –/+ |
| Myeloid- | % CD33 | 66.9 | 56.2 | 60.0 | 49.8 | + |
| | % CD11b | 3.2 | 7.5 | 3.2 | 6.2 | + |
| | % CD13 | 3.2 | 13.1 | 11.4 | 17.1 | –/+ |
| | % CD14 | 2.4 | 8.7 | 5.4 | 6.1 | – |

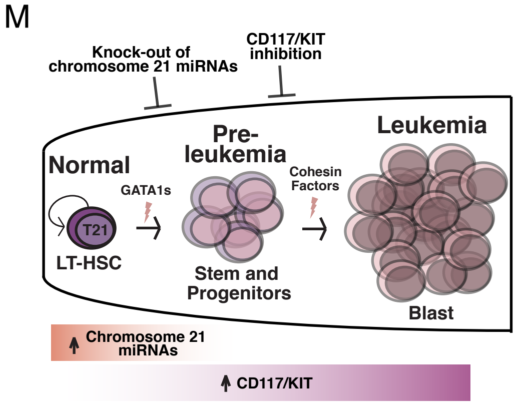


**Fig S10 Role of chromosome 21 miRNAs in preleukemic initiation and leukemic progression and CD117/KIT inhibition.**
**(A)** Percentage of CD33+CD45+ myeloid cells in N-FL transduced control and Chr21 miRNAs LT-HSC transplanted NSG mice (n = 9-10 mice per condition). **(B)** Percentage of CD19+CD45+ lymphoid cells in N-FL grafts in NSG mice (n = 9-10 mice per condition). **(C)** Percentage of CD41+CD45– megakaryocytic cells in N-FL grafts in NSG mice (n = 9-10 mice per condition). **(D)** Percentage of CD71+ early erythroid cells in N-FL grafts in NSG mice (n = 9-10 mice per condition). **(E)** Percentage of GlyA+ erythroid cells in N-FL grafts in NSG mice (n = 9-10 mice per condition). **(F)** Engraftment levels of N-FL transduced control and Chr21 miRNAs LT-HSC transplanted NSGW41 mice (only mice with >1% CD45+ cells in bone marrow were analyzed, n = 8-9 mice per condition). **(G)** Lineage marker distribution based on cell surface markers of engrafted NSGW41 mice from (F). **(H)** Percent expression of cell surface markers within the CD45+ blast population in primary xenografts of N-FL transduced control and Chr21 miRNAs LT-HSC transplanted NSG mice. Individual percentages are shaded based on a 2-color scale, normalized by the whole table and this also applies to (L) (data from pooled samples of multiple xenografts). **(I)** Schematic overview of genomic location of miR-99a, miR-125b-2 and miR-155 on chromosome 21. **(J)** Relative expression of miR-99a, miR-125b-2, miR-155 and let-7c upon knock-out of Chr21 miRNAs in N-FL LT-HSCs as measured by RT-qPCR (n = 1 fetal liver with three technical replicates per condition). **(K)** Percent off-target fusion events between the 5'-end of the miR-99a locus and the 3'-end of the miR-125b-2 locus upon CRISPR/Cas9 editing based on droplet digital PCR in N-FL LT-HSCs (n = 3 replicates per condition). **(L)** Percent expression of cell surface markers within the CD45+ blast population in T21-FL GATA1s LT-HSC grafts treated with control, imatinib, dasatinib and ripretinib (data from pooled samples of multiple xenografts). **(M)** Schematic displaying the cells of origin, dependencies and targetability of Down syndrome leukemogenesis. Unpaired t test: *p < 0.05; **p < 0.01; ***p < 0.001; ****p < 0.001; error bars represent standard deviation.
